# Microbial population dynamics decouple growth response from environmental nutrient concentration

**DOI:** 10.1101/2022.05.04.490627

**Authors:** Justus Wilhelm Fink, Noelle A. Held, Michael Manhart

## Abstract

How the growth rate of a microbial population responds to the environmental availability of chemical nutrients and other resources is a fundamental question in microbiology. Models of this response, such as the widely-used Monod model, are generally characterized by a maximum growth rate and a half-saturation concentration of the resource. What values should we expect for these half-saturation concentrations, and how should they depend on the environmental concentration of the resource? We survey growth response data across a wide range of organisms and resources. We find that the half-saturation concentrations vary across orders of magnitude, even for the same organism and resource. To explain this variation, we develop an evolutionary model to show that demographic fluctuations (genetic drift) can constrain the adaptation of half-saturation concentrations. We find that this effect fundamentally differs depending on the type of population dynamics: populations undergoing periodic bottlenecks of fixed size will adapt their half-saturation concentration in proportion to the environmental resource concentration, but populations undergoing periodic dilutions of fixed size will evolve half-saturation concentrations that are largely decoupled from the environmental concentration. Our model not only provides testable predictions for laboratory evolution experiments, but it also reveals how an evolved half-saturation concentration may not reflect the organism’s environment. In particular, this explains how organisms in resource-rich environments can still evolve fast growth at low resource concentrations. Altogether our results demonstrate the critical role of population dynamics in shaping fundamental ecological traits.

## INTRODUCTION

Microbial populations rely on a wide range of resources, including chemical nutrients such as sugars, minerals, and metals, as well as space, light, and prey [1]. These resources vary in abundance across time and environments, which typically elicits differences in growth rates [2–4]. A significant literature discusses how natural populations can be classified as oligotrophs or copiotrophs [4–6], that differ, among other things, in their growth rate response to resource concentration. The most widely-used quantitative model of the relationship between growth rate and resource concentration is attributed to Jacques Monod [7]. In the Monod model, growth rate increases linearly with resource concentration at low concentrations, and then saturates at high concentrations, reaching half its maximum value at some intermediate concentration of resources. This half-saturation concentration of the growth response, also known as the Monod constant, therefore plays a key role in determining the ability of the population to grow on scarce resources. This suggests that lower resource concentrations in the environment may drive populations to evolve commensurately lower half-saturation concentrations [8, 9], one of the main predictions of resource-ratio theory [10–12]. Quantitative models and data for the dependence of growth rate on resource concentration are important both for predicting the behavior of a population under different environmental conditions [13–15], as well as for inferring the natural environmental niche from evolved traits of the population. This inverse approach has been used, for example, to infer separate niches for ammonia-oxidizing archaea and bacteria in the global nitrogen cycle based on kinetic parameters for resource consumption [16–19].

Even though these concepts have been central elements of microbiology and ecology for decades, there is limited experimental evidence that directly demonstrates the evolution of growth rate response to resources. Continuous culture for 200–300 generations led to improved growth rate at low glucose concentrations for *Escherichia coli* [20, 21] and *Saccharomyces cerevisiae* [22], but these changes were not clearly attributable to genetic (rather than physiological) adaptation. The Long-Term Evolution Experiment (LTEE) of *E. coli* found that the half-saturation concentration for glucose actually increased over the first 2000 generations, although the maximum growth rate at much higher glucose concentrations significantly increased as well [23]. More recently, Bernhardt et al. [12] observed adaptation in the half-saturation concentration for phosphorus of *Chlamydomonas reinhardtii* when limited for phosphorus, but they did not obtain consistent outcomes for nitrogen and light. Perhaps the most explicit evidence so far is from Hart et al. [24], who found that a synthetic auxotroph strain of *S. cerevisiae* significantly reduced its half-saturation concentration for lysine through genetic adaptations.

While laboratory experiments can test the basic principle, mathematical models are better suited to exploring the wide range of environments necessary to establish the link between environment and evolved traits. Previous modeling studies on this topic have focused on how tradeoffs in growth rate at low versus high resource concentrations define an optimum strategy for a single strain [13] or can facilitate coexistence of multiple strains or species when resource concentrations fluctuate [25, 26]. More recent work has shown how this coexistence can spontaneously evolve if such tradeoffs constrain the effects of mutations [27, 28]. However, the evidence for these tradeoffs, especially on spontaneous mutations, is limited [27–31]. Thus their importance for explaining the evolved variation in growth rate response, especially the half-saturation concentration, is unclear.

Here we address this problem using both empirical and modeling approaches. We first perform a survey of data for the growth rate response to resource concentration across a wide range of organisms and resources. We find that the measured half-saturation concentrations vary over orders of magnitude, even within some single species on the same resource, such as *E. coli* strains on glucose. We also find no evidence for tradeoffs between growth rate at low versus high resource concentrations. To better understand the potential causes of this variation, we model evolution for populations with a single limiting resource under feast-and-famine conditions (batch dynamics with fixed biomass or fixed dilution factor) and steady-state growth (chemostat dynamics). We show how demographic fluctuations, known as genetic drift, inhibit selection on lower half-saturation concentration, which leads to a general relationship between the evolved half-saturation concentration, environmental resource concentration, and the effective population size. Using this result, we determine that populations with fixed-bottleneck batch dynamics will evolve half-saturation concentrations that are proportional to the environmental resource concentration, but populations with fixed-dilution batch dynamics evolve half-saturation concentrations that are practically independent of the environment. Besides providing a testable theory for laboratory evolution experiments, our results help to explain how species evolving under high concentrations can maintain fast growth at low concentrations and why evolved half-saturation concentrations may not reflect the environment of origin.

## RESULTS

### The Monod model quantifies growth rate response to resource concentration

Consider a population of microbes consuming a resource; we will generally focus on chemical nutrients such as carbon or nitrogen sources, but some aspects of the model apply to other types of resources as well (e.g., prey or light). While microbes consume many different resources simultaneously [32, 33], for simplicity here we assume only a single resource limits growth (Supplementary Information Sec. S1). The best-known dependence of population growth rate *g* on resource concentration *R* is the Monod model [7]:

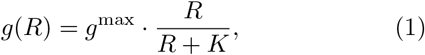

where *g*^max^ is the maximum growth rate — achieved when the resource is unlimited — and *K* is the concentration for the resource at which growth rate is slowed to half its maximum (Fig. 1). Decreasing the half-saturation concentration *K* therefore allows the population to grow faster at lower resource concentrations. The half-saturation concentration *K* is not to be confused with a related but distinct concept of *R*\* from resourceratio theory [10, 12]. Note that the Monod model of Eq. (1) is used to describe both steady state [12] and non-steady-state [25, 28] relationships between growth rate and environmental resource concentration. While there are many alternative models of how growth rate depends on resource concentration (Supplementary Information Sec. S2, Table S1), we focus on the Monod model due to its wider usage and available data.

**FIG. 1.**
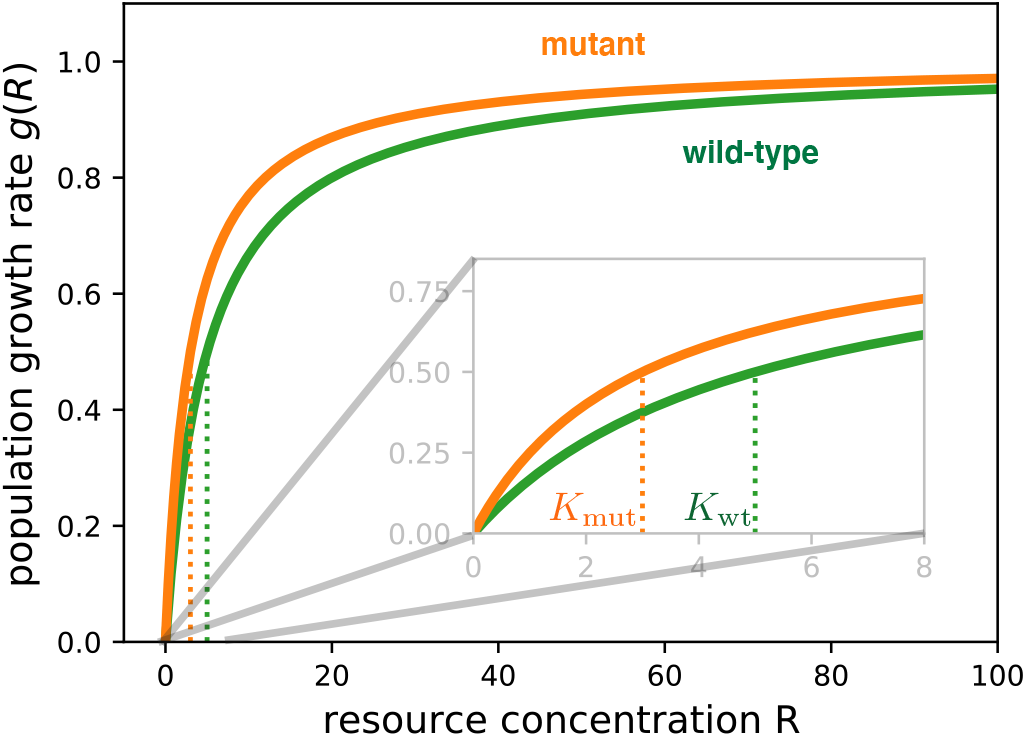
Monod model of growth rate response to resource concentration. The population growth rate *g*(*R*) as a function of the external resource concentration *R* for two hypothetical strains: a wild-type (green) and a derived mutant strain (orange), with equal maximum growth rates (*g*^max^ = 1) but different half-saturation concentrations (*K*_wt_ = 5, *K*_mut_ = 3). The inset shows a magnified view at low concentrations near *K*_wt_ and *K*_mut_ (dotted vertical lines). Note that the growth rates do not fully overlap at the highest concentration shown, but eventually converge to the same value *g*^max^ outside the range of this plot.

The parameter *K* is sometimes labeled as the affinity for the resource [34], but this is potentially misleading as *K* is inversely proportional to ability to grow on the resource. We instead use the term *specific affinity* to refer to the parameter combination *g*^max^/*K*, which measures how much the growth rate increases per unit change in resource concentration, starting from a low concentration [35]. The specific affinity is therefore a common measure for oligotrophic growth ability [9, 16, 19, 34]. Note that both *K* and *g*^max^ are required to fully characterize the growth rate dependence; for example, the specific affinity *g*^max^/*K* alone does not suffice because while it describes the growth rate response at low concentrations, it does not define the range of low concentrations (which is determined separately by *K*). Since we are primarily interested in how these traits evolve in relation to the environmental concentration *R*, we focus primarily on the half-saturation concentration *K* since one can directly compare it to *R*.

One can derive the Monod model of Eq. (1) by modeling biomass growth as a two-step process, in which uptake of the external resource into the cell occurs at a rate proportional to the external concentration *R* [36]. However, the dependence of growth rate on resource concentration expressed by Eq. (1) is surprisingly robust to additional model complexities [37, 38], albeit with the resulting traits *g*^max^ and *K* being emergent properties of whole cells or populations. In particular, the half-saturation concentration *K* is not equivalent to the Michaelis-Menten constant for resource uptake kinetics [37, 39, 40], despite the mathematical similarity between the Michaelis-Menten and Monod models (Eq. 1); this is because the Monod model describes the whole process of producing new biomass, of which uptake is just one step.

### Half-saturation concentrations vary widely across resources and organisms

To explore the diversity of microbial growth responses, we have compiled 247 measurements of half-saturation concentrations *K* from previously-published studies (Methods; Dataset S1; Fig. S1), substantially extending previous surveys [41–44]. Figure 2A shows an overview of this data, sorted by resource. The data includes a wide range of resources, with phosphate, glucose, and nitrate having the largest number of measurements due to their emphasis in marine and laboratory systems. Organisms include prokaryotes and eukaryotes as well as autotrophs and heterotrophs (marked by different symbols in Fig. 2A).

**FIG. 2.**
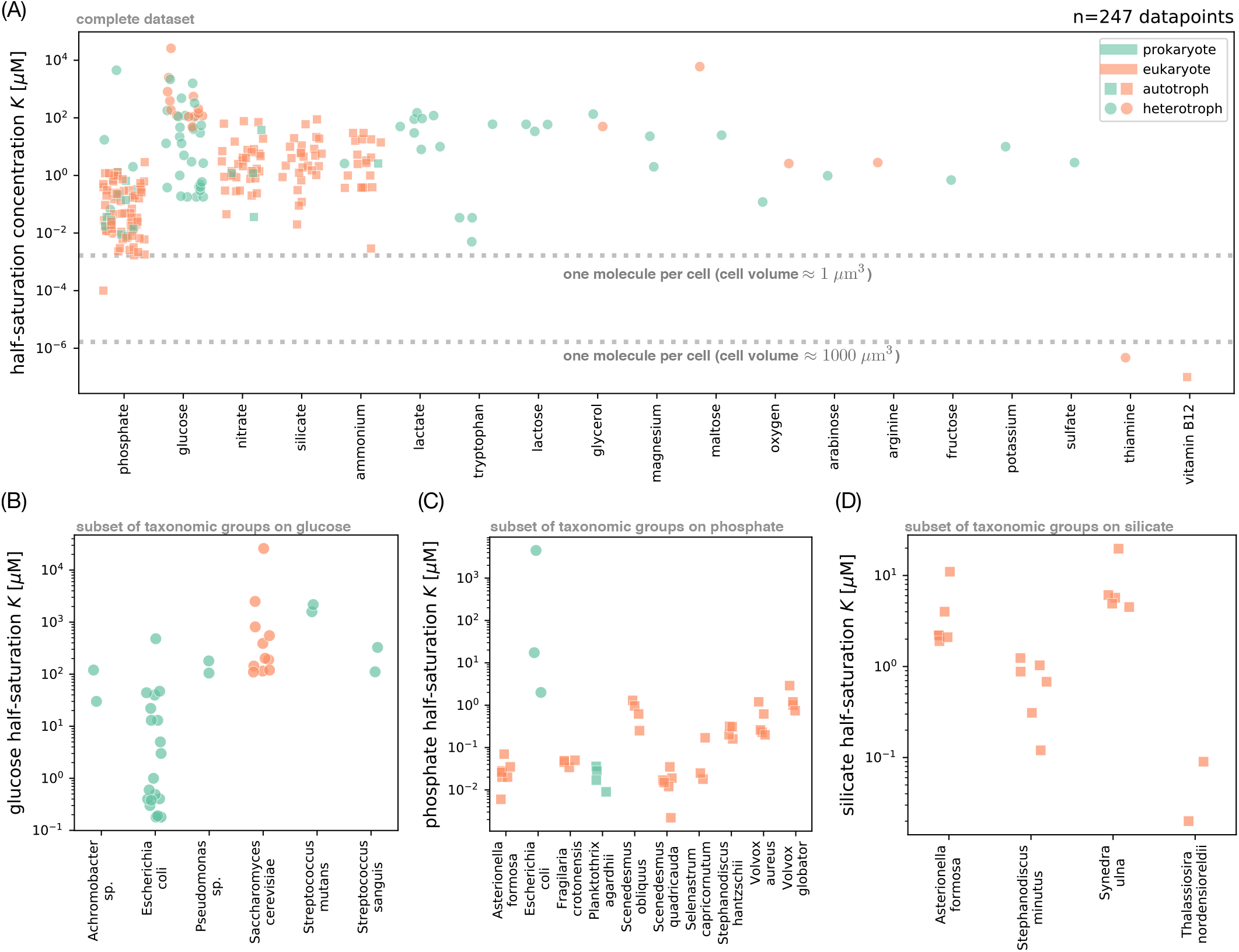
Survey of measured half-saturation concentrations. (A) Complete set of half-saturation concentrations *K* for the Monod model of growth rate (Eq. (1)) in our survey, grouped by resource (in decreasing order of number of data points). Each point represents a different measurement; color indicates whether the organism is a prokaryote (green) or eukaryote (orange), and shape indicates whether the organism can grow as an autotroph (square) or only as a heterotroph (circle). Dashed lines mark concentrations of one molecule per cell for approximate prokaryotic and eukaryotic cell volumes [45]. (B) Subset of *K* measurements from panel A for glucose, grouped by taxon (only those with at least two measurements). We use the taxonomic identity given in the original publications, where an ending in *sp*. means the isolate is a representative of the genus but was not identified at the species level. Symbols are the same as in panel A. For brevity, we use “glucose half-saturation” to refer to the half-saturation concentration for glucose as the limiting nutrient. (C) Subset of *K* measurements from panel A for phosphate, grouped by taxon (with at least three measurements). (D) Subset for silicate, grouped by taxon (with at least two measurements). See the Supplemental Information for additional plots with *K* measurements for nitrate (Fig. S4A) and ammonium (Fig. S4B).

Measured values of the half-saturation concentration *K* vary over several orders of magnitude, ranging from below 10^−6^ μM (for thiamine and vitamin B12) to above 10^4^ μM (for one glucose measurement). This variation is not attributable to measurement uncertainties, which never exceeded 20% in the studies that reported them. It also is not an artifact of technical aspects of the measurements (Fig. S2) such as temperature (linear regression, *R*^2^ ≈ 0.089, *p* ≈ 1.2 × 10^−5^) or experimental method (linear regression, *R*^2^ ≈ 0.160, *p* ≈ 1.3 × 10^−3^), nor does the variation appear to be systematically biased by experimental design such as the degree of pre-acclimation to the growth medium (Fig. S3). We furthermore find no evidence for a major bias from simultaneous limitation (colimitation) for other resources besides the focal resource (Supplementary Information Sec. S1).

Instead, most variation of concentrations *K* corresponds to variation in the identity of the organisms and resources themselves (Fig. S2A). Figure 2B shows a subset of measurements on glucose, which have systematic differences in *K* between taxa. For example, measurements of *S. cerevisiae* and *Streptococcus* almost all have *K* values higher than those of *E. coli* (Mann-Whitney U test, *p* ≈ 1.40 × 10^−6^). Phosphate and silicate similarly show significant variation between species (Fig. 2C,D), as do nitrate and ammonium (Fig. S4). Even within some taxa, there is large variation of *K*; glucose *K* in *E. coli* varies over three orders of magnitude (Fig. 2B). This variation within a single resource and taxon does not appear to be explained by technical covariates of the measurements (Fig. S2B), but rather corresponds to genetically-distinct strains of *E. coli* (Fig. S5), suggesting that even subspecies-level genetic variation can lead to significant differences in the half-saturation concentration *K*. Indeed, Ferenci [46] reported single target genes, like the membrane-associated lamB or the stress-factor rpoS, that affect the half-saturation concentration of *E. coli* on glucose when mutated. The genetic differences in our dataset are mostly unknown, but we grouped *E. coli* measurements by strain labels to find reproducible half-saturation concentrations for glucose within strains (e.g., ML 30, see Fig. S5A).

How can we explain this wide variation in half-saturation concentrations? Intuitively, we expect evolution to reduce *K*, since mutations that reduce *K* increase growth rate (Eq. (1)). For example, Fig. 1 shows the growth rate dependence for a hypothetical wild-type strain (green line) and a mutant (orange) with lower half-saturation *K*. Since the mutant has a greater relative growth rate advantage at low resource concentrations, there could be stronger selection pressure to reduce *K* at those low concentrations. This is hinted by some patterns in the data: for example, *E. coli* often grows in mammalian large intestines where there are few simple sugars such as glucose, while *S. cerevisiae* and *Streptococcus* often grow in high-sugar environments (fruit and the oral microbiome, respectively) [47, 48], which could explain their large difference in half-saturation concentrations for glucose.

### Variation in specific affinity has trends similar to those of the half-saturation concentration

Since *K* alone does not define the growth rate at low resource concentrations, it is essential to consider the maximum growth rate *g*^max^ or specific affinity *g*^max^/*K* as well. We show the variation in maximum growth rate *g*^max^ across resources in Fig. 3A (reported for 97.5% of all entries for half-saturation concentrations *K*; Dataset S1). The most striking feature of this data is that while maximum growth rates *g*^max^ vary less between resources than do half-saturation concentrations *K* (compare Figs. 3A and 2A), there is a clear bimodality between fast-growing heterotrophs (circles) and slow-growing autotrophs (squares). Indeed, a closer look at the covariation between *g*^max^ and *K* in autotrophs (squares in Fig. 3B) reveals that resources have comparable distributions of *g*^max^ but stratify in terms of half-saturation concentrations *K*, with the lowest values for phosphate. In particular, the distributions for phosphate and nitrate are indistinguishable in terms of maximum growth rate (Mann-Whitney U test, *p* = 0.0801), but clearly different in terms of half-saturation concentration (Mann-Whitney U test, *p* = 1.28 × 10^−12^). Also, the species differences in maximum growth rate on glucose and phosphate are less pronounced (Fig. S6) and more of the variation can be explained by experiment temperature (Figs. S7 and S8) compared to variation in *K*.

**FIG. 3.**
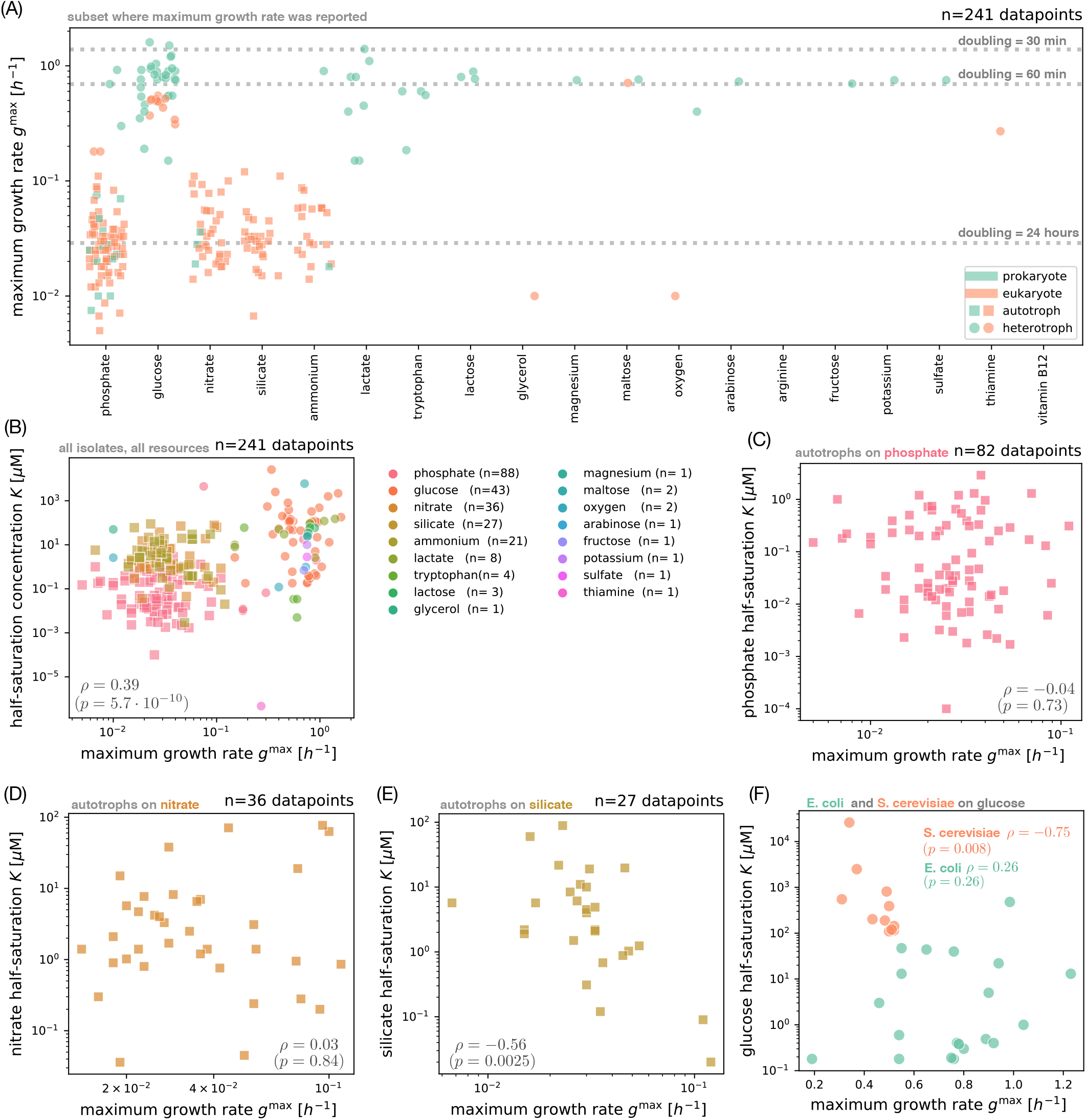
Survey of maximum growth rates and trait correlations. (A) Empirical maximum growth rates *g*^max^ for the microbial isolates in our survey. There are slightly fewer data points for maximum growth rate compared to half-saturation concentrations in Fig. 2A, since some publications only reported the half-saturation concentration. Markers indicate whether the organisms can grow as an autotroph (square) or only as a heterotroph (circle); colors indicate if the isolate is prokaryotic (green) or eukaryotic (orange). Dashed lines mark reference doubling times. (B) Covariation of maximum growth rate *g*^max^ and half-saturation concentration *K* across the entire set of isolates from panel A. Here colors indicate the limiting resource, with the number of measurements *n* given in parentheses. Marker shapes (squares are autotrophs, circles are heterotrophs) are the same as in panel A. We compute the Spearman rank correlation *ρ* and *p*-value across the pooled set of isolates. (C) Subset of measurements from panel B for phosphate (only autotroph isolates shown). (D) Subset of measurements from panel B for nitrate. (E) Subset of measurements from panel B for silicate. (F) Covariation between maximum growth rate *g*^max^ and half-saturation concentration *K* on glucose for measurements of *E. coli* (green) and *S. cerevisiae* (orange), with Spearman rank correlations *ρ* and *p*-values by species.

We can also compute the specific affinity *g*^max^/*K* for each data point. Figure S9 shows that the variation in specific affinity is similar to variation in *K*: the variation spans orders of magnitude, even for single species, and there are systematic differences between taxa (e.g., *E. coli* compared to *S. cerevisiae* and *Streptococcus*; Mann-Whitney U test, *p* ≈ 1.20 × 10^−6^; Fig. S9B). The similarity in patterns of variation between the half-saturation concentration and specific affinity is because variation in *g*^max^/*K* is dominated by variation in *K* (Fig. S7B); on a logarithmic scale, *g*^max^/*K* depends on additive contributions from *g*^max^ and *K*, and variation in *K* is much larger than variation in *g*^max^ (compare Figs. 2A and 3A).

### There is no evidence for a tradeoff between half-saturation concentration and maximum growth rate

Many previous studies have considered the possibility of tradeoffs between *g*^max^ and *K* (positive correlation), such that genotypes growing faster with abundant resources will grow slower when resources are scarce [13, 25–28]. If this were true, evolution at high resource concentrations may select for increasing maximum growth rate *g*^max^ at the expense of the half-saturation concentration *K*, leading to high values of *K*. If we consider all organisms and resources in our data set, we do find a significant positive correlation between *g*^max^ and *K* (Spearman *ρ* ≈ 0.39, *p* ≈ 5.7 × 10^−10^; Fig. 3B). However, this correlation is an artifact of the biased sampling of organism-resource pairs, which are dominated by fast-growing heterotrophs on glucose (which tend to have higher concentrations *K*) and slow-growing autotrophs on other resources (which tend to have lower concentrations *K* compared to glucose); the correlation disappears when we separate heterotrophs (Fig. S10A,B) from autotrophs (Fig. S10C,D). If we further separate individual resources, we see no significant correlations for phosphate, nitrate, ammonium, or glucose across organisms (Figs. 3C,D and S10E–H), while there is actually a negative correlation (opposite of a tradeoff) for silicate *g*^max^ and *K* (Spearman *ρ* ≈ −0.56, *p* ≈ 0.0025; Fig. 3E). In Fig. 3F we test the covariation of *g*^max^ with *K* for two individual species (*E. coli* and *S. cerevisiae*) for a single resource (glucose). The *E. coli* data shows a positive correlation indicative of a tradeoff, but it has modest magnitude and low statistical significance (Spearman *ρ* ≈ 0.26, *p* ≈ 0.26). *Saccharomyces cerevisiae*, on the other hand, shows a positive correlation between the two traits (Spearman *ρ* ≈ −0.75, *p* ≈ 0.008). The lack of tradeoff appears irrespective of experimental method (i.e., batch or chemostat; Fig. S3B) and also holds when comparing the maximum growth rate *g*^max^ to the specific affinity *g*^max^/*K* (Fig. S11).

Much of the previous literature arguing for tradeoffs in these traits based their evidence on measurements for resource uptake kinetics [27, 28, 30, 49] rather than on population growth as we consider here. However, we find little to no correspondence between traits of uptake kinetics with traits of population growth in data points where we have measurements for both (Fig. S12) [44], consistent with previous analyses [37, 39]. It is therefore not surprising that the observed tradeoffs in uptake do not translate to tradeoffs in growth. For example, Litchman et al. [30] reported a tradeoff between uptake traits for nitrate, but we see no correlation in growth traits for nitrate (Spearman *ρ* ≈ 0.03, *p* ≈ 0.84; Figs. 3D and S11C). Altogether the absence of evidence for a systematic correlation between *K* and *g*^max^ suggests that selection for *g*^max^ does not explain the evolved variation in *K*.

### Models of population dynamics with mutations to half-saturation concentration

To test how the environmental resource concentration shapes the evolution of the half-saturation concentration *K*, we turn to a model of population dynamics with mutations altering traits of the Monod growth rate response (Methods; Supplementary Information Secs. S3–S5; Table S2). We consider a microbial population consisting of a wild-type and a mutant, with biomasses *N*_wt_(*t*) and *N*_mut_(*t*) that vary over time *t*. They grow at rates depending on the resource concentration *R* according to the Monod model (Eq. (1)), but with potentially different values of the traits *g*^max^ and *K* depending on the effect of the mutation [25, 28]. The rate at which the mutant increases or decreases in frequency compared to the wild-type is given by the selection coefficient *s* (Supplementary Information Sec. S6) [50, 51]. We show that *s* decomposes into two additive terms

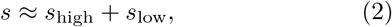

where *s*_high_ measures selection on growth at high resource concentrations, and is therefore proportional to variation in the maximum growth rate *g*^max^, while *s*_low_ measures selection on growth at low resource concentrations, and is therefore proportional to variation in the half-saturation concentration *K* (Figs. S13–S16; Supplementary Information Secs. S7–S9).

We consider selection in three prototypical regimes of population dynamics. In the first case, the population grows as a batch culture with serial transfers (Supplementary Information Sec. S3). That is, there is an initial concentration *R*_0_ of the resource, and the population grows until the resource is exhausted. Figure 4A shows these dynamics for the hypothetical wild-type and mutant strains of Fig. 1. Although the mutant has the same maximum growth rate *g*^max^ as the wild-type, its lower value of *K* allows it to continue growing fast at lower concentrations of the resource, decelerating more abruptly at the end of growth (see inset of Fig. 4A for more dramatic examples). Then a fixed amount of biomass *N*_0_ — sampled from the whole culture, so that the relative frequencies of the mutant and wild-type are preserved on average — is transferred to a new environment with the same initial concentration *R*_0_ of the resource as before, and the cycle repeats (Fig. top panel). This dilution step represents a form of mortality for the population. We refer to this regime as *fixed-bottleneck batch dynamics*, since the bottleneck of biomass between transfers is held fixed. Boom-bust dynamics such as these are believed to be common in some natural environments [52, 53], with a fixed bottleneck size being plausible for populations that serially colonize new environments [54] or are reset to a fixed density by culling [4] between cycles of growth.

**FIG. 4.**
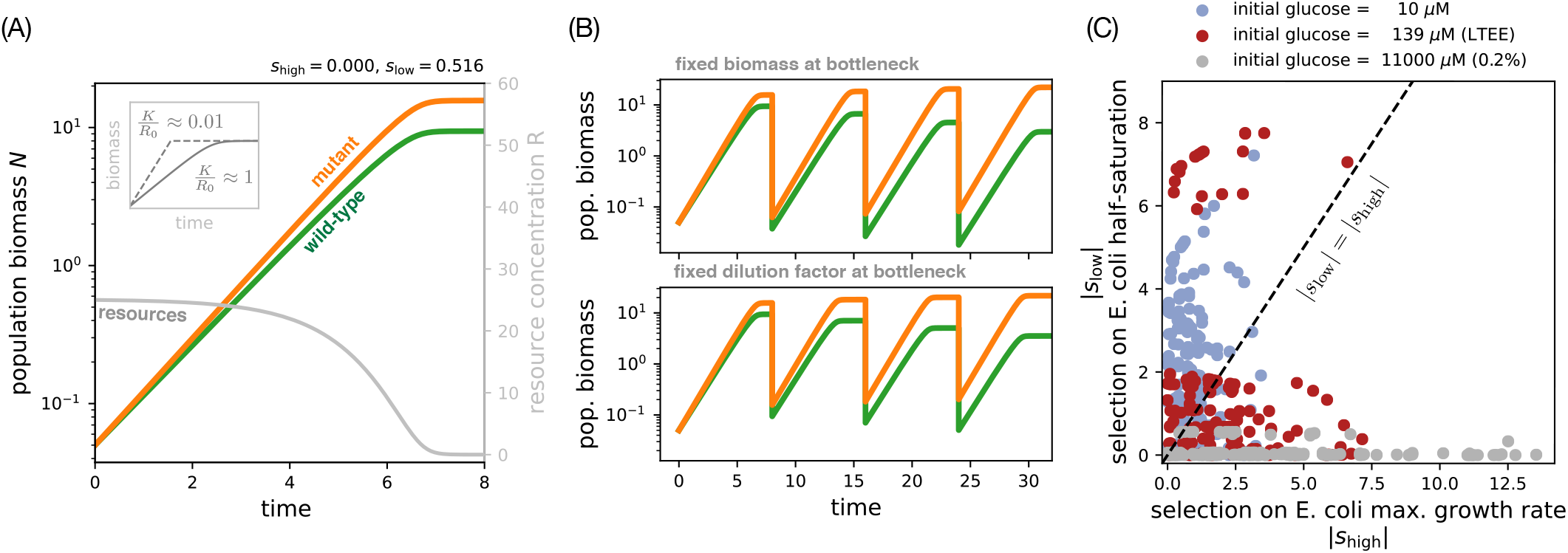
Selection on variation in half-saturation concentrations over batch population dynamics. (A) Simulated growth of wild-type (green) and mutant (orange) strains competing under batch dynamics, with the transient resource concentration (gray) on the right vertical axis (Supplementary Information Sec. S3). The strain pair is the same as in Fig. 1; the initial resource concentration is *R*_0_ = 25, with strains at equal initial frequencies and equal yields. (B) The same strain competition from panel A continued over multiple growth cycles under fixed-bottleneck batch dynamics (top panel, *N*_0_ = 0.01) and fixed-dilution batch dynamics (bottom panel, *D* = 100). (C) Each point represents the predicted selection coefficients |*s*_high_| and |*s*_low_| (Eq. (2); Supplementary Information Sec. S8) for pairs of *E. coli* isolates with measured growth traits on glucose (from Fig. 2D). The three colors represent different glucose concentrations. We assume the isolates in each pair start competing at equal initial frequencies, set the initial cell density to *N*_0_ = 4.6 × 10^5^ cells/mL, and use a biomass yield of *Y* = 3.3 × 10^8^ cells/μmol glucose measured by a previous study [23].

The second regime is the same as the first, except instead of transferring a fixed amount of biomass to the next cycle, we transfer a fixed fraction 1/*D*, where *D* is the dilution factor (Fig. 4B, bottom panel); we therefore refer to this regime as *fixed-dilution batch dynamics*. Note that the dilution factor *D* and the bottleneck biomass *N*_0_ are related according to *D* = *R*_0_*Y/N*_0_ + 1, where *Y* is the yield (biomass produced per unit resource; Supplementary Information Sec. S3). These dynamics are plausible for populations that experience a constant death rate between growth cycles or are regularly purged by the environment, as believed to occur in the human gut microbiome [55]. This case is also the most common protocol in laboratory evolution experiments owing to its simplicity [56]. While the differences between these two regimes of batch dynamics may appear to be subtle (comparing the two panels of Fig. 4B), we will show later that these two dilution protocols have different dependences on the resource concentration, which lead to different evolutionary outcomes.

Finally, we also consider the regime of *chemostat dynamics*, where the population grows as a continuous culture with a constant supply of the resource and a constant dilution rate *d* (Supplementary Information Sec. S5). Chemostats are used as devices for experimental evolution [12, 22] and the same dynamics are often applied to describe natural populations in the ocean [13, 57].

### Selection quantifies variation in growth traits between isolates at different resource concentrations

We previously observed wide variation in half-saturation concentrations *K* (Fig. 2A) and maximum growth rates *g*^max^ (Fig. 3A) across isolates, but the significance of this variation is difficult to assess by itself. For example, glucose *K* for *E. coli* varied across four orders of magnitude, but how significant is this variation for evolution? Our model of selection under different population dynamics gives us precisely the metric to quantify this variation. We demonstrate this in Fig. 4C by calculating the two components of selection (Eq. (2)) for hypothetical competitions between all pairs of *E. coli* isolates measured on glucose. We do this for batch dynamics starting at different initial concentrations *R*_0_ of glucose. While selection on variation in *g*^max^ (*s*_high_) always increases with higher *R*_0_, selection on variation in *K* (*s*_low_) depends non-monotonically on the concentration *R*_0_, such that selection is maximized at some intermediate concentration (Fig. S17, Supplementary Information Sec. S10). Intuitively, this optimal concentration approximately equals the half-saturation concentration *K* itself (Fig. S17C). On the other hand, if the resource concentration *R*_0_ also increases the initial population size *N*_0_ (i.e., transfer from a pre-growth cycle with fixed dilution factor), selection on variation in *K* depends monotonically on *R*_0_ and is maximized at the lowest concentration (Fig. S18).

We calculate selection between *E. coli* isolates at 10 μM glucose, which is in the middle of the range of observed half-saturation concentrations *K*, as well as at two higher concentrations corresponding to the conditions of the *E. coli* LTEE (139 μM) [58] and a common laboratory concentration (11000 μM ≈ 0.2% w/v). Figure 4C indeed shows that variation in the value of *K* is highly significant for evolution at concentrations around the half-saturation concentration, whereas at the highest concentration, selection on the variation in *K* is small compared to the selection in *g*^max^.

### The half-saturation concentration evolves downward over successive mutations

With our model of population dynamics, we can predict how the traits of the Monod growth rate response (Eq. (1)) will evolve over long times. For simplicity, we focus on the “strong-selection weak-mutation” (SSWM) regime of evolutionary dynamics, where each new mutation either fixes or goes extinct before the next mutation arises (Fig. S19; Supplementary Information Sec. S11) [59].

We first simulate a population growing under fixed-bottleneck batch dynamics, with an initial half-saturation concentration *K* that is higher than the external resource concentration *R*_0_; the population therefore decelerates gradually into starvation over each growth cycle (Fig. 5A, left inset). Mutations then regularly arise and alter the value of *K* with a random effect size (Fig. S19; Supplementary Information Sec. S11). Each mutation stochastically fixes or goes extinct according to a fixation probability, which depends on the mutation’s selection coefficient. Over time these beneficial mutations accumulate and the half-saturation concentration *K* systematically decreases. By the end of the simulation, the half-saturation concentration *K* is 1000 times smaller than the resource concentration *R*_0_, leading to growth curves that grow much faster and abruptly decelerate into starvation (Fig. 5A, right inset).

**FIG. 5.**
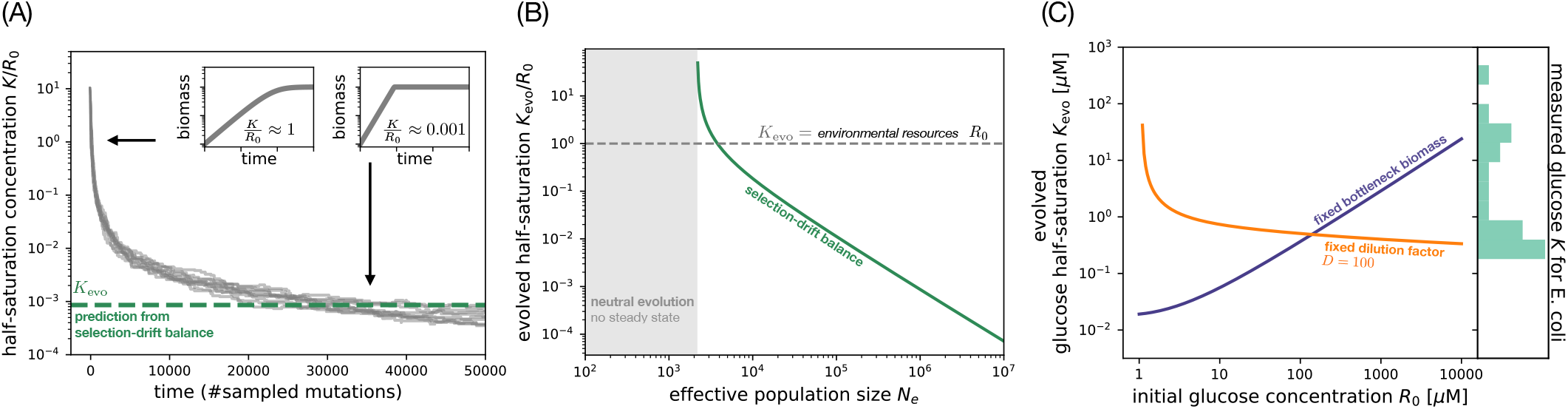
Evolution of the half-saturation concentration. (A) The half-saturation concentration *K* evolving under fixed-bottleneck batch dynamics. Each gray line is one of 10 independent stochastic simulations using an effective population size *N*_e_ = 1000 and mutation effects *κ* drawn from a uniform distribution (Supplementary Information Sec. S11). The insets show the growth curve in a single batch cycle before adaptation (left inset) and at the final state (right inset). The green dashed line marks our prediction *K*_evo_ at selection-drift balance. (B) Evolved half-saturation concentration *K*_evo_ as a function of the effective population size *N*_e_. In the gray region, the effective population size is too small and all evolution is neutral. If *N*_e_ is sufficiently large (white region), the evolved half-saturation *K*_evo_ has selection-drift balance along the green line. Parameters are |*κ*_max_| = 0.001, *g*^max^ = 1, *N*_0_ = 0.01, and *Y* = 1 for both strains. (C) The evolved glucose half-saturation *K*_evo_ as a function of initial glucose concentration *R*_0_ for two regimes of batch dynamics: fixed-bottleneck dynamics (blue line) and fixed-dilution dynamics (orange line). We use parameters based on the LTEE: *N*_0_ = 4.6 × 10^5^ cells/mL (for fixed-bottleneck case), *D* = 100 (for fixed-dilution case), *N*_e_ = *VN*_0_ where *V* =10 mL, *g*^max^ = 0.888/h, and *Y* = 3.3 × 10^8^ cells/μmol [23]. We also set *κ*_max_ = 6 × 10^−6^ (Fig. S27). On the right axis is a histogram of glucose half-saturation *K* data for *E. coli* isolates (from Fig. 2B).

Such an abrupt arrest is, for example, realized by *E. coli* in glucose-limited batch culture through a dynamic surge in gene expression late in the growth cycle [60], often involving the use of separate transporters with lower Michaelis-Menten constants [61]. The presence of these transporter systems has been raised as evidence for evolutionary adaptation of the species at micromolar glucose concentrations [8, 61, 62]. But our model shows that a feast-and-famine environment dominated by concentrations orders of magnitude higher would still allow *E. coli* to evolve the low half-saturation concentrations *K* observed in existing strains.

### Adaptation in the half-saturation concentration stalls when it reaches selection-drift balance

The value of *K* does not evolve downward forever; in Fig. 5A adaptation slows down and the half-saturation concentration levels off after a few tens of thousands of mutations, even though there is no change in the supply of beneficial mutations. This occurs because selection on beneficial mutations is inhibited by random demographic fluctuations in the population, known as genetic drift [63]. The strength of genetic drift is measured by 1/*N*_e_, where *N*_e_ is the effective population size (for the variance in mutant frequency change per unit time) [64, 65]; smaller populations experience greater fluctuations. In the simplest cases, *N*_e_ is proportional to the actual (“census”) population size, but in more complex systems *N*_e_ may depend on other aspects of demography (such as spatial dynamics [66] or age structure [67]) as well as additional sources of noise in the population dynamics [68].

Beneficial mutations will therefore no longer fix with high probability if their selection equals genetic drift, a condition known as selection-drift balance [69–71]:

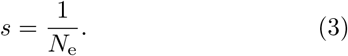

Selection-drift balance occurs in our model under batch dynamics because the growth deceleration phase becomes shorter as *K* decreases over evolution (insets of Figs. 4A and 5A), which means there is weaker selection to reduce it further. Once the half-saturation concentration *K* becomes sufficiently small, selection is no longer strong enough to overcome genetic drift (Supplementary Information Sec. S12; Fig. S20).

By combining Eqs. (2) and (3), we can calculate the value of the evolved half-saturation concentration at which selection-drift balance occurs (Fig. S21). For typical regimes of the parameters, the evolved concentration is approximately (Supplementary Information Sec. S13)

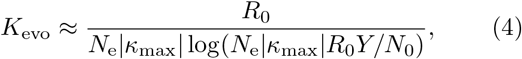

where *κ*_max_ is the maximum effect size of a beneficial mutation reducing *K*. We calculate an example of *K_evo_* in Fig. 5A (dashed green line), which corresponds well with the simulations. This result is robust to a wide range of effective population sizes and frequency-dependent effects (Fig. S22; Supplementary Information Sec. S11). We also observe an equivalent result for the adaptation of the specific affinity *g*^max^/*K* (Fig. S23; Supplementary Information Sec. S14) instead of the half-saturation concentration *K* alone.

One salient feature of Eq. (4) is that the evolved half-saturation concentration *K*_evo_ scales inversely with the effective population size *N*_e_, as shown in Fig. 5B. That is, larger populations or those with lower genetic drift can evolve proportionally lower half-saturation concentrations *K*_evo_ that are orders of magnitude lower than the environmental resource concentration *R*_0_. This potentially explains why we observe such low values of *K* for many organisms and resources (Fig. 2); this also explains why these half-saturation concentrations are difficult to measure from time-series data, since low half-saturation concentrations produce extremely abrupt deceleration at the end of growth (insets of Figs. 4A and 5A and Fig. S24; Supplementary Information Sec. S15). Hints of the influence of *N*_e_ are found in ammonia-oxidizing archaea and bacteria from marine environments, which tend to have lower half-saturation concentrations than isolates from soil [18]. Our scaling relationship Eq. (4) suggests that this ordering can arise from the smaller effective population size *N*_e_ for spatially-structured environments like soil.

The other important feature of Eq. 4 is the dependence of the evolved half-saturation concentration *K*_evo_ on the resource concentration *R*_0_. For a fixed effective population size *N*_e_, there is an optimal value of *R*_0_ that minimizes the evolved concentration *K*_evo_ (left insets of Fig. S21A,B), just as we observed for selection on individual mutations (Fig. S17). We note that for sufficiently low values of the effective population size *N*_e_, genetic drift is stronger than selection on any mutation *κ* (Fig. S20A), and so the half-saturation concentration *K* evolves neutrally (gray region in Fig. 5B).

In contrast to batch dynamics, selection under chemostat dynamics does not depend on the half-saturation concentration *K* itself (Supplementary Information Sec. S9). Intuitively, this is because reductions in *K* cause the environmental resource concentration to decrease proportionally (Supplementary Information Sec. S5), such that the growth rate remains constant. Not only does this keep a constant strength of selection on new mutations, but the effective population size will actually increase as *K* evolves lower, making beneficial mutations even easier to fix. Therefore selection-drift balance never occurs for *K* under chemostat dynamics; the half-saturation concentration *K* will continue to evolve downward until adaptation is limited by the supply of mutations or other factors (Discussion). Note that selection-drift balance also does not occur for mutations to the maximum growth rate *g*^max^ under either batch or chemostat dynamics, since selection does not depend on the magnitude of growth rate (Supplementary Information Secs. S8 and S9).

### Population dynamics can decouple the evolved half-saturation concentration from the resource concentration

In general the effective population size *N*_e_ that controls genetic drift may be shaped by a variety of demographic factors besides the census population size [65]. However, in well-mixed batch cultures, *N_e_* is primarily determined by the number of cells at the bottleneck of each transfer [69, 72]; we assume that other sources of stochasticity (such as individual cell division events) are much weaker than the sampling noise of these transfers. Therefore the effective population size *N_e_* is proportional to the bottleneck biomass *N*_0_ (assuming constant biomass per cell).

Under fixed-bottleneck batch dynamics, the effective population size *N_e_* is thus an independent parameter of the population, so that the strength of genetic drift does not depend on the resource concentration (Fig. S25A). In this case, the evolved trait *K*_evo_ is in approximately linear proportion to the resource concentration *R*_0_ (Eq. (4); Figs. 5C and S26A), making the evolved half-saturation concentration a biomarker of the resource’s environmental concentration. This is consistent with our original speculation about the systematic differences in glucose *K* between *E. coli* and *S. cerevisiae*, owing to the different glucose availability in their different environments.

However, for fixed-dilution batch dynamics, the bottleneck biomass *N*_0_, and therefore the effective population size *N_e_*, are coupled to the resource concentration *R*_0_ because the dilution factor *D* is fixed: *N_e_* ∝ *N*_0_ = *R*_0_*Y*/(*D* – 1) (Supplementary Information Sec. S3). This coupling occurs because increasing the resource concentration increases the biomass at the end of each growth cycle, but then the fixed dilution factor means that this must also increase the biomass at the bottleneck. The scaling of *N*_e_ with *R*_0_, though, cancels out the scaling of *K*_evo_ with *R*_0_ in Eq. (4), leading to an evolved half-saturation concentration *K*_evo_ that is approximately independent of the environmental concentration *R*_0_ (Figs. 5C and S26B). Conceptually, fixed-dilution batch dynamics do not allow the strength of selection to be tuned independently from genetic drift: the decrease in selection magnitude on *K* with higher resource concentration *R*_0_ is compensated by weaker genetic drift, due to a higher effective population size *N*_e_ (Fig. S25B). Thus, the population dynamics decouple the evolved half-saturation concentration of the organism from the environmental concentration.

This has major consequences for interpreting empirical variation. We predict the evolved half-saturation concentration *K*_evo_ for *E. coli* on glucose as a function of glucose concentration *R*_0_ in Fig. 5C, using parameters estimated from the LTEE (Fig. S27). On the same plot, we show a histogram of all measured glucose *K* values for *E. coli* (from Fig. 2B) on the right vertical axis. We see that, under fixed-bottleneck batch dynamics, we would expect *E. coli* to have evolved in glucose concentrations above 100 μM to account for the observed half-saturation concentrations. However, under fixed-dilution batch dynamics, the evolved half-saturation concentration depends so weakly on the environmental concentration that almost any concentration of glucose is possible to explain the data.

## DISCUSSION

### Modeling insights to interpret half-saturation data

Since it is often difficult to measure resource concentrations and population dynamics in natural environments, can we use the evolved half-saturation concentration *K* as a biomarker to infer them? This logic is often implicit in environmental studies, which attempt to draw conclusions about the environmental conditions of an isolate based on its abilities to grow at different resource concentrations [16–19]. However, our model shows that it is not as simple as assuming the half-saturation concentration *K* for a resource is proportional to its concentration in the environment, since that proportionality is altered by the population dynamics, at least through the effective population size *N*_e_ (Eq. (4)). In particular, this proportionality is confounded in the case of fixed-dilution batch dynamics, where the evolved half-saturation concentration *K* is largely independent of the resource concentration *R*_0_ (Fig. 5C).

Under fixed-bottleneck batch dynamics, though, the linear scaling of *K* with *R*_0_ does approximately hold. In this case, one can compare two populations with unknown, but identical effective population sizes *N*_e_ and mutation effects *κ*; for example, two isogenic populations located at different points along a resource gradient. In this case, one can calculate the ratio of evolved half-saturation concentrations *K*_evo_ for the two populations to estimate the ratio of resource concentrations. But in many scenarios, one might not even know the type of bottlenecks the population is experiencing. To classify the population dynamics as fixed-bottleneck or fixed-dilution, one could correlate a set of evolved concentrations *K*_evo_ with their different resource concentrations *R*_0_; a strong linear correlation would support fixed-bottleneck batch dynamics, while little to no correlation would indicate fixed-dilution batch or chemostat dynamics.

### Role of the mutation supply in shaping evolved half-saturation concentrations

We have focused on the role of selection-drift balance as a null model for the evolved variation in half-saturation concentrations, since the competition between selection and genetic drift is a universal feature of all evolving populations. In doing so we have assumed the supply of mutations on *K* is constant, but real populations will at some point run out of beneficial mutations on the trait value *K*, potentially reaching this mutation-selection balance before selection-drift balance [70]. Many mutations will also be pleiotropic, affecting both the half-saturation concentration *K* and the maximum growth rate *g*^max^ (as well as possibly other traits) simultaneously. The correlation between pleiotropic effects on both traits is important: if pleiotropy is synergistic, so that mutations that decrease *K* also tend to increase *g*^max^, then the population might evolve lower *K* than otherwise expected since its selection is enhanced by additional selection on *g*^max^. On other other hand, if there is a tradeoff between *K* and *g*^max^, the population might evolve higher *K* if its selection is outweighed by selection for higher *g*^max^. Indeed, this is what appears to have happened in the LTEE, where *K* for glucose actually increased over the first 2000 generations, but that was offset by a stronger improvement in the maximum growth rate *g*^max^ [23].

Such a tradeoff between *K* and *g*^max^ is interesting both for its consequences on the stochiometric composition of community biomass [49, 73] as well as from an evolutionary point of view, since the population can then diversify into stably-coexisting lineages. While there is significant theoretical work on this hypothesis [25–28], it has limited empirical evidence. Some of these previous studies claiming tradeoffs found them only in parameters for the Michaelis-Menten model of resource uptake [27, 28, 30, 49, 74], which we and others have shown are not equivalent to parameters of the Monod model of growth (Fig. S12) [37, 39]. In the larger set of data we have collected in this work (Fig. 3F), we find no compelling evidence of a correlation; *E. coli* shows a weak but insignificant tradeoff, while *S. cerevisiae* shows a slight synergy [75].

Interpretation of this tradeoff (or lack thereof) is also complicated by the sample of strains and environmental conditions being considered. For the tradeoff to affect the evolved half-saturation concentration as we have discussed, the tradeoff must exist across the entire spectrum of spontaneous mutations available to an organism (i.e., there is an underlying physiological constraint). This has also been the underlying assumption of previous models on this topic [25–28]. Testing this would require a distribution of *K* and *g*^max^ values over a large mutant library in a single environment, which has not been measured to our knowledge. An experimental study in *E. coli* [31] reports a tradeoff between half-saturation concentration *K* and maximum growth rate *g*^max^, but this screen was restricted to mutations in the single gene lamB, which may not be representative of genome-wide mutations. However, even in the absence of an underlying correlation in mutation effects, such a tradeoff could still emerge across clones within a rapidly evolving population, at least transiently [76, 77]. Further systematic measurements of these traits within and between populations will be necessary to resolve the issue of a tradeoff in the future.

### Other factors shaping evolved half-saturation concentrations

Besides mutation supply, there are other phenomena that may lead to different evolved outcomes for the half-saturation concentration *K*. One important assumption in our model is that we only consider a single resource, whereas real populations are dependent on several resources [78], including those from biotic sources such as cross-feeding and predation. Some of these resources may be rarely or never limiting, and therefore their half-saturation concentrations *K* will evolve only as byproducts of selection on mutations for other traits. In this sense many observed half-saturation values may actually be spandrels, an evolutionary term (defined in analogy with the architectural structure) for traits that evolve for reasons other than direct selection [79]. Selection for other traits may occur simply because competition in natural environments is likely more complex and could include lag phases [51] and other strategies for low-resource survival [5, 80–82]. On the other hand, multiple resources could also be simultaneously colimiting [32, 33]. While we have shown how colimitation under measurement conditions affects estimates of *g*^max^ and *K* (Supplementary Information Sec. S1), the effect of colimitation, as well as more complex sources of nutrients such as cross-feeding and predation, on the evolution of these traits remains an important problem for future work.

We can predict the consequences of relaxing other assumptions in our model as well. For example, simultaneous competition of multiple mutations (clonal interference) generally reduces the efficacy of selection [83, 84], which would make it more likely to evolve higher half-saturation concentrations than what we predict from SSWM dynamics. Another assumption in our model is that the population under batch dynamics always grows until complete exhaustion of the resources during each cycle, but earlier transfers could reduce the amount of growth occurring during deceleration, which would reduce selection on the half-saturation *K*. However, the population may adapt its maximum growth rate to simply saturate earlier and restore selection on its deceleration phase. Finally, populations may also have higher than expected *K* values if they simply have not had enough time to reach selection-drift balance, which takes a timescale of order *N*_e_ generations (Fig. S22) [85].

### Population dynamics are essential for understanding microbial ecology

Broadly speaking, our results provide a valuable example of how ecological traits are influenced by factors other than abiotic environmental features. In particular, we have shown how population dynamics can confound our naive expectations for the evolutionary fate of such traits. While here we have focused on the role of genetic drift, other potentially important factors include mutation supply, pleiotropy, recombination, and spatial structure. Altogether our results mean that the half-saturation concentration *K* may not be a reliable biomarker of environmental resource concentrations. This does not mean that *K* evolves independently of the environment, however. Rather, it is linked to additional environmental processes like the bottleneck between growth cycles. To understand the systematic differences between species, we need to know not only the resource concentrations they have evolved in, but also which type of population dynamics best reflects the time scales of growth, death, and resource supply in their environment of origin.

## Materials and Methods

### Literature survey of measured growth rate dependence on resources

We collected 247 measurements of Monod model parameters (*K* and *g*^max^; Eq. (1)) through a targeted literature search that included prior surveys and reviews [41, 43], the phytoplankton trait database (130 data points) by Edwards et al. [44], as well as original research papers. In all but two cases, we traced data from surveys and reviews back to their original papers, which we report in Dataset S1 (sheet 1). We included only experiments that directly measured population growth rates, rather than nutrient uptake rates or respiration. We excluded measurements where the actual limiting resource was unclear, such as measurements in rich medium with added glucose. Where possible we checked the raw data of growth rate over resource concentrations to determine if the focal resource concentration was measured up to saturation and had sufficient sampling of concentrations around *K*. For a subset of measurements of *E. coli* on glucose, we also checked for the concentration of a nitrogen source to determine the relative impact of colimitation (Dataset S1, sheet 2; Supplementary Information Sec. S1). If the original *K* value was reported as weight per volume, we converted these into units of micromolar (μM) using the calculated molecular weight of the compound’s chemical formula. We preserved significant digits from the original studies. See Dataset S1 for more details.

### Models of population dynamics

We mathematically model population dynamics using systems of ordinary differential equations for the wildtype and mutant biomasses as well as the extracellular resource concentration (Supplementary Information Secs. S3 and S5). We numerically integrate these equations using standard algorithms in Scipy [86] (Supplementary Information Sec. S4).

---

**FIG. S1.**
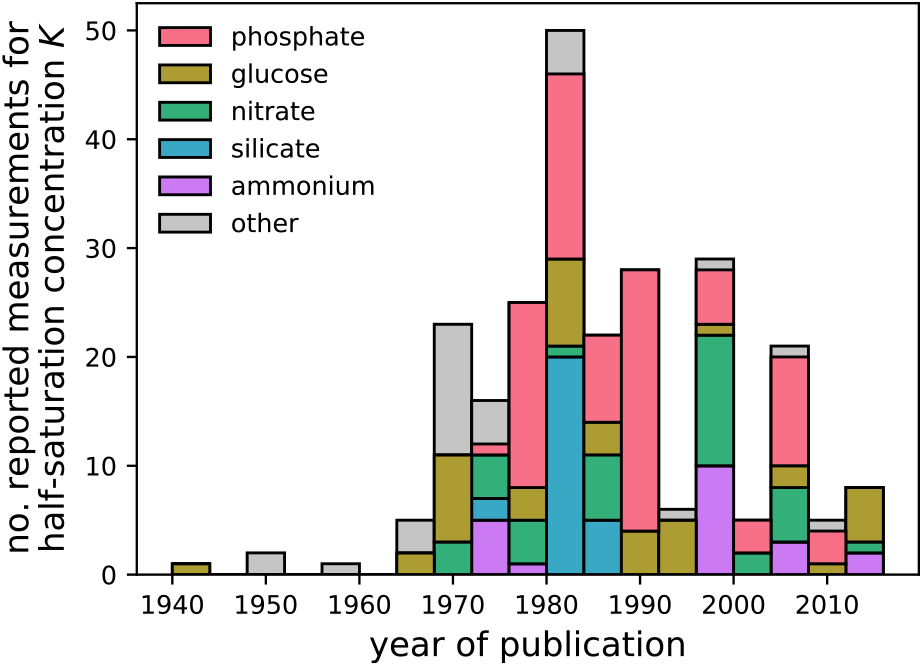
Historical trends of half-saturation concentration measurements. The number of measured half-saturation concentrations *K* published in peer-reviewed journals aggregated by year, based on our literature survey (Dataset S1). Colors indicate the number of measurements for individual resources.

**FIG. S2.**
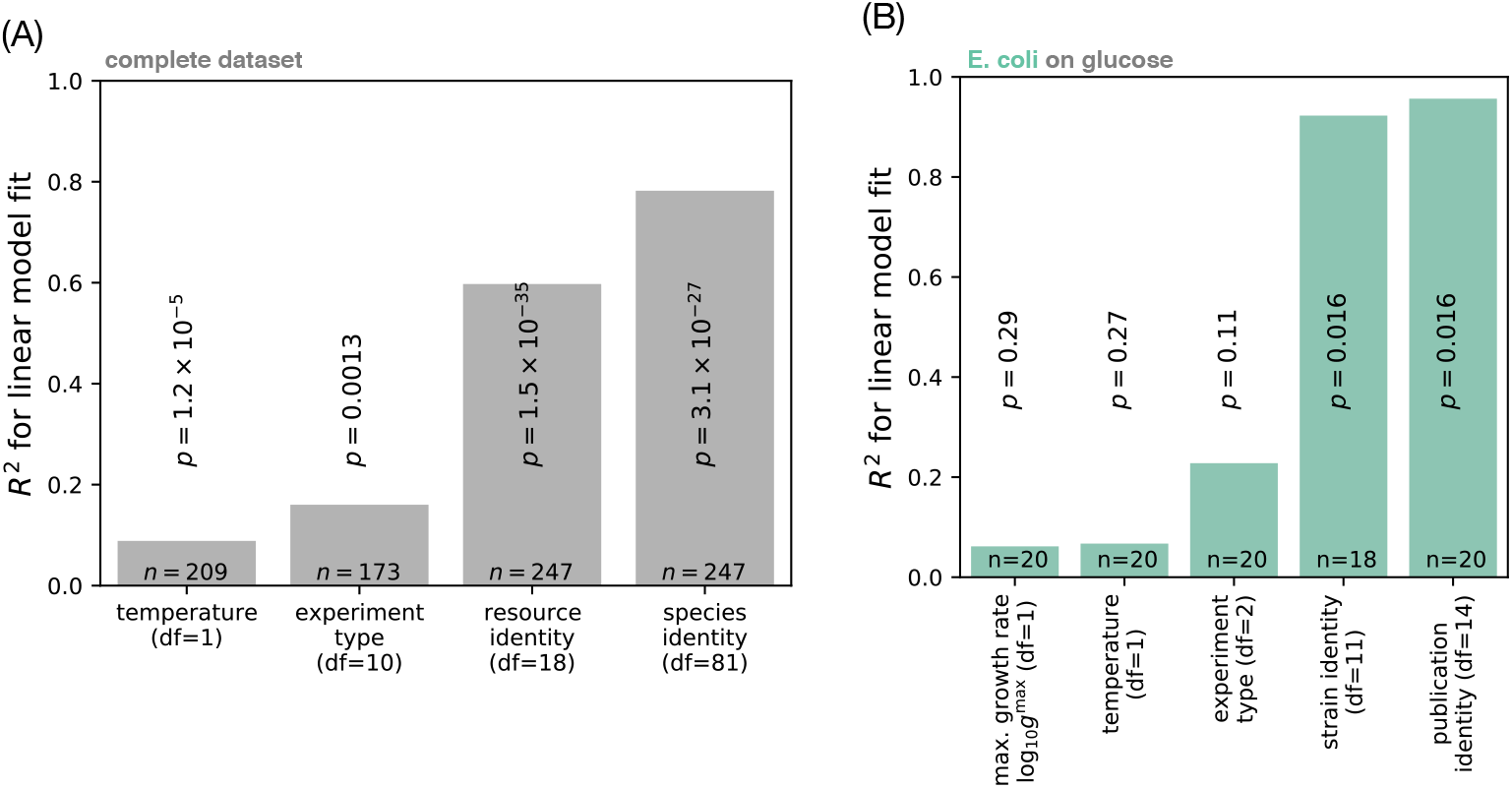
Comparison of technical covariates for the half-saturation concentration. (A) Linear regressions of technical covariates against half-saturations log_10_ *K* from the complete dataset (Fig. 2A), with degrees of freedom (df), number of data points (*n*), and *p*-values indicated. Each bar represents a separate regression fit, where *R*^2^ measures the variation explained by a single variable as predictor for the half-saturation concentration. (C) Linear regressions of technical covariates against glucose half-saturations log_10_ *K* for all *E. coli* measurements (shown in Fig. 2B).

**FIG. S3.**
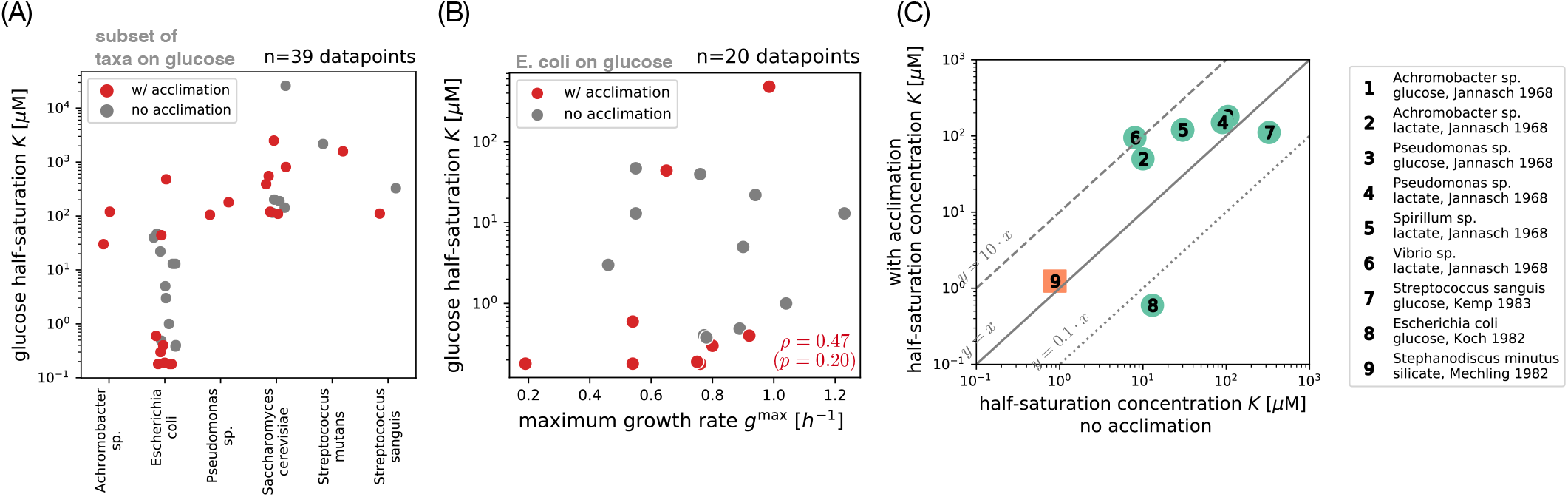
Comparison of half-saturation measurements with and without acclimation. (A) Empirical half-saturation concentrations for glucose, grouped by taxon (only those with at least two measurements). The data shown here are identical to Fig. 2B, but colors indicate which measurements included a phase of acclimation (red). We infer acclimation from the type of experiments used to measure the half-saturation concentration: For *batch solid culture*, growth rate is inferred from the area increase of single cell colonies on agar plates. For *batch experiments*, the growth rate is observed from exponential phase of a liquid culture with varying initial resource concentration. For *chemostat experiments*, the residual resource concentration is observed in steady state with varying growth rate by tuning the rate of liquid outflow. For *serial transfer experiments*, the growth is only measured in exponential phase after multiple transfers. We consider measurements to be acclimated if they derive from chemostat or serial transfer experiments. (B) Covariation between maximum growth rate *g*^max^ and glucose half-saturation *K* for isolates of *E. coli*. The data shown here are identical to *E. coli* data points in Fig. 3F. We calculate the Spearman rank correlation *ρ* and *p*-value across all isolates with acclimation (red dots). (C) Pairwise comparison of half-saturation measurements before and after acclimation. We identify a subset of publications in our database (see legend) which have explicitly tested the effect of acclimation. Each publication has two measurements for the organism’s half-saturation concentration *K* which we report together with full citations in our database (Dataset S1). A black diagonal line indicates exact match between measurements with and without acclimation, with diagonal lines in dashes (*y* = 10*x*) and dots (*y* = 0.1*x*) as visual guides for the eye.

**FIG. S4.**
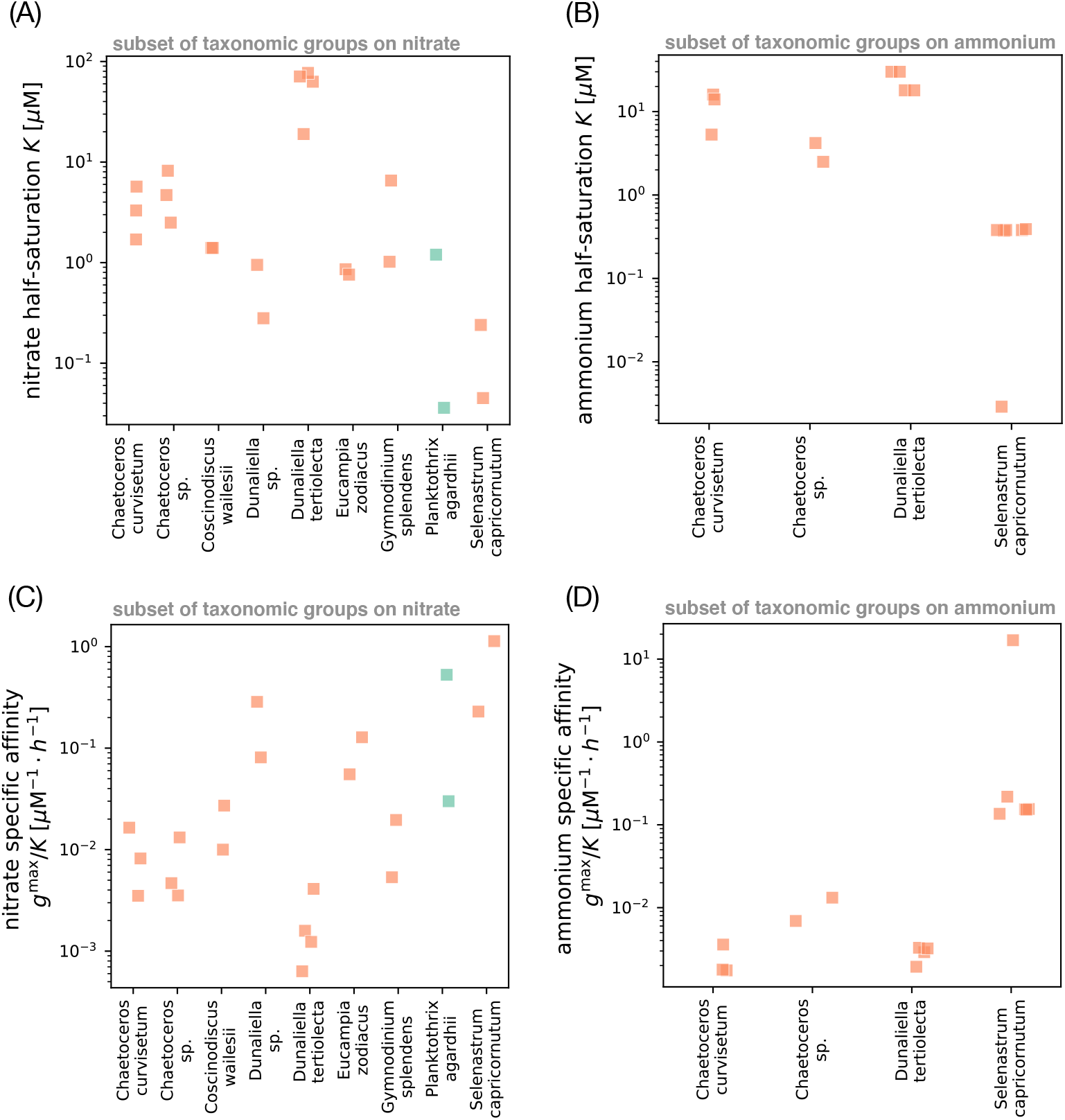
Survey of half-saturation concentrations and specific affinities for nitrate and ammonium in our survey. (A) Subset of *K* measurements from Fig. 2A for nitrate, grouped by taxon (only those with at least two measurements). Symbols are the same as in Fig. 2A: Color indicates whether the organism is a prokaryote (green) or eukaryote (orange), and shape indicates whether the organism can grow as an autotroph (square) or only as a heterotroph (circle). We use the taxonomic identity given in the original publications, where an ending in *sp*. means the isolate is a representative of the genus but was not identified at the species level. (B) Subset of *K* measurements from Fig. 2A for ammonium, grouped by taxon (with at least two measurements). (C) Subset of *g*^max^/*K* measurements from Fig. S9A for nitrate, grouped by taxon (with at least two measurements). (D) Subset of *g*^max^/*K* measurements from Fig. S9A for ammonium, grouped by taxon (with at least two measurements).

**FIG. S5.**
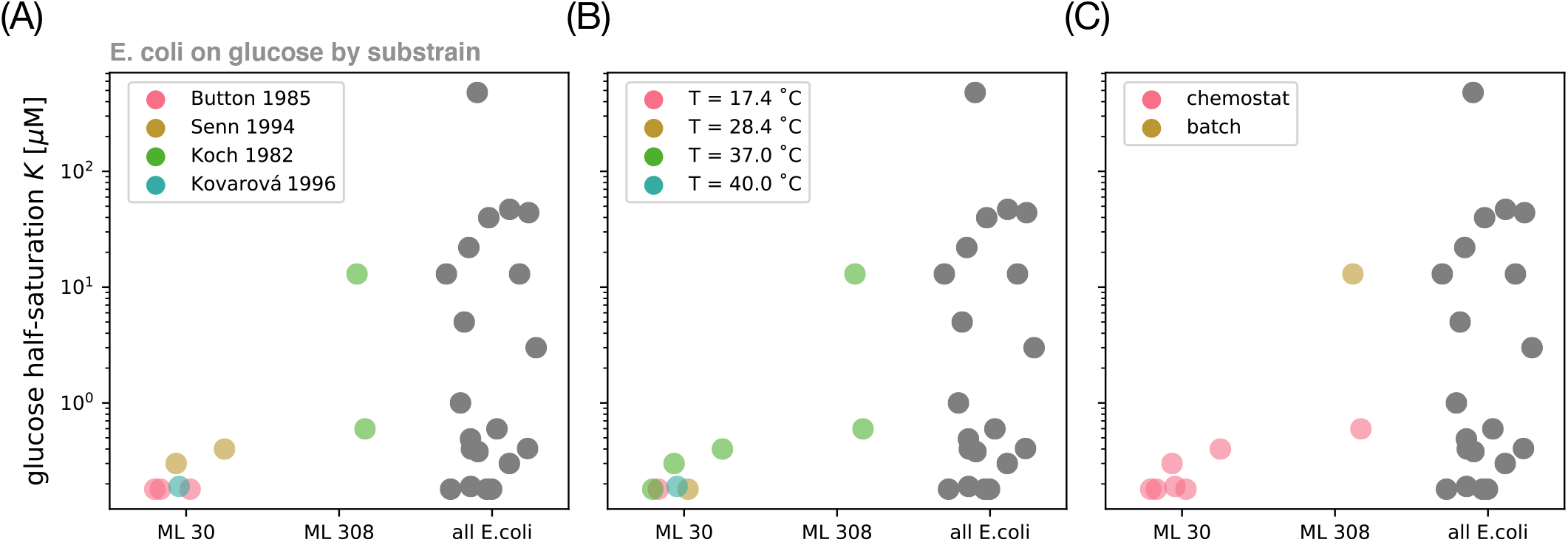
Variation in glucose half-saturation concentrations by experiment type and substrain label. Subset of data from Fig. 2B for *E. coli* on glucose, with different strains separated. The strains ML 30 and ML 308 were derived from a natural isolate in human feces by Jacques Monod in 1946 and differ in their genes for lactose utilization [87]: the lacI repressor is non-functional in ML 308. We only show substrains with two or more measurements from the data. The three panels show the same data but are colored according to (A) publication, (B) temperature, and (C) experimental method (batch or chemostat).

**FIG. S6.**
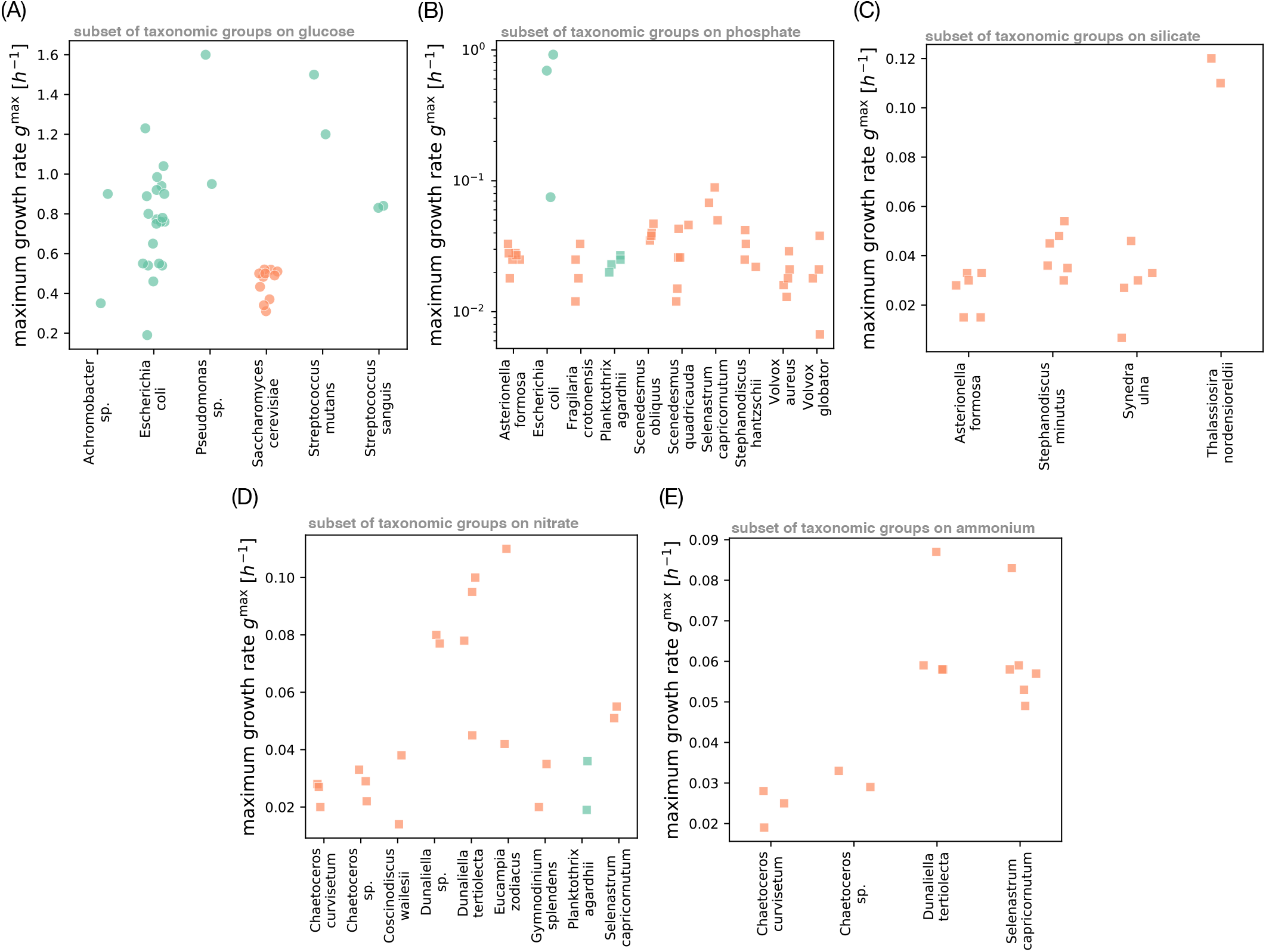
Survey of maximum growth rates in our survey grouped by resource and taxon. (A) Subset of *g*^max^ measurements from Fig. 3A for glucose, grouped by taxon (only those with at least two measurements). Symbols are the same as in Fig. 3A: Color indicates whether the organism is a prokaryote (green) or eukaryote (orange), and shape indicates whether the organism can grow as an autotroph (square) or only as a heterotroph (circle). We use the taxonomic identity given in the original publications, where an ending in *sp*. means the isolate is a representative of the genus but was not identified at the species level. (B) Subset of *g*^max^ measurements from Fig. 3A for phosphate, grouped by taxon (with at least three measurements). Note that we use a logarithmic scale on the y-axis, since this comparison includes both heterotroph isolates (circles) and autotroph isolates (squares) which differ by an order of magnitude in their growth rate. (C) Subset of *g*^max^ measurements from Fig. 3A for silicate, grouped by taxon (with at least two measurements). (D) Subset of *g*^max^ measurements from Fig. 3A for nitrate, grouped by taxon (with at least two measurements). (E) Subset of *g*^max^ measurements from Fig. 3A for ammonium, grouped by taxon (with at least two measurements).

**FIG. S7.**
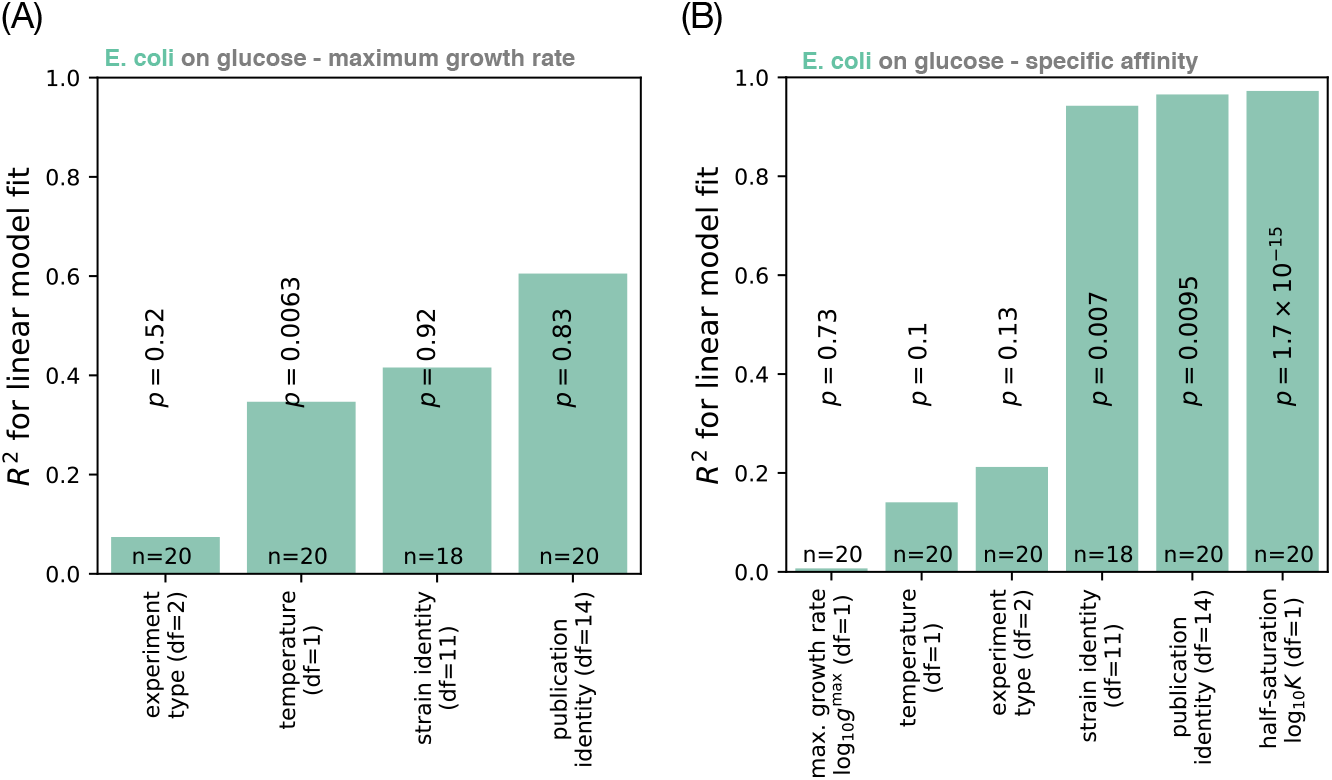
Comparison of technical covariates for maximum growth rate and specific affinity. (A) Linear regression of technical covariates against maximum growth rate on glucose *g*^max^ for all *E. coli* measurements, with degrees of freedom (df), number of data points (*n*), and *p*-values indicated. We follow the same analysis as in Fig. S2B, but using *g*^max^ as the target variable for regression (no log transform). (B) Linear regression of technical covariates against the specific affinity log_10_(*g*^max^/*K*) on glucose for all *E. coli*. The set of underlying isolates is identical to panel A. Here we use the log-transformed maximum growth rate log_10_ *g*^max^ as a predictor, to compare the contributions of variation in log_10_ *g*^max^ and variation in log_10_ *K* to the total variation in log_10_(*g*^max^/*K*). The fraction of variation *R*^2^ explained by log_10_ *g*^max^ is too small to be visible.

**FIG. S8.**
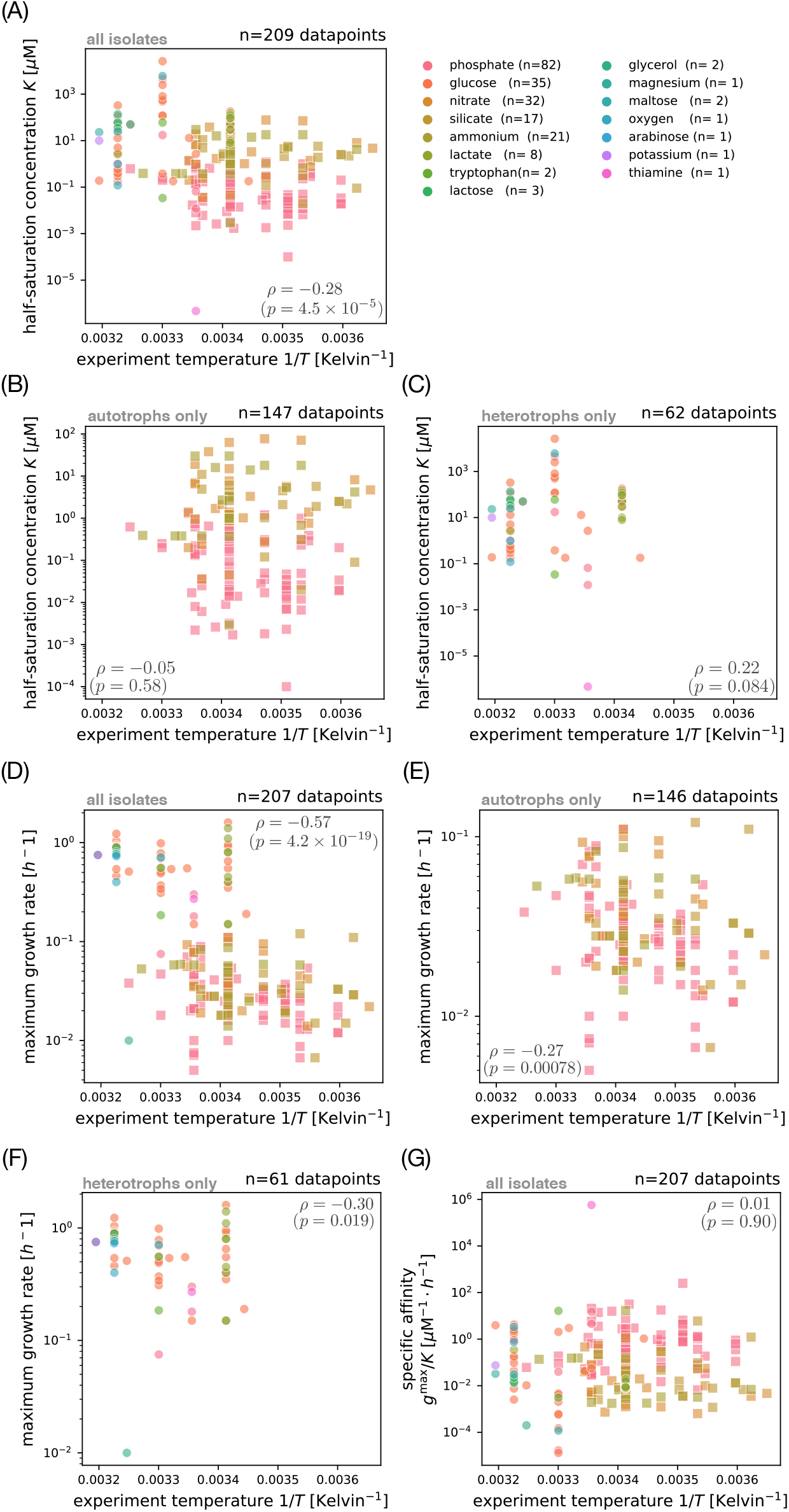
Covariation of Monod growth traits with experiment temperature. (A) Covariation of the half-saturation concentration *K* with the experiment temperature reported in the original publication. Some publications in our survey did not report temperature, so this plot has fewer data points than the full dataset (compare Fig. 2A). We compute the Spearman rank correlation *ρ* and *p*-value across all resources. Colors indicate the limiting resource, with the number of measurements *n* in parentheses. Marker shape separates isolates with an autotroph lifestyle (squares) from heterotrophs (circles). (B) Covariation of the half-saturation concentration *K* with experiment temperature for all autotrophs (subset of points from panel A). (C) Covariation of the half-saturation concentration *K* with experiment temperature for all heterotrophs (subset of points from panel A). (D) Covariation of the maximum growth rate *g*^max^ with experiment temperature. The data shown is less than in panel A, since some publications did not report maximum growth rate. (E) Covariation of the maximum growth rate *g*^max^ with experiment temperature for all autotrophs (subset of points from panel D). (F) Covariation of the maximum growth rate *g*^max^ with experiment temperature for all heterotrophs (subset of points from panel D). (F) Covariation of the specific affinity *g*^max^/*K* with experiment temperature. We compute the specific affinity for all isolates with maximum growth rate in panel D.

**FIG. S9.**
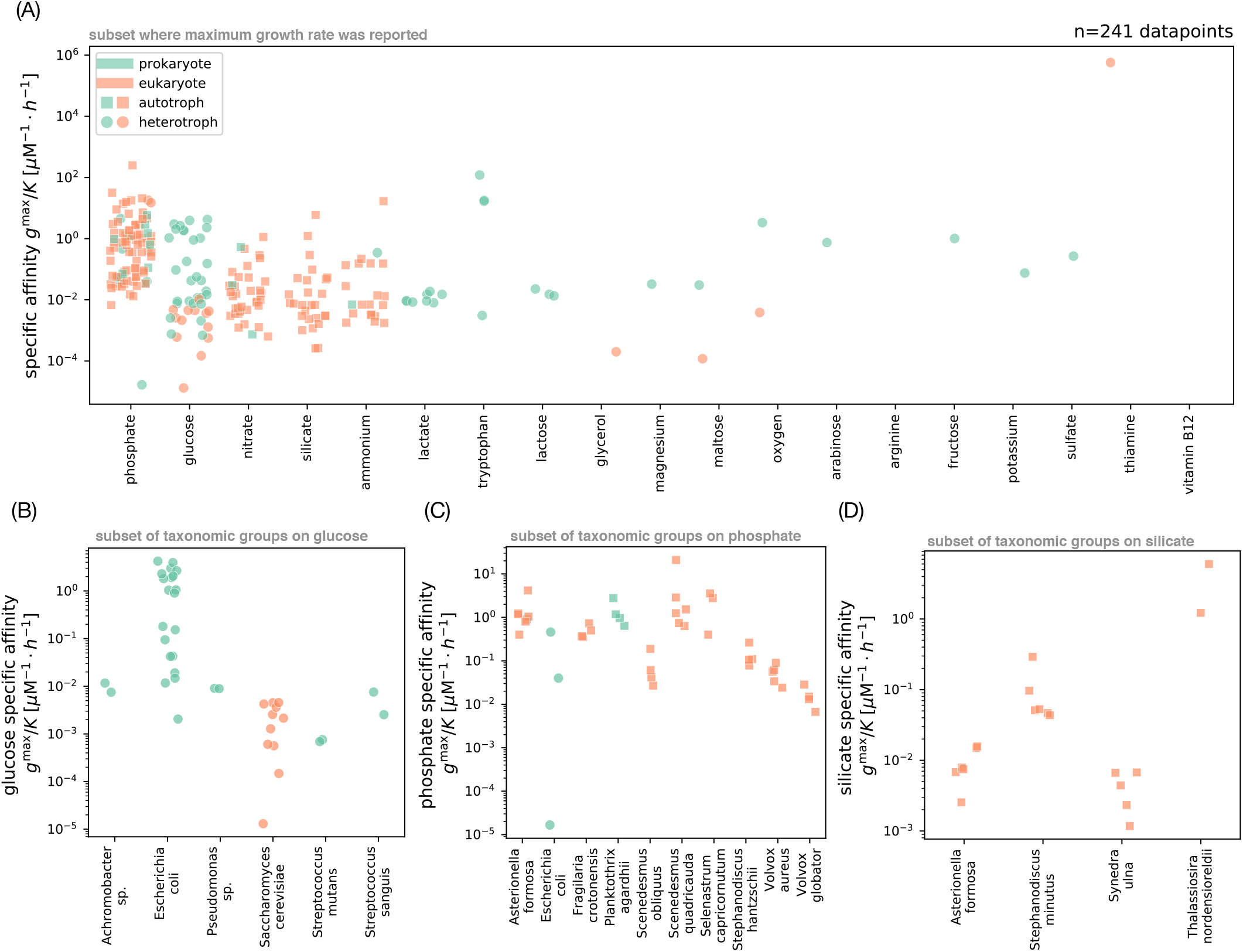
Survey of specific affinities. (A) Variation in specific affinity *a* = *g*^max^/*K* for the microbial isolates in our survey. For each isolate, we compute the trait value from the maximum growth rate *g*^max^ (Fig. 3A) and half-saturation concentration *K* (Fig. 2A). Each point represents a different measurement; color indicates whether the organism is a prokaryote (green) or eukaryote (orange), and shape indicates whether the organism can grow as an autotroph (square) or only as a heterotroph (circle). The set of isolates shown here is fewer than in the total dataset, since some publications only reported the half-saturation concentration *K* and not the maximum growth rate *g*^max^. (B) Subset of *K* measurements from panel A for glucose, grouped by taxon (only those with at least two measurements). We use the taxonomic identity given in the original publications, where an ending in *sp*. means the isolate is a representative of the genus but was not identified at the species level. Symbols are the same as in panel A. (C) Subset of *K* measurements from panel A for phosphate, grouped by taxon (with at least three measurements). (D) Subset for silicate, grouped by taxon (with at least two measurements). Compare also additional plots with *g*^max^/*K* measurements for nitrate (Fig. S4C) and ammonium (Fig. S4D).

**FIG. S10.**
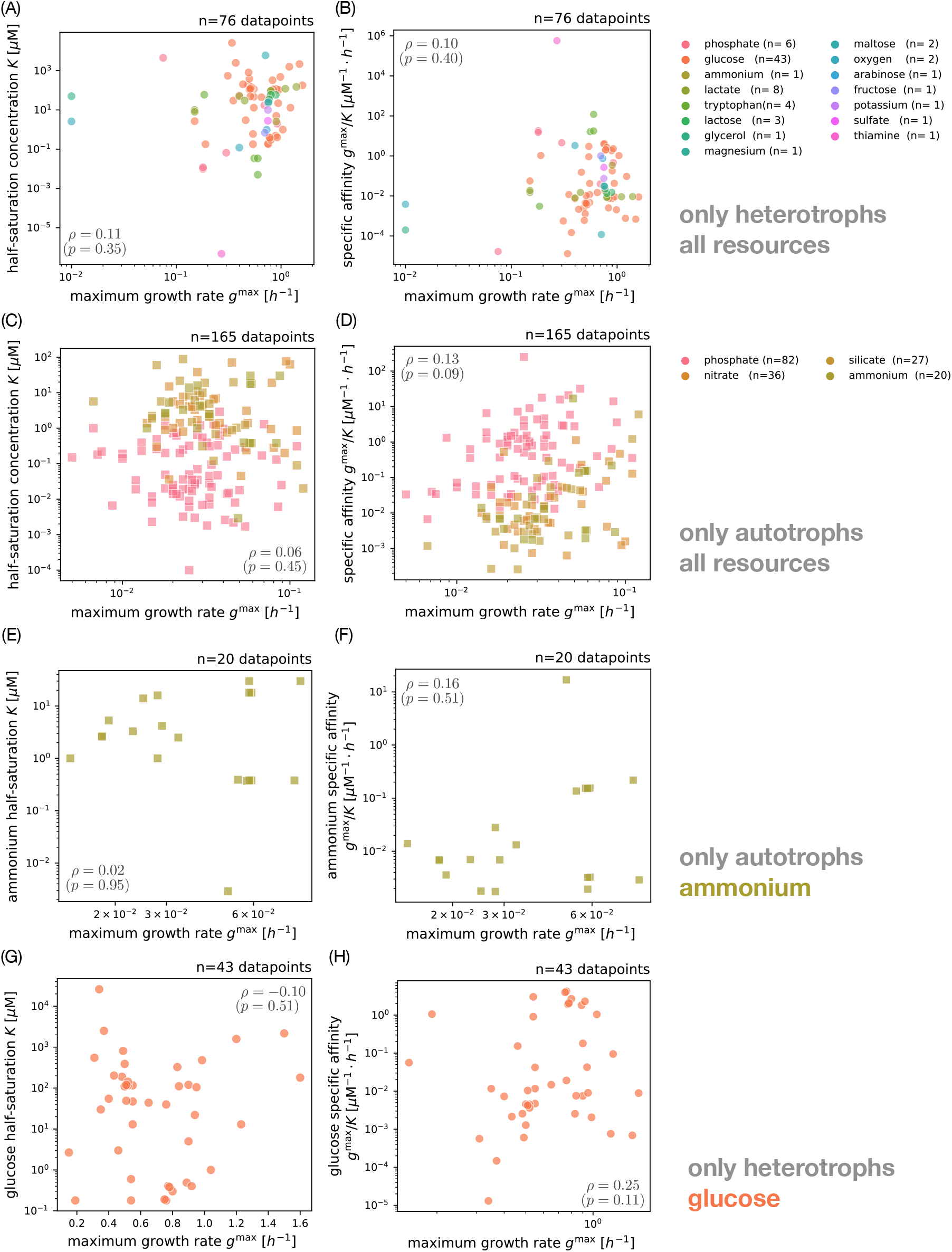
Covariation of Monod growth traits for autotroph and heterotroph isolates. (A) Covariation of half-saturation concentration *K* with maximum growth rate *g*^max^ for all heterotrophs (subset of points from Fig. 3B). We compute the Spearman rank correlation *ρ* and *p*-value across all resources. Colors indicate the limiting resource, with the number of measurements *n* in parentheses. (B) Covariation of specific affinity *g*^max^/*K* with *g*^max^ for all heterotrophs (subset from Fig. S11A). (C) Covariation of half-saturation concentration with maximum growth rate for all autotrophs (subset from Fig. 3B). (D) Covariation of specific affinity with maximum growth rate for all autotrophs (subset from Fig. S11A). (E) Covariation of half-saturation concentration with maximum growth rate for ammonium only (subset from panel C). See Fig. 3C–E for phosphate, nitrate, and silicate. (F) Covariation of specific affinity with maximum growth rate for ammonium only (subset from panel D). See Fig. S11B–D for phosphate, nitrate, and silicate. (G) Covariation of half-saturation concentration with maximum growth rate for glucose only (subset from panel A). See Fig. 3F for covariation within species. (H) Covariation of specific affinity with maximum growth rate for glucose only (subset from panel B). See Fig. S11D for covariation within species.

**FIG. S11.**
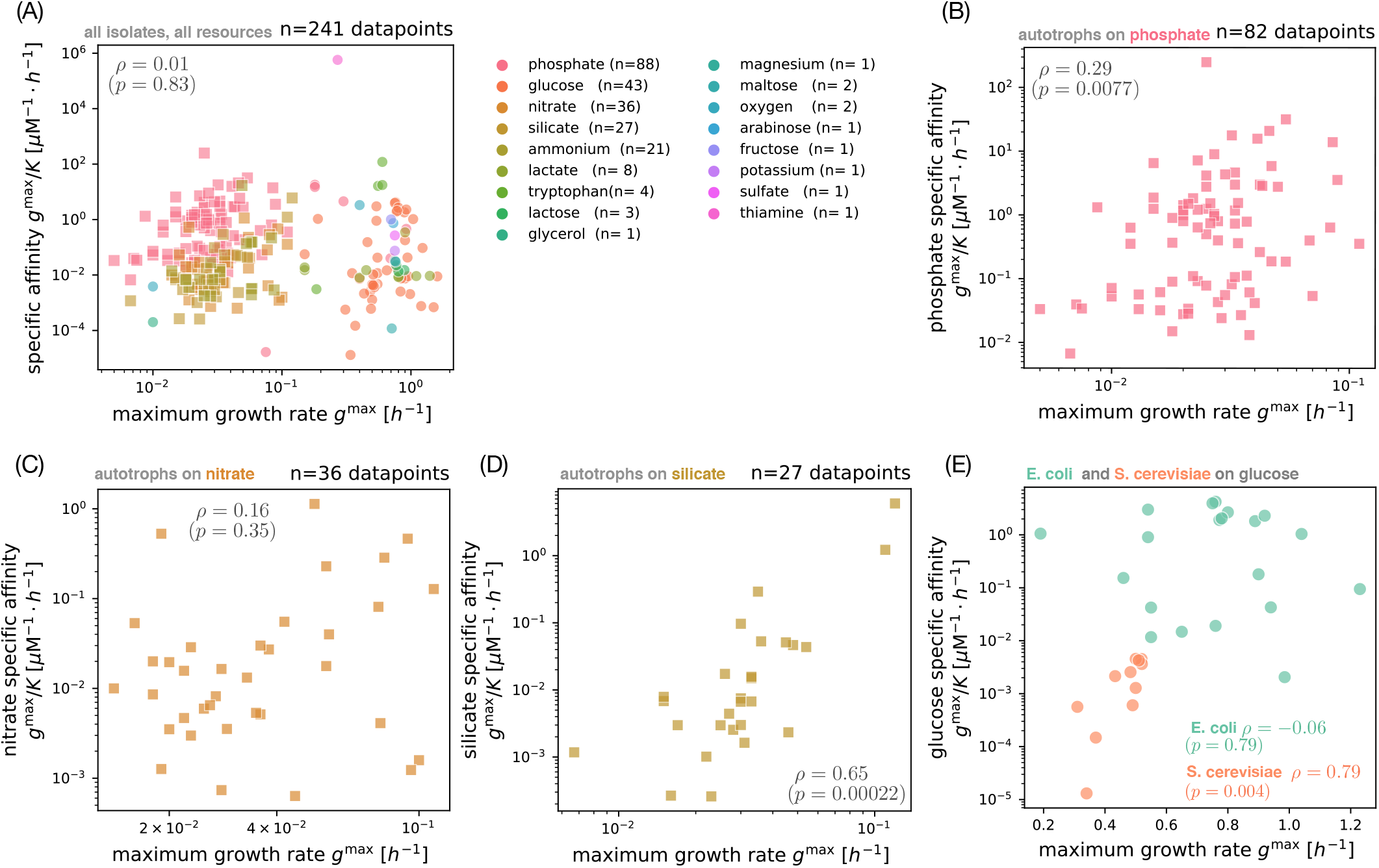
Covariation between maximum growth rate and specific affinity by resource. (A) Covariation of maximum growth rate *g*^max^ and specific affinity *g*^max^/*K* across all resources and isolates (from Fig. S9A). Marker shapes distinguish autotrophs (squares) from heterotrophs (circles); colors indicates the limiting resource, with the number of measurements *n* given in parentheses. We compute the Spearman rank correlation *ρ* and *p*-value across the pooled set of isolates. (B) Subset of measurements from panel A for phosphate (only autotroph isolates shown). (C) Subset of measurements from panel A for nitrate. (D) Subset of measurements from panel A for silicate. (E) Covariation between maximum growth rate *g*^max^ and glucose specific affinity *g*^max^/*K* for measurements of *E. coli* (green) and *S. cerevisiae* (orange), with Spearman rank correlations *ρ* and *p*-values by species.

**FIG. S12.**
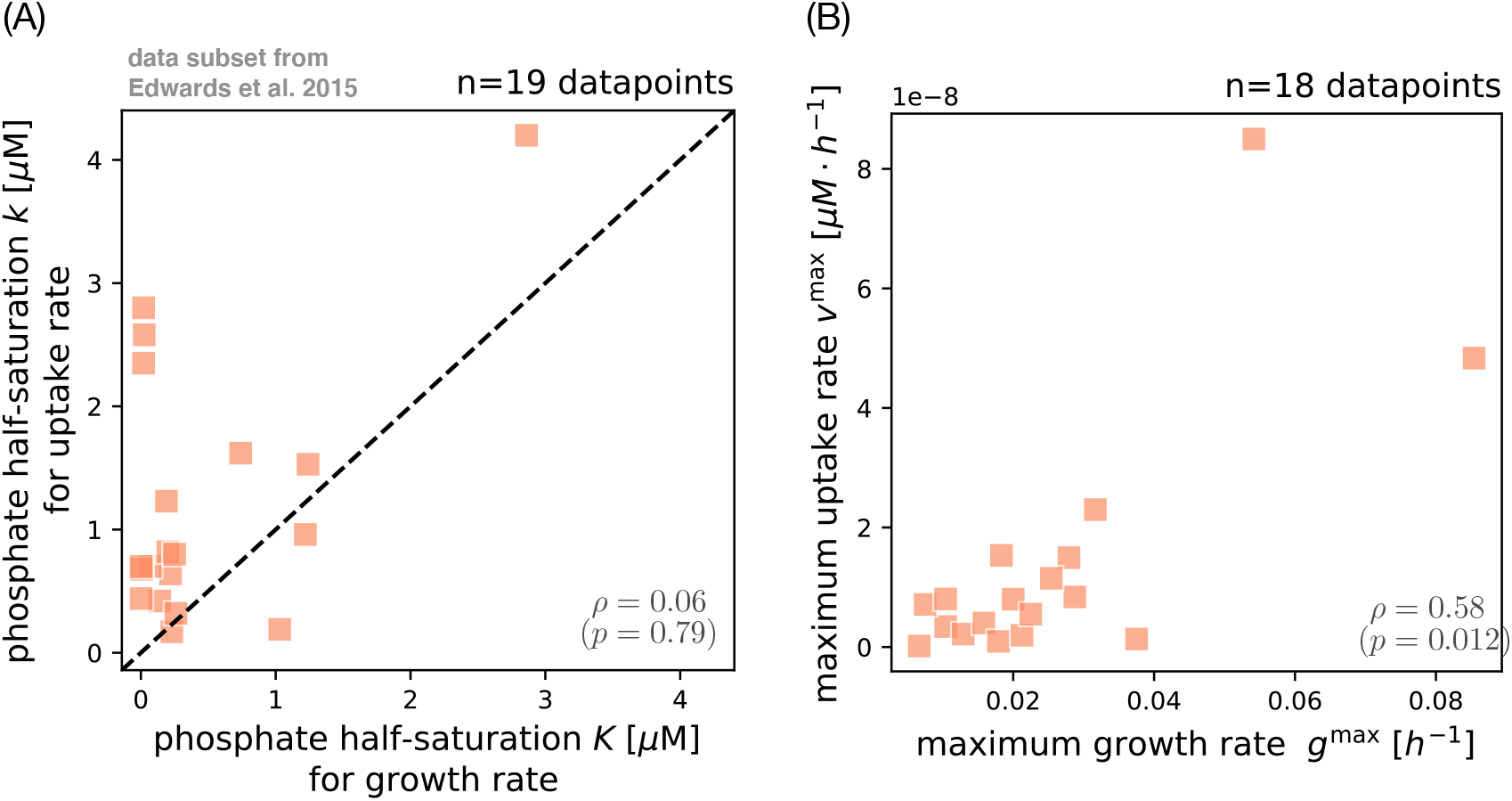
Covariation between uptake and growth rate parameters for phosphate based on the phytoplankton trait database by Edwards et al. [44]. (A) Covariation between the half-saturation concentration *k* for uptake rate and the half-saturation concentration *K* for growth rate (Eq. (1)). The dashed diagonal line indicates perfect agreement (*x* = *y*), and we calculate the Spearman rank correlation *ρ* with *p*-value. We show all data points from Edwards et al. [44] which included half-saturation concentrations for uptake and growth rate. These data points are for phosphate as the limiting resource. (B) Covariation between maximum uptake rate *v*^max^ in the Michaelis-Menten model and the maximum growth rate *g*^max^ in the Monod model, with Spearman rank correlation *ρ* and *p*-value. The data shown here corresponds to the same measurements as in panel A but with one fewer data point, since one isolate lacked the measurement for maximum growth rate. Color and marker shape are equivalent to Fig. 2A and indicate that the subset of data shown here includes only eukaryotic organisms (orange fill) capable of autotrophy (square shape).

**FIG. S13.**
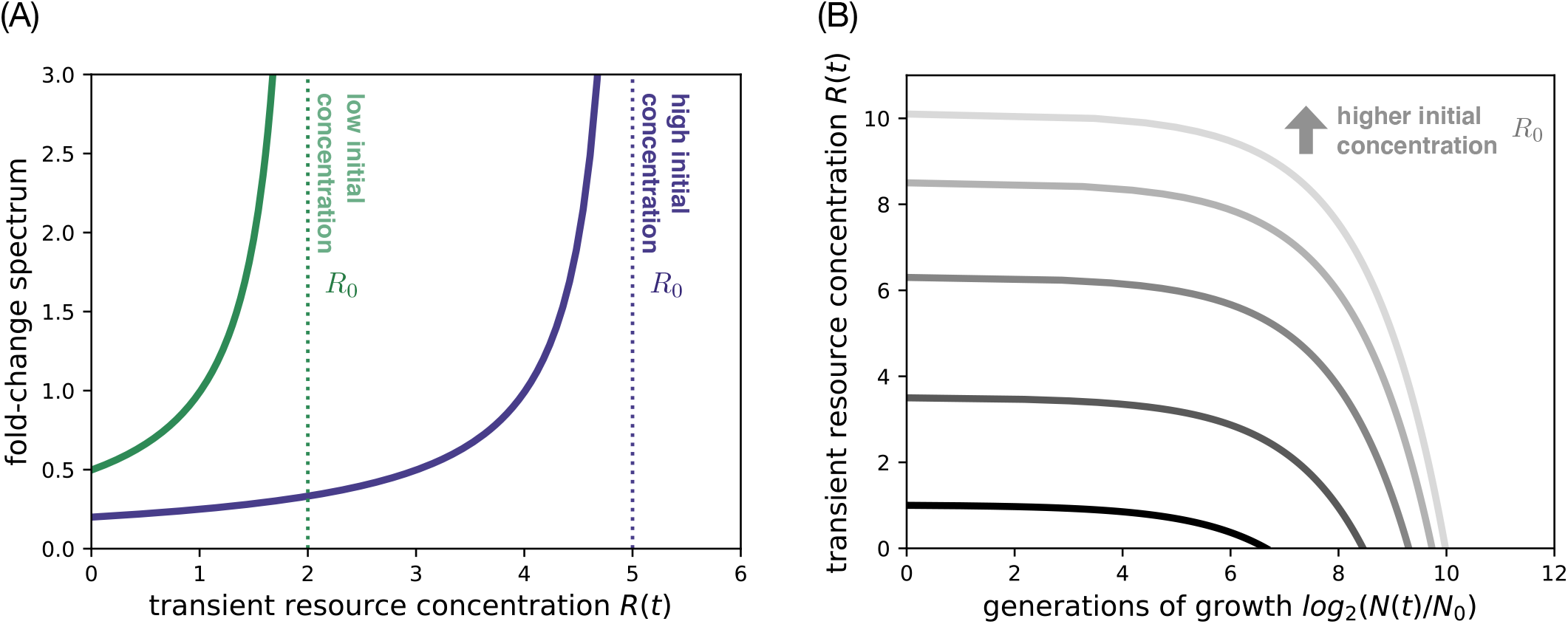
Selection within a batch growth cycle. (A) The fold-change spectrum (thick lines) throughout the growth cycle for high and low initial concentration *R*_0_ (dotted lines). Curves are computed from the weight term in Eq. (S42) with effective biomass yield *Ȳ* = 1 and *N*_0_ = 0.01. (B) The transient resource concentration, starting from different initial concentrations, versus generations of biomass growth (gray lines). The lowest line (*R*_0_ = 1) corresponds to the resource trajectory for the selection scenario used for the phase diagram in Fig. S15. The transient resources are converted into generations using the equations for resource consumption, assuming an identical biomass yield *Y* = 1 for both strains and initial biomass *N*_0_ = 0.01.

**FIG. S14.**
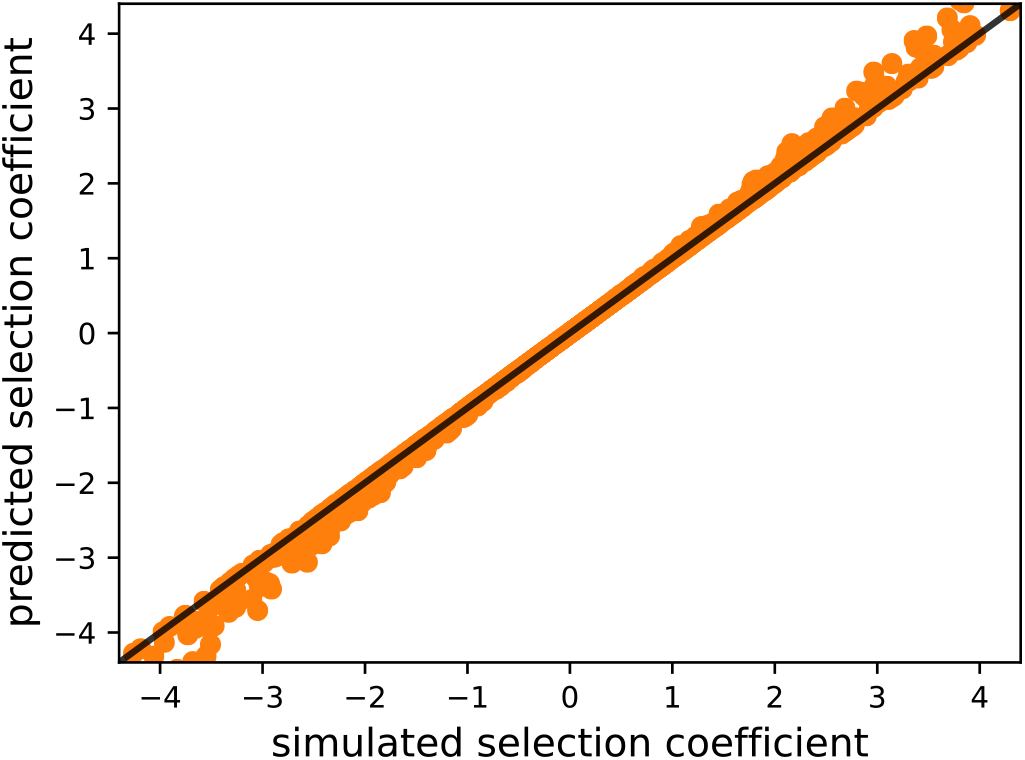
Test of the selection coefficient approximation. The predicted selection coefficient across a sample of wild-type and mutant strains, compared to the selection coefficient (Supplementary Information Sec. S6) from simulation of the differential equations (Supplementary Information Secs. S3 and S4). The black diagonal line indicates perfect agreement between simulation and prediction. We draw the wild-type traits over four orders of magnitude and sample relative mutant effects on maximum growth rate, half-saturation concentration, and biomass yield from a cubic region in trait space: [−0.5, 0.5]^3^. Each strain pair is systematically evaluated at different initial frequencies *x* = 0.01, 0.5, 0.99 using the general Eq. (S47) and contributes three data points. Without loss of generality, we fix the initial biomass to *N*_0_ = 0.01 and initial resource concentration to *R*_0_ = 1. The trait values for the half-saturation concentration *K* span two orders of magnitude around this concentration such that we cover both limiting scenarios with dominant selection on maximum growth rate (*R*_0_ ≫ *K*) and half-saturation concentration (*R*_0_ ≪ *K*).

**FIG. S15.**
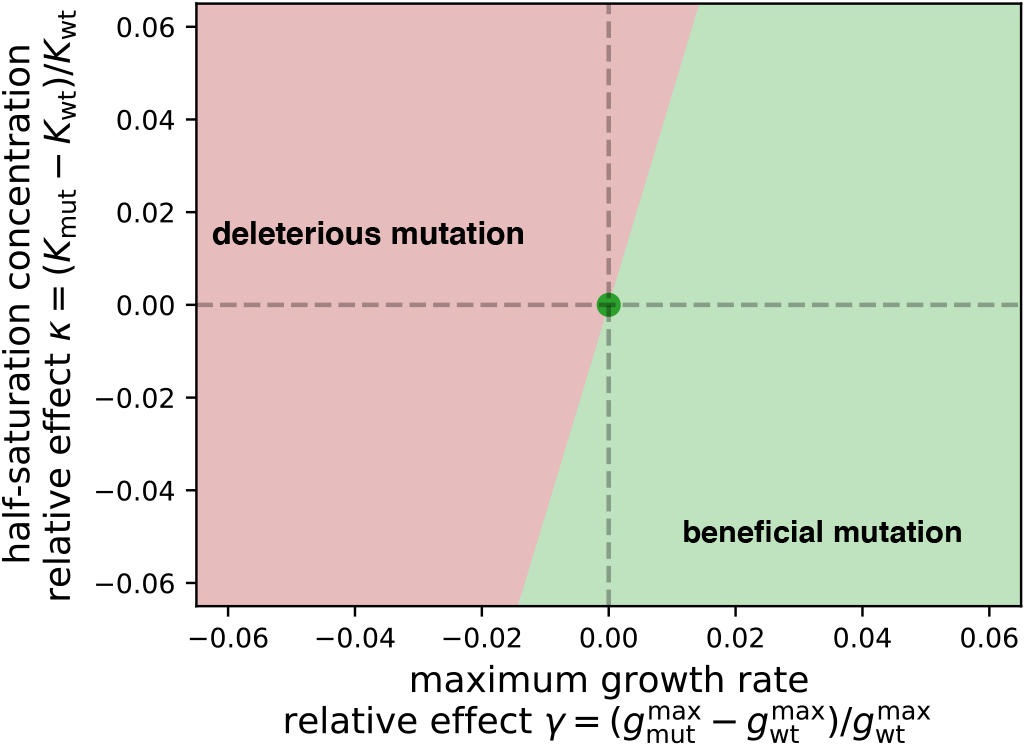
Diagram of selection across mutation effects under batch growth. The space of mutation effects on maximum growth rate 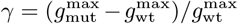 and half-saturation concentration *κ* = (*K*_mut_ – *K*_wt_)/*K*_wt_ relative to a wild-type strain (central dot), with green marking the space of mutations that are overall beneficial (*s* > 0) and red marking mutations that are overall deleterious (*s* < 0) according to Eq. (S47).

**FIG. S16.**
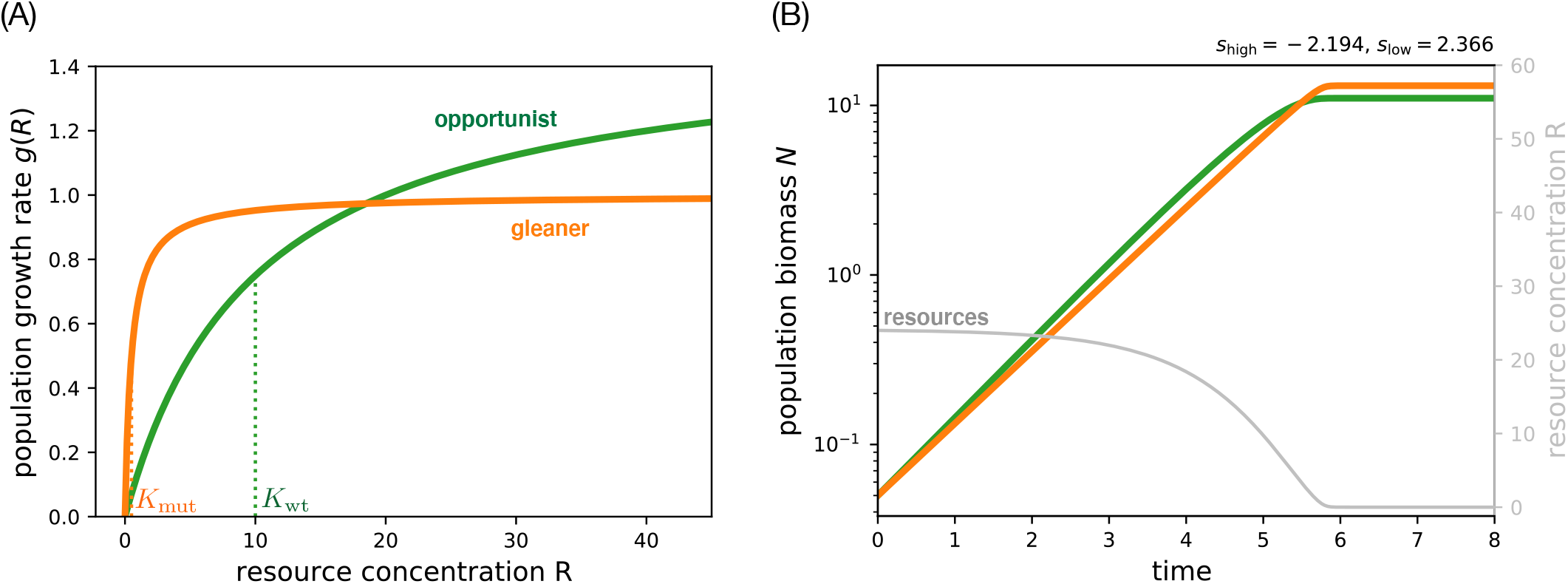
The gleaner and opportunist strategies in the Monod growth model. (A) The growth rate *g*(*R*) as a function of the external resource concentration *R* for two strains with a tradeoff. The opportunist strain (green) has a higher maximum growth rate *g*^max^ = 1.5 compared to the gleaner strain (orange) with *g*^max^ = 1. But the gleaner has the growth rate advantage at low concentrations due to a smaller half-saturation concentration *K* = 0.5 (orange dotted line) relative to the opportunist with *K* = 10 (green dotted line). (B) A single growth cycle for the gleaner and opportunist strain pair from panel A in competition. We simulate the population dynamics according to Eq. (S11), starting from an initial mutant frequency *x* = 0.5 and total initial biomass *N*_0_ = 0.01. On a separate axis, the transient resource concentration *R* (gray line, initial value *R*_0_ = 24) and in the panel title, the components of selection as computed from Eq. (S47).

**FIG. S17.**
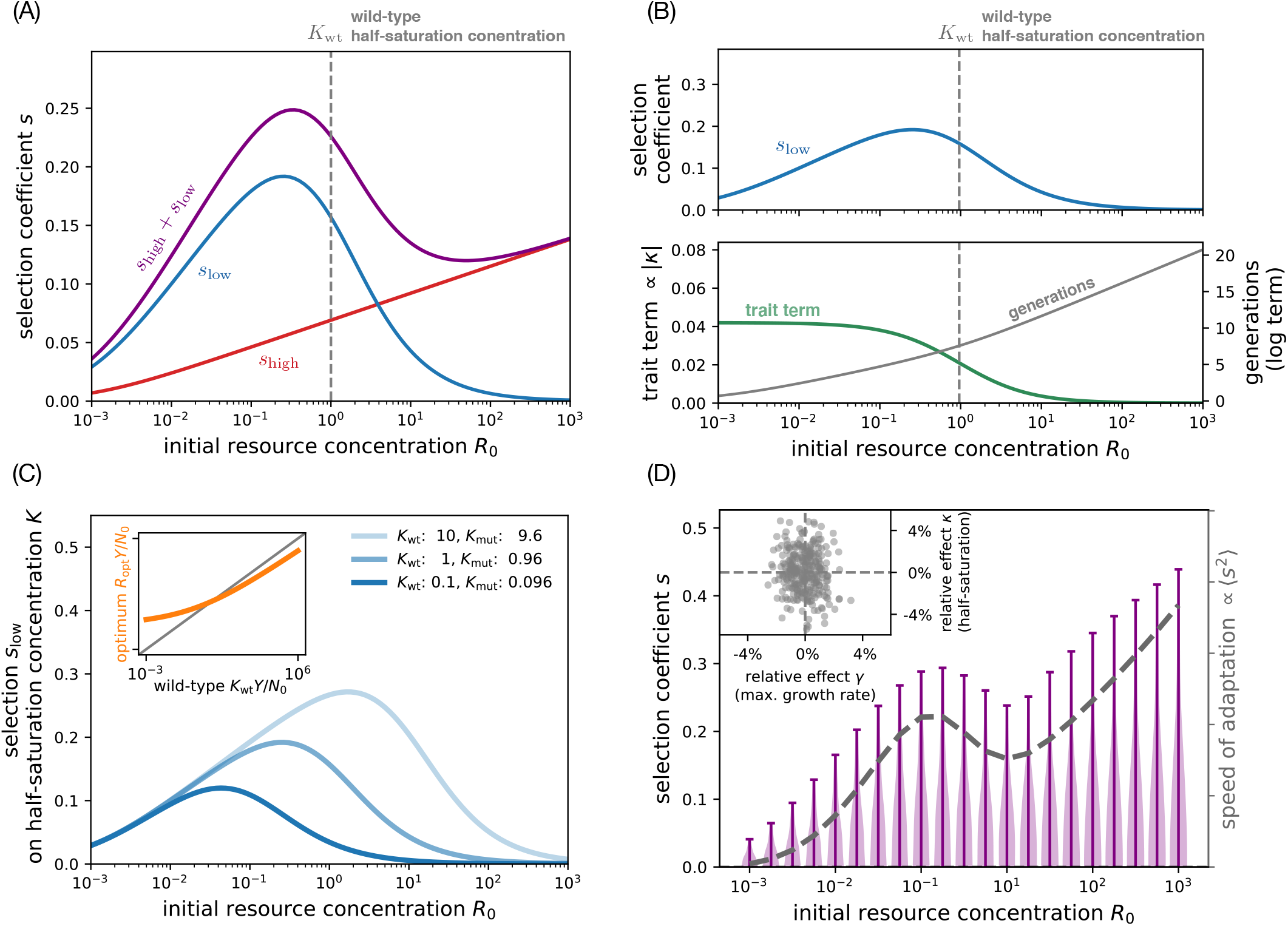
Selection dependence on resource concentration under fixed-bottleneck batch dynamics. (A) Dependence on external resource concentration *R*_0_ of the total selection coefficient (purple) and its two components at high (*s*_high_, red) and low (*s*_low_, blue) resource concentrations under a fixed bottleneck (Eq. (S47)). (B) Selection *s*_low_ on growth at low resource concentrations (top panel) decomposed into two constituent factors (bottom panel) at fixed bottleneck biomass (*N*_0_ = 10^−3^). The top panel is the same as panel A for the approximate selection coefficient *s*_low_; in the bottom panel, the two factors that constitute *s*_low_ are the trait factor from Eq. (S60) in green and the log term from Eq. (S61) in gray. The log term is related but not identical to the number of generations in the growth cycle. Panels A and B are based on an example mutation with relative effects *γ* = 0.01 on maximum growth rate and *κ* = −0.04 on half-saturation concentration over the wild-type traits *g*_wt_ = 1 and *K*_wt_ = 1. (C) Selection on the half-saturation concentration *K* as a function of resource concentration *R*_0_ for three different values of *K* (different shades of blue). The inset shows a numerical calculation (orange points) of the optimal resource concentration *R*_opt_ that maximizes selection on *K* as a function of the wild-type half-saturation *K*_wt_; the gray line is the identity. Parameters are the same as in panels A and B, but we include two alternative wild-type half-saturation concen-trations *K*_wt_ = 10 (lightest blue) and *K*_wt_ = 0.1 (darkest blue). (D) The distribution of beneficial selection coefficients (purple) as a function of initial resource concentration *R*_0_, with the variance (which is proportional to the speed of adaptation) shown as the dashed gray line and plotted against the right axis. The inset shows the underlying sample of mutations according to their relative effects on maximum growth rate *γ* and half-saturation concentration *κ*. We sample the effects of mutations from independent Gaussian distributions for *γ* (mean *μ* = 0, s.d. *σ* = 0.01) and *κ* (mean *μ* = 0, s.d. *σ* = 0.02). All panels assume initial population biomass *N*_0_ = 0.001, initial mutant frequency *x* = 0.01, and equal yields for mutant and wild-type.

**FIG. S18.**
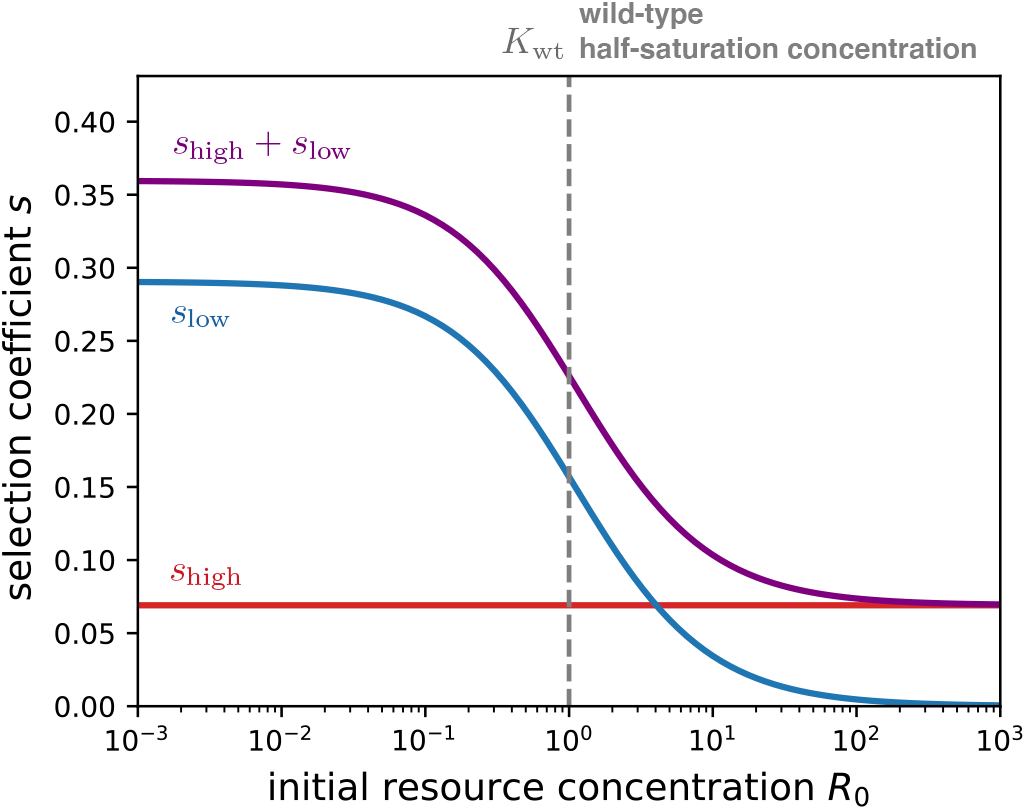
Selection dependence on resource concentration under fixed-dilution batch dynamics. Same as Fig. S17A, but for fixed-dilution batch dynamics with *D* = 1000.

**FIG. S19.**
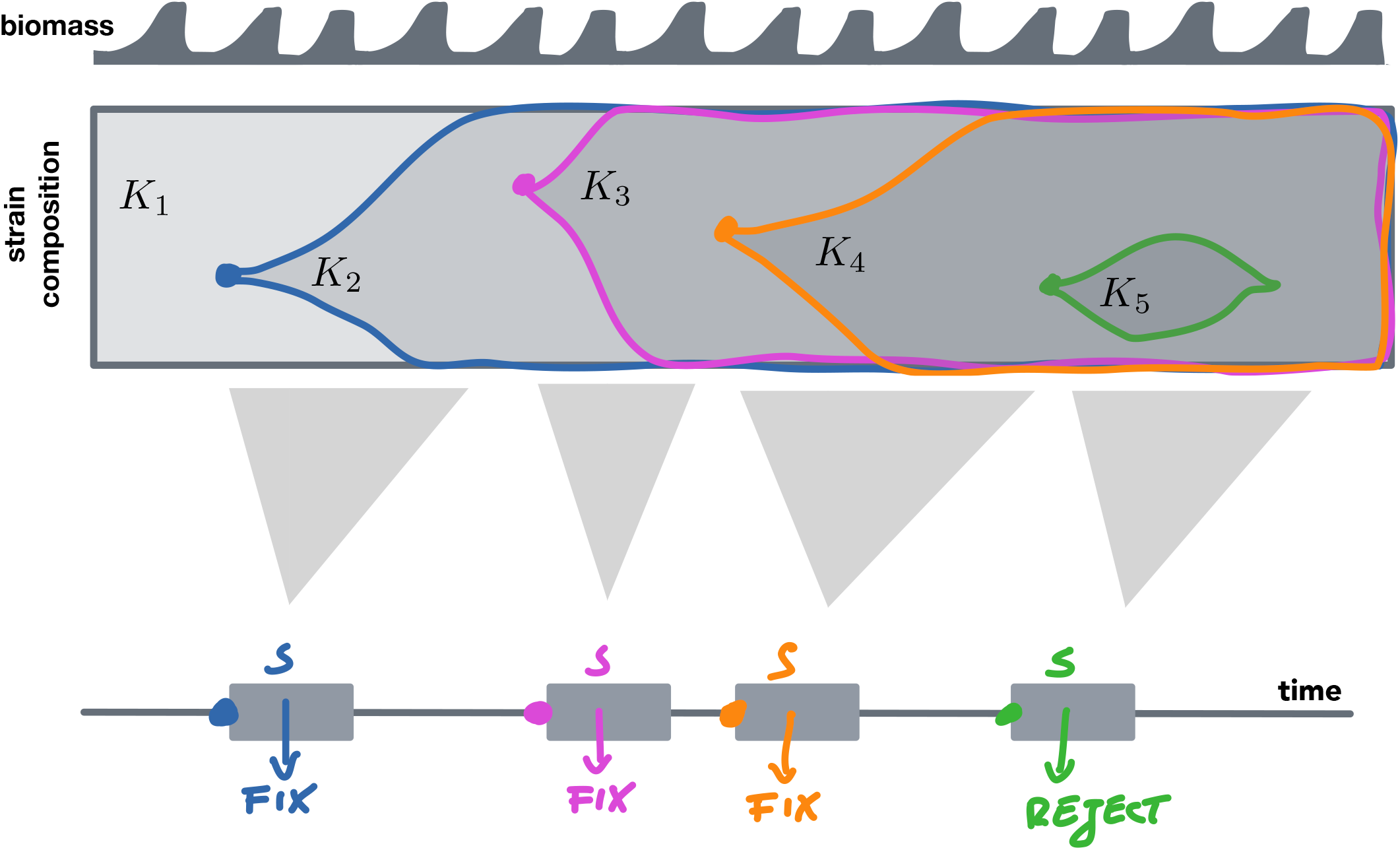
Schematic of evolutionary dynamics in the strong-selection weak-mutation (SSWM) regime. The top panel shows a schematic of the population biomass undergoing cycles of batch dynamics with serial transfers. The middle panel shows the genetic composition of the population. The population begins with a half-saturation concentration *K*_1_. Then a mutation arises with a different half-saturation *K*_2_ (blue), which increases in frequency until it fixes. Then another mutation with a half-saturation value *K*_3_ arises (magenta), and the process continues. The bottom panel shows a simplified algorithm for this process that we use in our simulations (Supplementary Information Sec. S11), where mutations are determined to fix or go extinct one at a time based on their selection coefficients, without explicitly simulating their intermediate frequency dynamics.

**FIG. S20.**
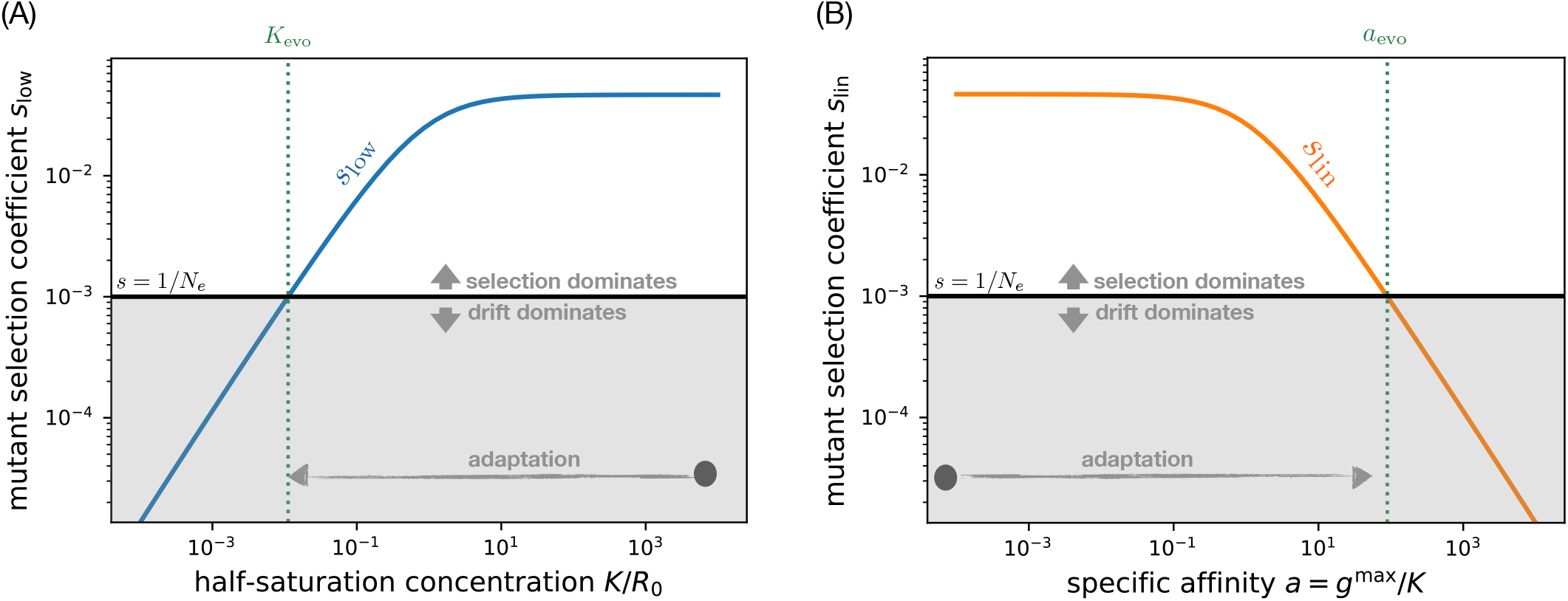
Selection-drift balance under batch dynamics. (A) Selection *s*_low_ on the half-saturation concentration *K* as a function of the current wild-type trait value in the population (blue line, Eq. (S47)), with mutation effect *κ*_max_ = −10^−2^. The horizontal black line marks the strength of genetic drift; its intersection with the selection coefficient defines the value of *K*_evo_ at which selection-drift balance occurs (vertical dotted line; Eq. (3)). Above this point, selection is stronger than genetic drift, and so the half-saturation concentration will adapt downward until it reaches that point. (B) Selection on the specific affinity *a* = *g*^max^/*K* as a function of the current wild-type trait value in the population (orange line, Eq. (S52c)) assuming a relative mutation effect *α* = 10^−2^ that acts directly on the specific affinity instead of on the half-saturation concentration. For the specific affinity, adaptation means the trait value increases. Similar to panel A, the intersection of the selection coefficient with the black line (strength of genetic drift) defines the evolved trait value *a*_evo_ at selection-drift balance. Parameters are identical in both panels with *R*_0_ = 1, *N*_0_ = 0.01, *N*_e_ = 10^3^, and *x* = 0.001. We set mutant and wild-type to equal maximum growth rates and equal yields. This plot is based on fixed bottleneck biomass *N*_0_, but we observe similar dependences for fixed dilution factor *D*. In that case, we rewrite Eq. (S47) (resp. Eq. (S52c)) in terms of *D* (replacing *N*_0_ using Eq. (S21)) and see that the selection coefficient depends on the wild-type trait *K* with the same functional form.

**FIG. S21.**
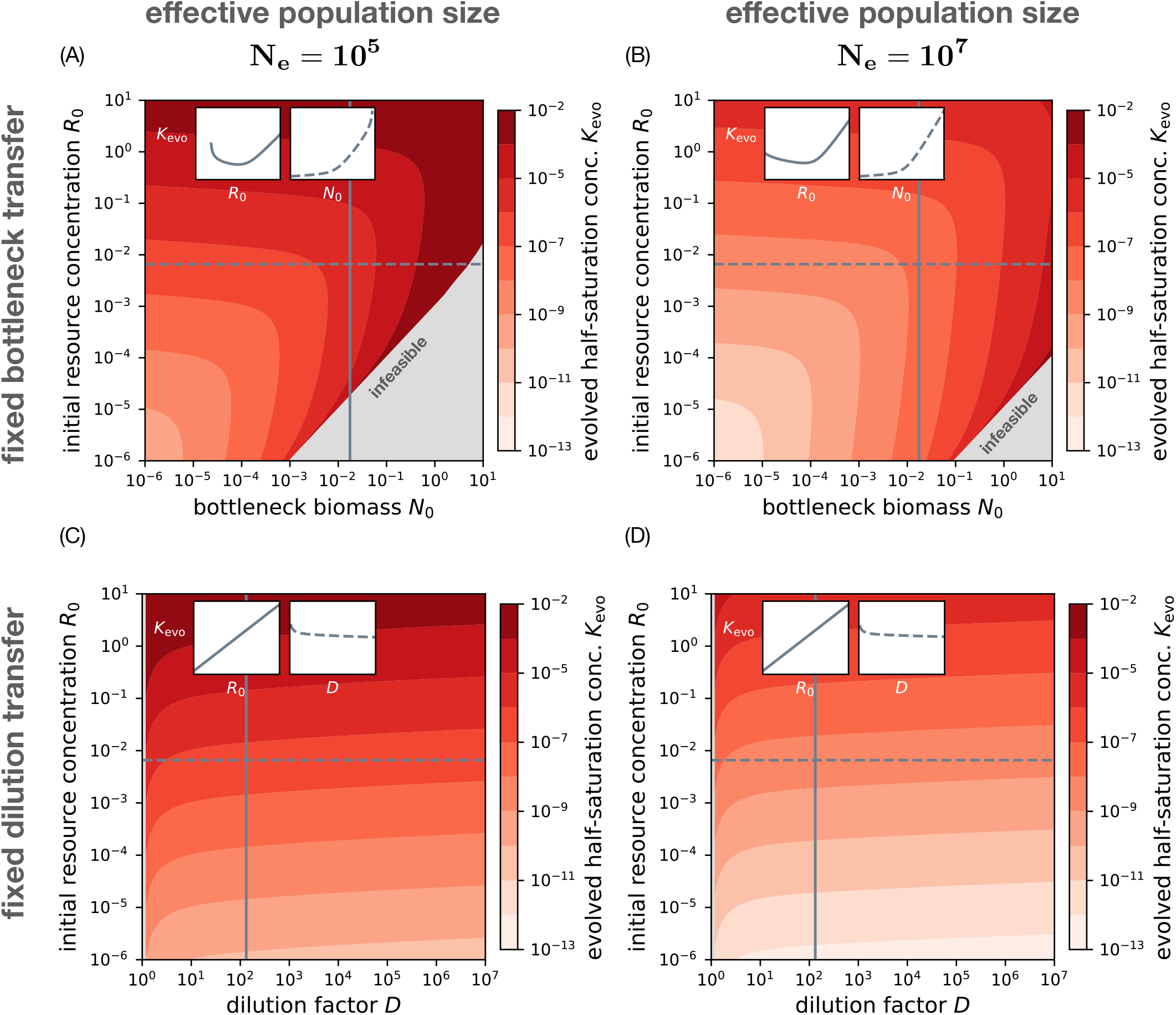
The evolved half-saturation concentration as a function of experimental parameters with independent genetic drift. (A) We numerically solve for the evolved half-saturation concentration *K*_evo_ under selection-drift balance (using Eqs. (2) and (3)) as a function of the fixed-bottleneck biomass *N*_0_ and initial resource concentration *R*_0_, where the effective population size *N*_e_ = 10^5^ is an independent parameter. Where the selection-drift balance condition is infeasible (gray area), the half-saturation concentration evolves neutrally without steady state. The insets show cross-sections along initial resource concentration *R*_0_ (solid line) and bottleneck biomass *N*_0_ (dashed line). (B) Same as panel A, but for a larger effective population size *N*_e_ = 10^7^. (C) Same as panel A but for fixed-dilution batch dynamics, with varying D instead of *N*_0_. (D) Same as panel C, but for a larger effective population size *N*_e_ = 10^7^. All panels use identical growth rates *g*^max^ = 1 and biomass yields *Y* = 1 for wild-type and mutant strain with a fixed mutation effect *κ* = 0.01 on the half-saturation concentration. The initial mutant frequency *x* = 1/*N*_e_ is adjusted to the effective population size *N*_e_.

**FIG. S22.**
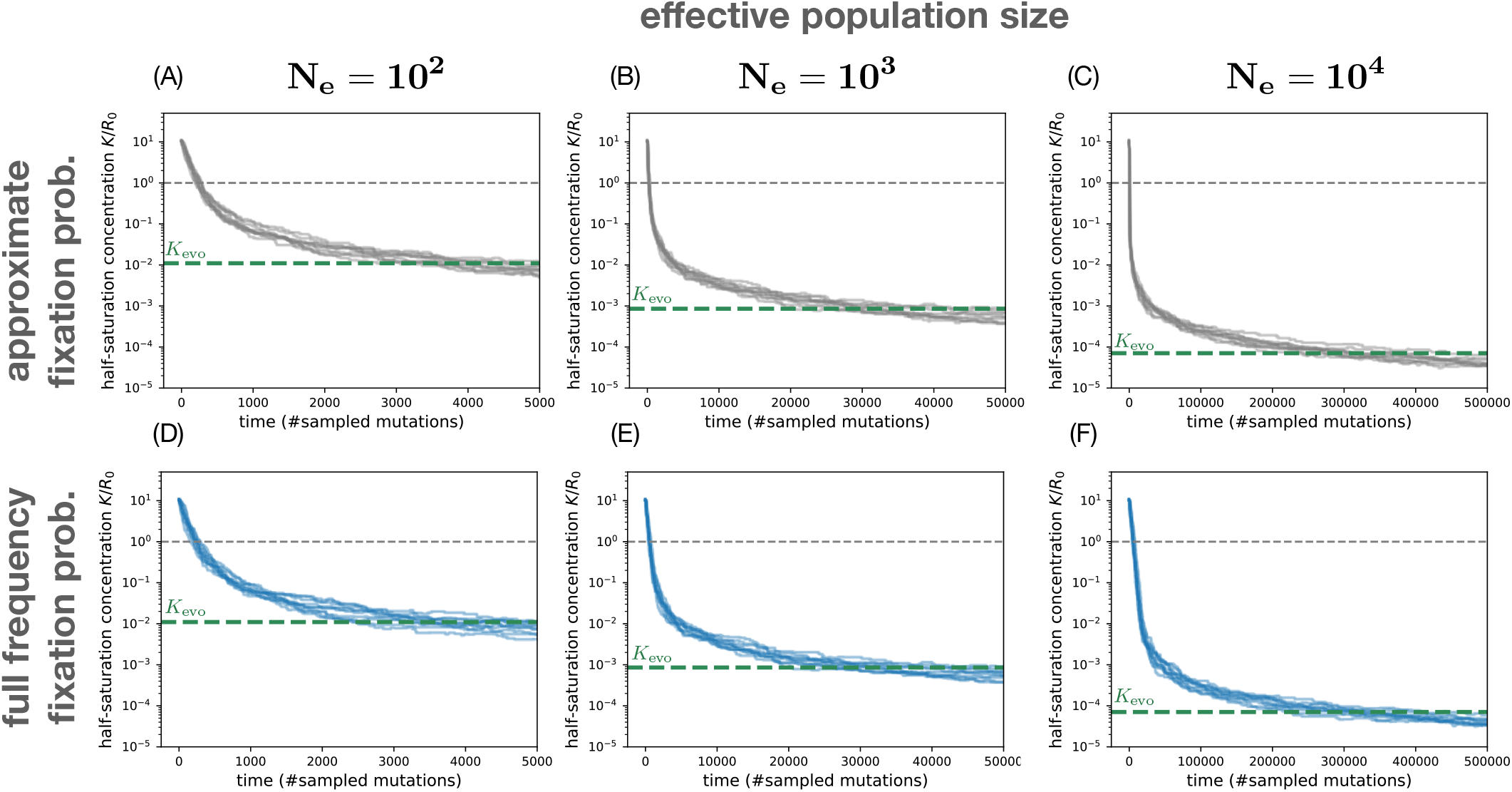
Simulated evolutionary trajectories of the half-saturation concentration under batch dynamics. We simulate the time-course of evolution in the half-saturation concentration under the SSWM regime (Supplementary Information Sec. S11) for different strengths of genetic drift (1/*N*_e_). Each line corresponds to a separate run of the stochastic evolution process. Here mutations are sampled with random effect *κ* = (*K*_mut_ – *K*_wt_)/*K*_wt_ from a uniform distribution in [−0.1, 0.1] and accepted or rejected according to their probability of fixation (Supplementary Information Sec. S11). (A)-(C) In the top row, we use the approximate fixation probability Eq. (S63) which depends only on the selection coefficient at the initial mutant frequency *x* = 1/*N*_e_. (D)–(F) In the bottom row, we use the integrated form of the fixation probability from Eq. (S62) that takes into account the frequency-dependence of the mutant selection coefficient (Eq. (S47)). For each panel, we numerically calculate the half-saturation concentration *K*_evo_ at selection-drift balance (dashed line) using Eqs. (2) and (3). To guide the eye, we also mark the half-saturation concentration *K* = *R*_0_ that matches the environmental concentration (gray line). All panels are based on identical maximum growth rates *g*^max^ = 1 and biomass yields *Y* = 1 for the mutant and wild-type strain such that only the half-saturation concentration evolves. The length of the growth cycle is constant with ≈ 6.6 generations at initial resource concentration *R*_0_ = 1 and fixed-bottleneck biomass *N*_0_ = 0.01.

**FIG. S23.**
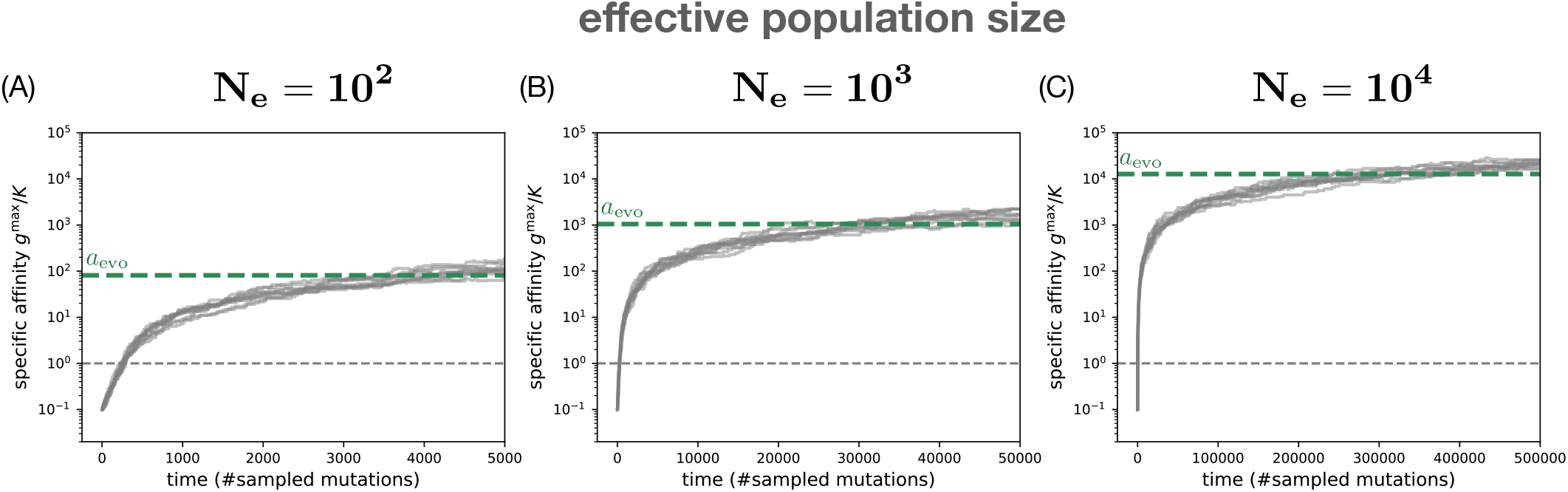
Simulated evolutionary trajectories for the specific affinity under batch dynamics. We simulate the time-course of evolution in the SSWM regime (Supplementary Information Sec. S11) similar to Fig. S22, but assuming that mutations directly affect the specific affinity *a* = *g*^max^/*K* instead of the half-saturation *K* alone. Here mutations are sampled with random effect *α* = (*a*_mut_ – *a*_wt_)/*a*_wt_ from a uniform distribution in [−0.1, 0.1] and accepted or rejected according to their probability of fixation (compare also Sec. S14). Here we use the approximate fixation probability Eq. (S63) which depends on the selection coefficient at the initial mutant frequency *x* = 1/*N*_e_. The panels (A)–(C) only differ in the effective population size *N*_e_ used for the simulation. For each panel, we numerically calculate the specific affinity *a*_evo_ at selection-drift balance (dashed line) using Eqs. (S52c) and (3). All panels are based on identical maximum growth rates *g*^max^ = 1 and biomass yields *Y* = 1 for the mutant and wild-type strain such that only the specific affinity evolves. The length of the growth cycle is constant with ≈ 6.6 generations at initial resource concentration *R*_0_ = 1 and fixed-bottleneck biomass *N*_0_ = 0.01.

**FIG. S24.**
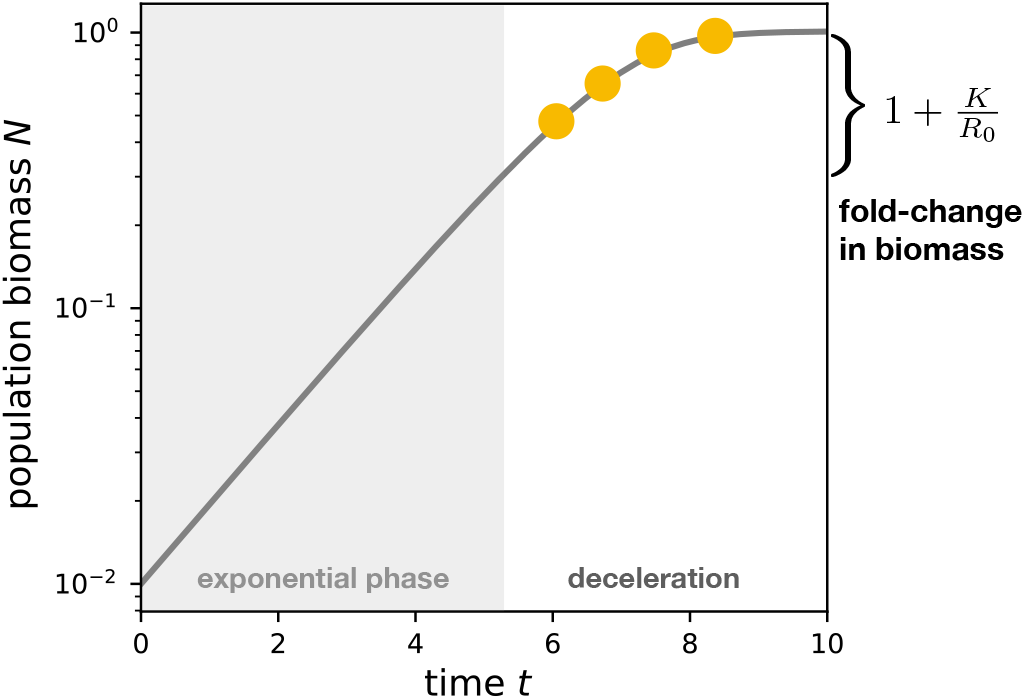
Detecting the half-saturation concentration *K* from time-series data. We use an initial resource concentration *R*_0_ = 10 close to the half-saturation concentration of the wild-type strain (*K*_wt_ = 5; see Fig. 1) to simulate a monoculture growth curve from Eq. (S11) (Supplementary Information Sec. S3). The population leaves steady exponential growth phase (gray area) to enter the deceleration phase (white area). To fit the half-saturation concentration *K*, the time-series must include multiple data points in the deceleration phase (orange dots; Supplementary Information Sec. S15). On the right axis, a bracket marks the fold-change from the onset of deceleration at biomass to the saturation.

**FIG. S25.**
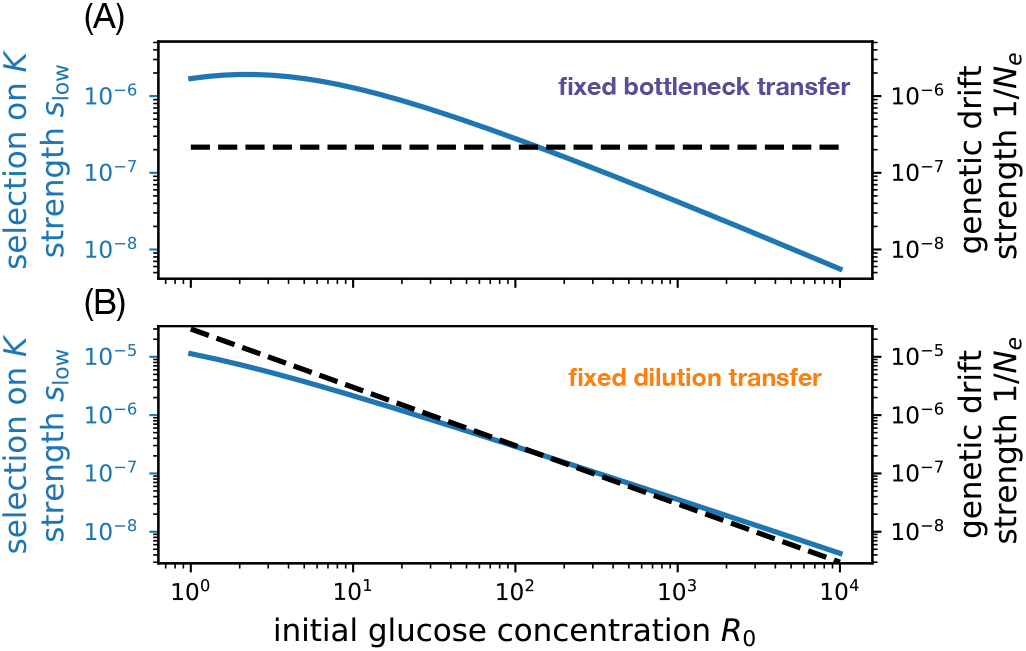
Environmental dependence of selection and genetic drift under batch dynamics. The top panel shows selection *s*_low_ on the half-saturation concentration *K* (blue solid line, left axis) and the strength of genetic drift 1/*N*_e_ (dashed black line, right axis) as functions of the resource concentration *R*_0_ under fixed-bottleneck batch dynamics. In this case, the effective population size is independent of the resource concentration. We use parameters based on the LTEE (same as in Fig. 5C): *N*_0_ = 4.6 × 10^5^ cells/mL and *N*_e_ = *VN*_0_ using culture volume *V* = 10 mL, *g*^max^ = 0.888/h, and *Y* = 3.3 × 10^8^ cells/μmol [23]. The bottom panel shows the same but for fixed-dilution batch dynamics, with *D* = 100; in this case the effective population size is proportional to the resource concentration, and thus the strength of genetic drift decreases with *R*_0_.

**FIG. S26.**
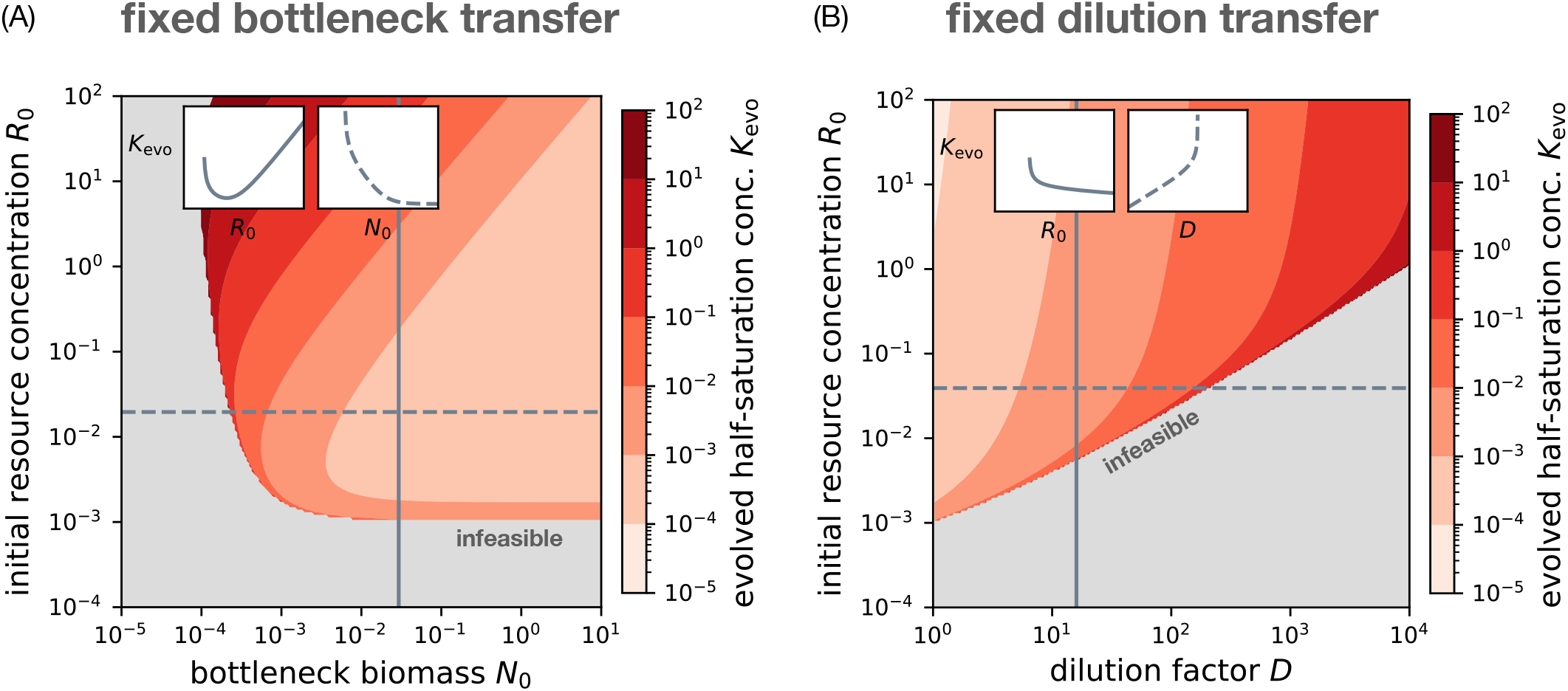
The evolved half-saturation concentration as a function of experimental parameters under coupled genetic drift. (A) Same as Fig. S21A, but where the effective population size *N*_e_ is proportional to the biomass bottleneck *N*_0_ as in well-mixed laboratory experiments with fixed-bottleneck batch dynamics. We set *N*_e_ = *N*_0_*V*, where *V* = 10^5^ is the culture volume such that a bottleneck biomass of *N*_0_ = 0.01 corresponds to an effective population size of *N*_e_ = 10^3^ cells. (B) Same as panel A but for fixed-dilution batch dynamics, where the effective population size is *N*_e_ = *N*_0_*V* = *VR*_0_*Y*/(*D* – 1) (Eq. (S21)).

**FIG. S27.**
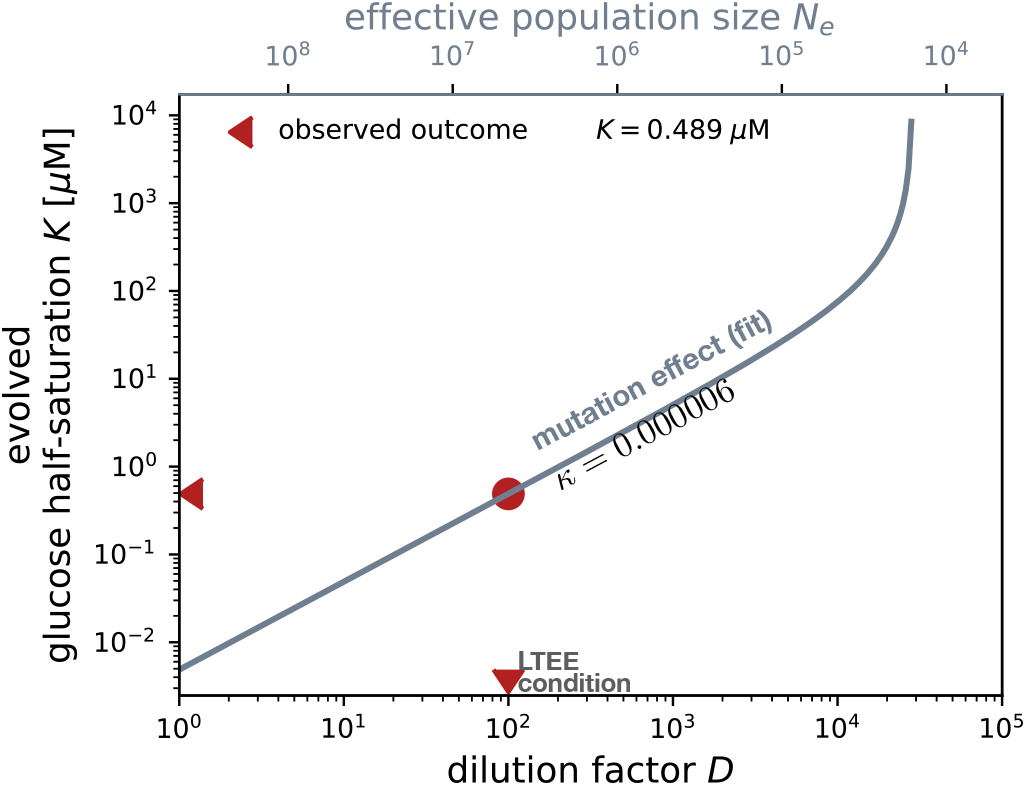
Inferred mutation effect for the Long-Term Evolution Experiment. Evolved half-saturation concentration *K*_evo_ for glucose as a function of the dilution factor *D* under fixed-dilution batch dynamics. If we assume the glucose half-saturation for *E. coli* in the LTEE is under selection-drift balance, then we can use this dependence to infer the value of the mutation effect *κ* that would be consistent with the other known parameters of the system. We numerically solve for selection-drift balance using Eqs. (2) and (3) with dilution factor *D* = 100, initial glucose concentration *R*_0_ = 139 μM, and evolved half-saturation concentration *K* = 0.489 μM (red dot). We obtain an estimate of *κ* = 6 × 10^−6^.

## Supplementary Information

### S1. EFFECT OF COLIMITATION ON ESTIMATES OF MONOD GROWTH TRAITS

The Monod model (Eq. (1)) assumes there is only a single limiting resource whose concentration affects growth rate. However, microbes rely on multiple resources to grow, and therefore their growth rate may depend on the concentrations of all these resources simultaneously. Here we address how these other resources would affect the estimation of Monod growth traits for a focal resource. For simplicity, we consider the case of two essential, independent resources, where resource 1 is the focal resource (e.g., glucose) that we vary over a range of concentrations to measure its Monod parameters 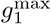 and *K*_1_, and resource 2 is another resource (e.g., ammonium) that is fixed in the background medium. While there is no consensus on the best model for this behavior, we consider three of the most widely-used models:

Liebig model [1–3]:

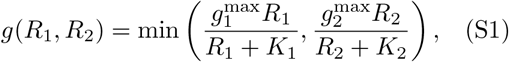

Additive model [1, 4]:

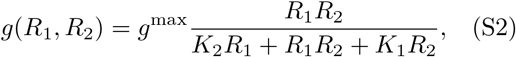

Multiplicative model [2, 4]:

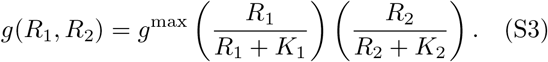

Assuming one of these models is the true description of how growth rate depends on resource concentrations, we imagine fitting an apparent Monod model *g*_app_(*R*_1_) for resource 1 to data generated by the true model, with fixed *R*_2_:

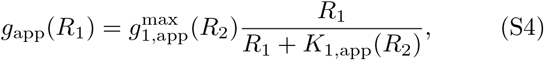

where 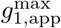 is the apparent maximum growth rate for resource 1 and *K*_1,app_ is its apparent half-saturation concentration, both of which may depend on the concentration *R*_2_ of resource 2. All of the true models correspond exactly to the apparent Monod model — with apparent parameters equaling the true ones, 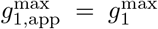 and *K*_1,app_ = *K*_1_ — if the concentration *R*_2_ is much larger than its half-saturation concentration *K*_2_, since the growth rate no longer depends on resource 2 once its concentration is saturating. Therefore *R*_2_ ≫ *K*_2_ is the general condition on the background resource which determines whether we are in the desired regime of limitation only for resource 1.

If the concentration *R*_2_ is smaller or not much larger than its half-saturation concentration *K*_2_, we can then use the models to determine how colimitation with resource 2 affects estimates of Monod parameters for resource 1. For all of the true models, the apparent maximum growth rate 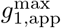 is an underestimate of the true maximum growth rate 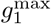. Specifically,

Liebig model:

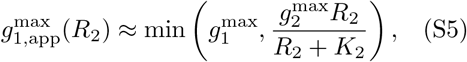

Additive model:

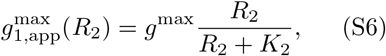

Multiplicative model:

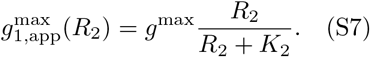

That is, the apparent maximum growth rate for resource 1 is a Monod-type function of resource 2, meaning it is very close to the true 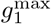 for large *R*_2_ as expected, but becomes a significant underestimate when *R*_2_ is below its half-saturation concentration *K*_2_. Note that the apparent parameters for the Liebig model are only approximate because the Liebig model will not exactly fit the Monod model for a single resource; this is because at some concentration there is a sharp transition in limitation between resources, owing to the minimum function.

The apparent half-saturation *K*_1,app_ is also an underestimate of the true *K*_1_ for the Liebig and additive models, but equals the true value for the multiplicative model:

Liebig model:

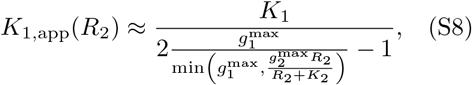

Additive model:

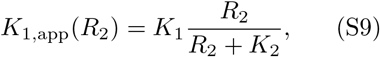

Multiplicative model:

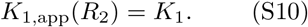

Note also that this means the apparent specific affinity 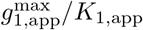 is always correct for the additive model, since the dependence on *R*_2_ cancels out between the apparent maximum growth rate and half-saturation, while it is biased for the Liebig and multiplicative models.

To what extent might these biases affect our data? We can test this condition in a subset of measurements for *E. coli* on glucose where the nitrogen source is ammonium and has a reported concentration. The measured *K* for ammonium in *E. coli* is 2.6 μM (Dataset S1, sheet 1). In the experiments that measure *K* for glucose, the ammonium concentrations are all orders of magnitude higher (0.16 mM to 18.7 mM; Dataset S1, sheet 2). This indicates that ammonium was not colimiting with glucose in these experiments. Indeed, for almost all the resources included in our data, the *K* half-saturation concentrations are much lower than typical laboratory concentrations. This is not surprising in light of our evolutionary model that predicts *K* will often evolve to be much lower than the environmental concentration of the resource (Eq. (4)), and presumably explains why colimitation of essential independent resources has been rarely observed empirically [3].

### S2. ALTERNATIVE MODELS OF GROWTH RATE DEPENDENCE ON RESOURCE CONCENTRATIONS

Table S1 lists several common models for growth rate dependence on resource concentration *R*. Some of these models are mathematically equivalent; for example, Holling [11] proposed a classification scheme for growth models (commonly referred to as Type I, II, and III) for the response of predator growth rate on prey density, which exactly corresponds to other models of growth in Table S1. Some of these models are also equivalent in certain limits. At high resource concentrations *R/K* ≫ 1, all of the models are approximately equivalent to the constant growth model, since the assumption is that resources are saturating and growth is limited by other processes. On the other hand, at low concentrations *R/K* ≪ 1, the Monod, Blackman, and Bertalanffy models are approximately equivalent to the linear model.

There are also some important differences between models. The Blackman, Monod, Bertalanffy, and Hill models all saturate at high resource concentrations, but the nature of that saturation qualitatively differs. That is, the Monod model converges most slowly due its power law dependence on *R*. The Hill model also converges as a power law, but assuming *n* > 1, it does so more quickly than Monod. The Bertalanffy model converges even more rapidly due to its exponential dependence on *R*. Finally, the Blackman model converges to a constant immediately at the half-saturation concentration *R* = *K*.

The model most significantly different from the rest is the Droop model, since it depends not on the external resource concentration directly, but only on the resource concentration internal to the cell. Therefore this requires inclusion of a separate resource uptake process to be included in our framework. Under steady-state (chemostat) growth, this will also be equivalent to the Monod model under a shift in the resource concentration parameter *Q* – *Q*_0_ → *R*, but under non-steady state conditions (e.g., batch dynamics), the Droop model can differ [16].

### S3. MODEL OF BATCH POPULATION DYNAMICS

For batch culture we describe the dynamics of the wild-type and mutant biomasses *N*_wt_(*t*) and *N*_mut_(*t*) and the extracellular resource concentration *R*(*t*) using the following differential equations [20, 21]:

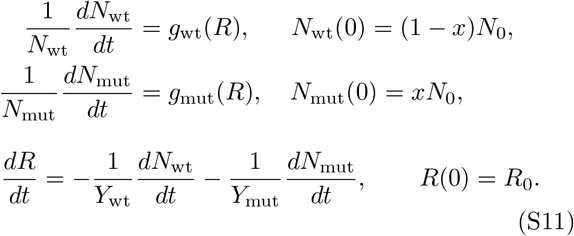

**TABLE S1.**
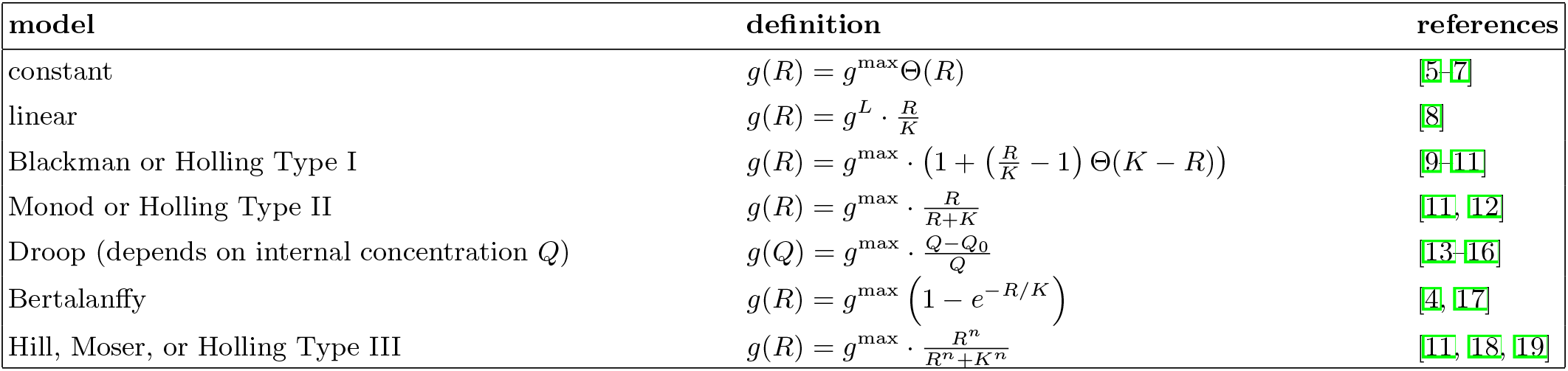
Overview of models for microbial population growth rate. For each entry, the column “references” lists works that establish or build on the model and have been cited elsewhere in this text. The symbol Θ denotes the Heaviside step function which is 1 for a positive argument and zero otherwise.

**TABLE S2.**
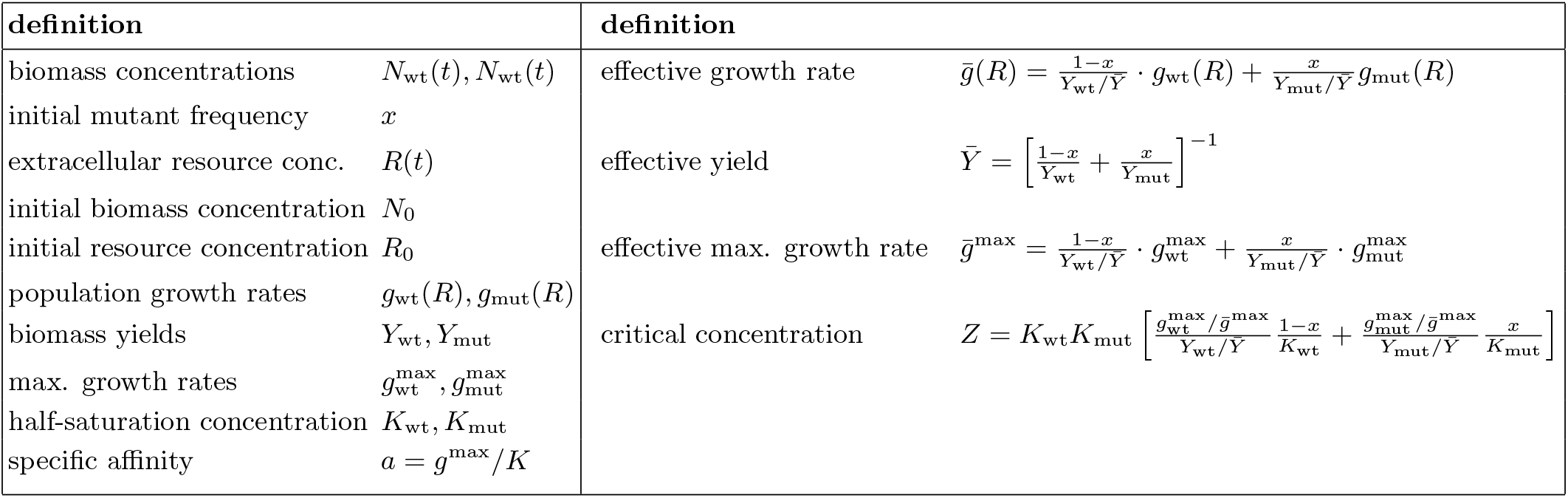
Key notation and definitions used in the model. The subscripts “wt” and “mut” correspond to *wild-type* and *mutant*. Sometimes we drop the subscript “wt” and use a plain letter (*K* or *g*^max^ or a) for the wild-type trait (for example, in the main text).

See Table S2 for a summary of the main notation and definitions used throughout this article. Growth begins with an external resource concentration *R*_0_ and total biomass *N*_0_, a fraction *x* of which is the mutant strain. The strains then grow with per-capita rates *g*_wt_(*R*) and *g*_mut_(*R*), which depend on the extracellular resource concentration *R*(*t*); here we neglect other growth dynamics such as lag [5, 6] and death [22] for simplicity, but they are straightforward to add within this framework. The resource concentration *R*(*t*) declines in proportion to growth of biomass, where the yields *Y*_mut_ and *Y*_wt_ for each strain set the amount of new biomass per unit resource. Here we neglect resource consumption due to maintenance of existing biomass [23], since we expect consumption for maintenance to be much less than consumption for growth during rapid growth. Growth continues until the resource is depleted or the growth rates reach zero. While it is difficult to analytically solve these dynamics in general, it is straightforward to numerically solve the model for a given set of parameters (Sec. S4).

We note that for the Monod model in the limit of low resource concentration *R*, or any model of growth rate that depends approximately linearly on *R* (Table S1), the batch dynamics of Eq. S11 are equivalent to a logistic growth model. We can integrate the equation for resource consumption *dR/dt* in Eq. S11 to express the current resource concentration *R*(*t*) as a function of the biomasses of wild-type *N*_wt_ and mutant strain *N*_mut_:

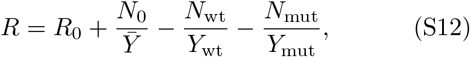

where *Ȳ* is the effective population yield (Table S2). Substituting *R* from Eq. (S12) into the equations for *dN*_wt_/*dt* and *dN*_mut_/*dt* from Eq. S11 with linear growth rate dependence (*g*(*R*) ≈ *g*^max^*R/K*), we obtain

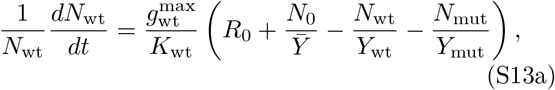

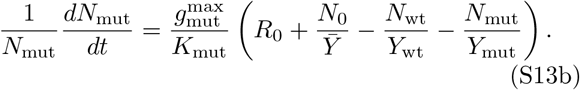

This is equivalent to logistic growth for both species or competitive Lotka-Volterra dynamics.

Once the resource *R* is depleted during a single cycle of batch growth, we transfer a fraction 1/*D* of the population to an environment with a new supply of resources at the original concentration *R*_0_, after which the population resumes growth in the new environment according to Eq. (S11). The factor *D* is known as the dilution factor and is the ratio of the total biomass at the end of the previous growth cycle and the total biomass at the beginning of the next growth cycle [7].

In principle the dilution factor *D* and the bottleneck biomass concentration *N*_0_ can vary over each growth cycle, depending on how we perform the transfers. Let superscript *n* refer to the dynamics during the *n*th batch growth cycle over a series of dilutions and transfers. The biomass at the beginning of the (*n* + 1)th cycle, 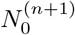, equals the biomass at the end of the previous cycle *n* divided by the dilution factor *D*^(*n*)^ for that cycle:

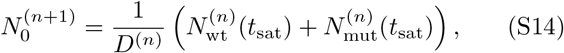

where *t*_sat_ is the saturation time of the growth cycle. To determine the relationship with the bottleneck size of the previous growth cycle, we use the relationship between resource and biomass concentrations (Eq. (S12)) to show that at the end of the growth cycle, *R*(*t*_sat_) = 0, and so *R*^(*n*)^(*t_sat_*) = 0

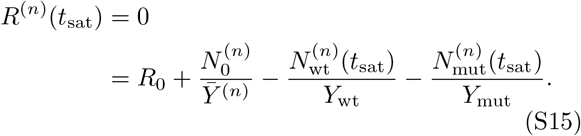

Using this, we can insert the identity to obtain

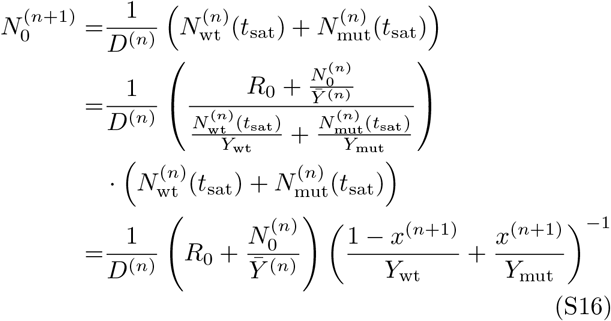

where we have used the fact that the frequencies of each strain at the end of the *n*th cycle equal their frequencies at the beginning of the (*n* + 1)th cycle:

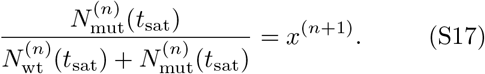

Using the equation for the effective yield (Table S2), we obtain

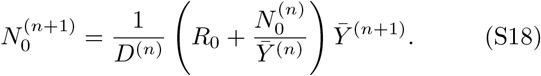

This establishes the general relationship between the bottleneck size and the dilution factor.

Under fixed-bottleneck batch dynamics (Fig. 4B, top panel), 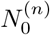 is a constant value *N*_0_, and so we can re-arrange Eq. (S18) to determine how the dilution factor varies at each cycle:

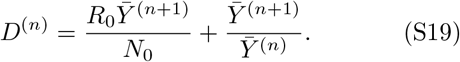

This shows that the dilution factor changes only if the strains have different yields, such that the effective yields *Ȳ*^(*n*)^ change over cycles as the strain frequencies change. On the other hand, under fixed-dilution batch dynamics (Fig. 4B, bottom panel), *D*^(*n*)^ is a constant *D*, and Eq. (S18) simplifies to

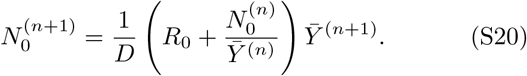

Under both serial transfer regimes, the steady state occurs when *D*^(*n*+1)^ = *D*^(*n*)^, 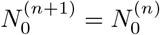, and *Ȳ*^(*n*+1)^ = *Ȳ*^(*n*)^, which implies

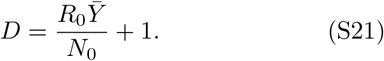

This steady state occurs if 1) all strains have the same yields, such that the effective yield is constant; 2) one strain goes extinct; or 3) the two strains stably coexist.

### S4. NUMERICAL METHODS FOR BATCH DYNAMICS

It is not possible to analytically solve the ordinary differential equations for batch dynamics (Eq. S11). To obtain explicit solutions to this model, we therefore numerically integrate the equations using the Scipy package [24]. We use the default Runge-Kutta algorithm “RK45” in the function `solve_ivp`. This interpolates the differential equation in fourth-order expansion over a short step size *δt*. The step size is automatically adjusted by `solve_ivp` to keep the error of integration below a threshold fixed by the user through the parameters `atol` and `rtol`. Our choices of `atol` = 10^−12^ and rtol = 10^−8^ are more restrictive than the default setting and ensure low errors on the state variables *N*_wt_, *N*_mut_, and *R*.

The population dynamics in Eq. S11 reach the final equilibrium when all resources have been converted into biomass. This final equilibrium is the only attractor since the resources are finite and biomass is strictly increasing (no cell death within a batch growth cycle); in particular, this system does not allow for limit cycles. However, the time to reach this equilibrium is infinite for all growth models in Table S1 (including the Monod model) except for the constant growth rate model. This is because the smooth decline of growth rate prevents full depletion of resources and allows populations to grow indefinitely at infinitesimal but strictly positive growth rates. (The constant growth rate model allows for the same growth rate at arbitrarily low resource concentrations, which means the resources deplete to zero in finite time [5–7].)

For numerical calculations we must therefore set a finite saturation time *t*_sat_ such that the population dynamics are sufficiently close to their equilibrium state. We choose this time using the selection coefficient, which quantifies the relative change in the strain frequencies. Define the cumulative selection coefficient up to time *t* for a batch growth cycle as

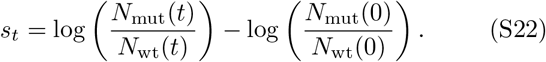

(We further motivate this definition of selection in Sec. S6.) The total selection coefficient for the batch cycle is the cumulative selection coefficient in the limit of infinite time:

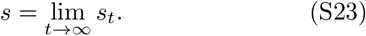

We want to define the saturation time *t*_sat_ as the time where the difference between the cumulative selection up to that time and the total selection is less than some tolerance. We can do this by determining an upper bound on the difference between total selection *s* and the selection at finite time *t*. As the population continues to grow after time *t*, the change in frequencies is bounded by the remaining available resources *R*(*t*). The two possible extremes are if all remaining resources go to the wild-type, in which case the biomass of the wild-type increases by *R*(*t*)*Y*_wt_ and the mutant biomass remains constant, or if all remaining resources go to the mutant, in which case the biomass of the mutant increases by *R*(*t*)*Y*_mut_ and the wild-type remains constant. Therefore the largest possible change in selection occurs in one of these two scenarios, and so the deviation in selection at time *t* from its equilibrium value is bounded by

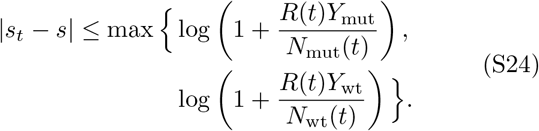

We define the saturation time *t*_sat_ as the shortest time (infimum) such that the difference between the cumulative selection at that time and the total selection is smaller than a given error tolerance *ϵ* > 0:

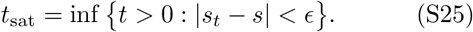

We implement this algorithmically by evaluating the simulation up to an initial time *t*, then evaluating the maximum future error on the selection coefficient from the right hand side of Eq. (S24), and then extending the simulation to *t* + 10 if the error exceeds a defined tolerance *ϵ* = 10^−8^. We iterate this process until the error is less than the threshold.

### S5. MODEL OF CHEMOSTAT POPULATION DYNAMICS

Similar to the batch model of Eq. (S11), the dynamics of biomass and resource concentrations under continuous culture (chemostat) are

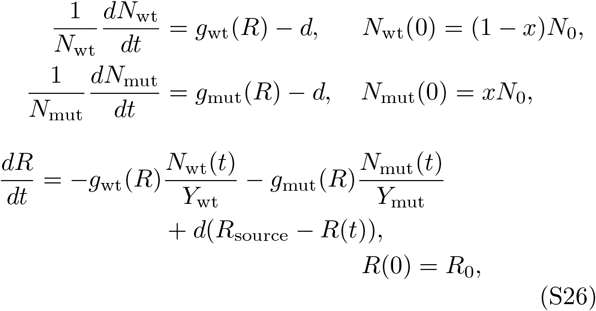

where *R*_source_ is the concentration of the resource in the source media fed into the culture. In a laboratory chemostat, the dilution rate is *d* = *ω*/*V*, where *ω* is the outflow rate (volume per time) and V is the volume of the culture vessel [25].

In the SSWM regime where mutations arise only rarely (Sec. S11), we can assume that the mutant arises on the background of the wild-type at steady-state growth. Let 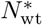 be the steady-state concentration of wild-type biomass and *R*\* be the steady-state concentration of the resource. Note that *R*\* here is the chemostat-specific realization of the ecological concept of a minimum resource concentration required for positive net growth, as used in resource-ratio theory [26, 27]. Since *dN*_wt_/*dt* = 0 in steady state, the resource concentration *R*\* must satisfy

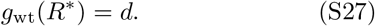

For the Monod model, we can solve this explicitly for *R*\* to obtain

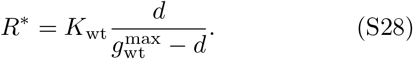

Note that this concentration *R*\* is independent of the source concentration *R*_source_. Using the steady-state condition for the resource *dR/dt* = 0, we can then obtain the steady-state biomass concentration

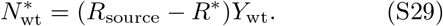

This establishes a feasibility condition for steady state: the dilution rate *d* must be less than the growth rate at the source concentration *g*_wt_(*R*_source_), which is the maximum that the culture can realize for the given resource supply. This criterion has been used by Jannasch [28, 29] to define a *minimum resource threshold* required for population growth at a given dilution factor *d*. This minimum resource threshold corresponds to the steady-state concentration *R*\*, which is related to the parameter *K* but also depends on *d*.

### S6. DEFINITION OF SELECTION COEFFICIENT

The instantaneous selection coefficient *σ*(*t*) measures the rate of change in the logarithm of relative mutant frequency:

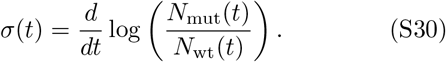

This is a sufficient statistic for frequency change in the sense that knowledge of the instantaneous selection coefficient and the current mutant frequency is sufficient to predict the future mutant frequency over a short time horizon.

For population growth under batch dynamics, the repeated bottlenecks between growth cycles introduce randomness in the frequency trajectory of a mutant. We assume that this stochastic sampling at transfer dominates over the random fluctuations in individual birth rates within the growth cycle. Thus, the genetic drift in our model of serial transfer evolution occurs at the timescale of one growth cycle. To compare the strength of drift and selection on the same timescale, we integrate the instantaneous selection coefficient (Eq. (S30)) over time

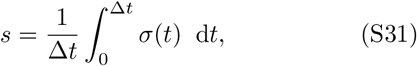

where Δ*t* is the length of the growth cycle. Note that the selection coefficient *s* is still defined as a rate per unit time and in the limit Δ*t* → 0 exactly matches the instantaneous selection coefficient (Eq. (S30)).

For batch dynamics the selection coefficient s determines the change of frequency over multiple growth cycles. At the beginning of the nth cycle, the initial mutant frequency *x*^(*n*)^ is given by

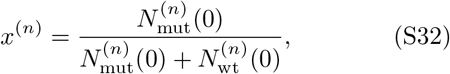

where 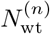 and 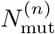 refer to the biomass of wild-type and mutant strains. The population grows to saturation and possibly experiences some frequency change, which sets the mutant frequency *x*^(*n*+1)^ of the next cycle. This change is summarized by the selection coefficient

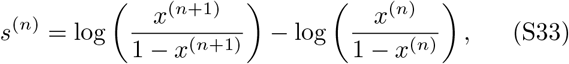

which we compute from the integral definition (Eq. (S31)) using a timescale of Δ*t* = 1 (per growth cycle). Knowledge of *s*^(*n*)^ is sufficient to predict the initial mutant frequency in the next cycle

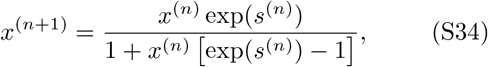

neglecting the stochastic effects of the dilution. Thus, given the starting mutant frequency *x*^(1)^ and the selection coefficients for each growth cycle *s*^(*n*)^, the recursion in Eq. (S34) allows us to predict the mutant frequency trajectory without simulating the population dynamics within each growth cycle.

### S7. DERIVATION OF THE SELECTION COEFFICIENT FOR BATCH DYNAMICS

For populations growing in batch culture, the selection coefficient reduces to the cumulative difference of growth rates:

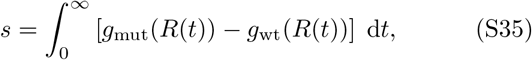

where we have inserted the equations for mutant and wild-type growth from our model of populations dynamics (Eq. S11) into the definition of s from Eq. (S31). The integral extends to infinite time for the growth dynamics to reach equilibrium (Sec. S4) so we therefore do not normalize by the time scale as in Eq. (S31); rather we leave it as understood that the selection coefficient is defined per growth cycle. We can change variables of the integral in Eq. (S35) from time *t* to resource concentration *R*:

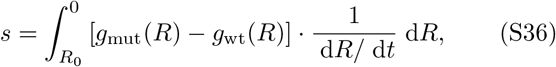

where we have used the fact that *R* ranges from *R*_0_ at the beginning of the growth cycle to 0 at the end of the growth cycle, and that *R* depends monotonically on *t* so that *dt/dR* = (*dR/dt*)^−1^.

To compute this integral, we need to express the transient resource consumption rate *dR/dt* as an explicit function of current resource concentration *R*. As a first step, we rewrite the differential equation for resources (Eq. S11) into the product form

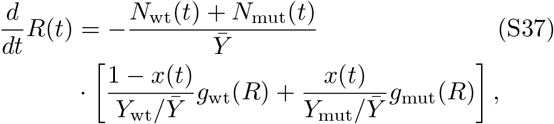

where we use the shorthand *Ȳ* for the effective biomass yield (Table S2). This product separates into the joint biomass *N*_wt_(*t*) + *N*_mut_(*t*) and a new parameter, that we term the *effective growth rate*:

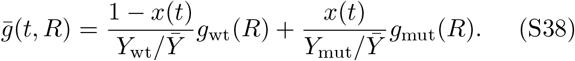

Equation (S37) suggests that this mean of wild-type and mutant growth rates acts as the effective growth rate of the joint population *N*_wt_(*t*) + *N*_mut_(*t*). This effective growth rate is time-dependent due to the underlying frequency change *x*(*t*). Using this equation for the joint biomass (derived from Eq. (S12))

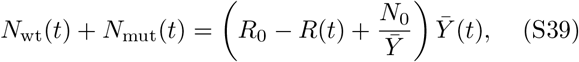

we insert this and the equation for mean growth rate (Eq. (S38)) into Eq. (S36) for the selection coefficient:

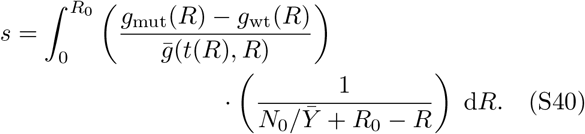

Equation (S40) is an exact expression but requires full knowledge of the resource trajectory *R*(*t*) and its inverse *t*(*R*) to calculate the mean growth rate *ḡ*(*t*(*R*), *R*) in the denominator. For a constant growth rate model (Table S1), this exact expression can be computed [5, 6]. However, for general growth models *g*(*R*) and the Monod model in particular, the integral Eq. (S40) can only be solved under an approximation. Previous work invoked the assumption of small initial mutant frequency *x* ≪ 1 to replace mean growth rate and effective biomass yield by the wild-type traits [20, 21], but here we introduce a novel approximation that holds for all initial mutant frequencies.

We assume that the frequency change over the growth cycle is small, such that the mean growth rate only depends on the resource concentration

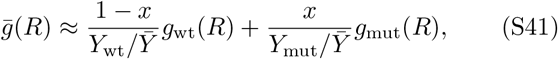

but not otherwise on time *t*. That is, we neglect the time dependence of the mutant frequency *x*(*t*). Thus, we get the explicit integral formula for the selection coefficient:

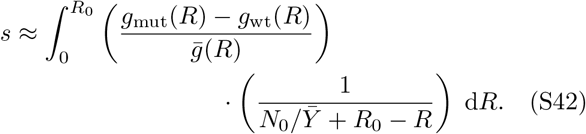

This equation neglects the frequency change *x*(*t*) within the growth cycle but still includes dependence on the initial mutant frequency *x*. One can show that the approximate integral in Eq. (S42) corresponds to a first-order expansion of the exact integral (Eq. (S40) in terms of transient selection coefficients inside the growth cycle, meaning that it is equivalent to a weak-selection approximation. We numerically evaluate the accuracy of this approximation in the case of the Monod model in the next section (Sec. S8).

The exact selection coefficient in its integral form (Eq. (S40)) reveals generic properties of batch-culture competition. First, there is no direct selection for cell yield. A mutant with higher efficiency *Y*_mut_ but equal growth response is neutral. Thus, with an uncorrelated mutation supply, we expect cell yield to evolve neutrally [7, 30]. Second, the selection on the growth rate function *g*(*R*) is distributed unequally across concentrations. In the integrand of Eq. (S40), the difference in growth rates at each resource concentration *R* is weighted by the *fold-change spectrum* 1/(*N*_0_/*Ȳ* + *R*_0_ – *R*). This weight peaks at the initial resource concentration *R*_0_ (see Fig. S13A) and is independent of the growth rate model *g*(*R*). For growth cycles with large fold-change (*R*_0_*Y/N*_0_ ≫ 1), the selection coefficient *s* roughly corresponds to the growth rate difference at initial concentrations because most generations occur at near-constant concentrations close to *R*_0_ (compare Fig. S13B).

A third important property holds only approximately in Eq. (S42), where we see that selection only acts on ratios of growth rates, since the growth rates appear in both the numerator and denominator of the integrand. The dependence on growth rate ratios means that alternative growth models can still lead to equivalent selection on traits. For example, if we take any growth rate model from Table S1 where the mutant and wild-type differ only in their maximum growth rates *g*^max^ (but not other parameters such as *K*), then their selection coefficients will depend only on the ratio 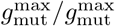 and not other details of the specific model.

### S8. CALCULATION OF THE SELECTION COEFFICIENT FOR THE MONOD MODEL

In this section, we apply the integral formula Eq. (S42) to calculate the selection coefficient for a wild-type and mutant strain competing under the Monod model. Let

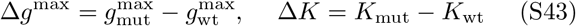

denote the absolute trait differences in maximum growth rate and half-saturation concentration between the two strains. First, we rewrite the relative growth rate difference

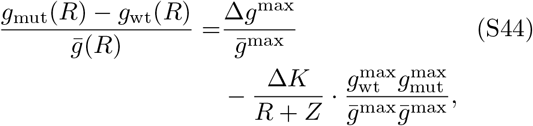

using the effective maximum growth rate

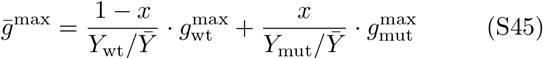

and the critical resource concentration

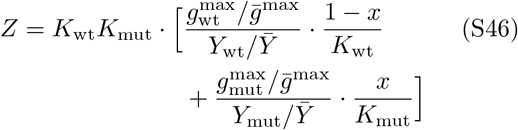

as effective traits of the joint population to simplify the notation (Table S2). Equation (S44) consists of two terms, one proportional to the difference in maximum growth rates Δ*g*^max^ and the other proportional to the difference in half-saturation concentrations Δ*K*. Therefore after substituting this expression into Eq. (S42) and carrying out the integral over *R*, we obtain a selection coefficient consisting of two distinct components:

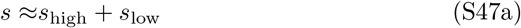

where

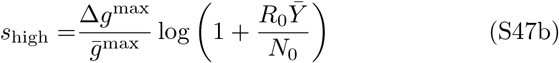

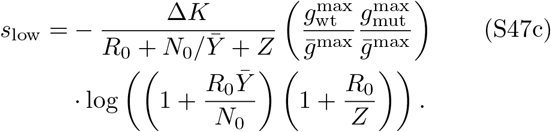

This is the basis for Eq. (2) in the main text under batch dynamics.

The formula for the selection coefficient in Eq. (S42) is based on an approximation of small frequency change. In Fig. S14 we compare the approximate selection coefficient against the exact selection coefficient obtained from numerically solving the differential equations for batch dynamics (Eq. S11). The simulations show that the approximate selection coefficient is accurate up to large values of order *s* ≈ 1. This means that, while we mainly consider the scenario of weak selection (|*s*| < 1), the approximation is excellent even when selection is strong. Intuitively, the approximation should break down because of wrongly estimating the mean resource consumption rate, which we expect to occur when the yields and realized growth rates differ strongly between the two strains. In Fig. S15 we also show a phase diagram of this selection coefficient as a function of the mutant’s traits *g*^max^ and *K* relative to their wild-type values.

The decomposition in Eq. (S47) is useful because the terms correspond to components of selection on distinct phases of growth. The first component, *s*_high_, measures selection on growth at high resource concentrations, and is therefore proportional to the mutational change Δ*g*^max^ in the trait *g*^max^. This mutational effect is weighed by the logarithm of the total fold-change of growth, which equals the dilution factor *D* = *R*_0_*Y/N*_0_ + 1 (Eq. (S21)). An important feature of selection *s*_high_ is that it depends on the nominal maximum growth rate *g*^max^, which is always greater than the realized maximum growth rate *g*(*R*_0_) that actually occurs at the beginning of growth. Therefore the calculation of selection from actual growth data requires an inference of these nominal rates, since the realized rates measured at the beginning of growth curves could produce misleading results if growth begins at low resource concentrations [31].

The second component of selection, *s*_low_, corresponds to growth at low resource concentrations, and is proportional to the mutant’s change Δ*K* of the half-saturation *K*. There is a negative sign in *s*_low_ since selection is positive for mutations that decrease *K* (Δ*K* < 0). For the hypothetical mutant and wild-type in Figs. 1 and 4A,B, *s*_high_ = 0 since the mutant does not change *g*^max^, while *s*_low_ ≈ 0.516, since the mutant has a significantly lower half-saturation concentration *K*. In Fig. S16 we show a more complex pair of strains with a gleaner-opportunist tradeoff (one strain has higher *g*^max^ but also higher *K*), where both components of selection are nonzero [20, 21, 27, 32].

We also briefly discuss an interpretation for the parameter *Z*. The instantaneous selection coefficient (Eq. (S30)) within the batch culture growth cycle

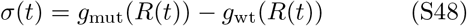

can be decomposed into two components

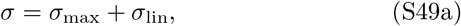

where

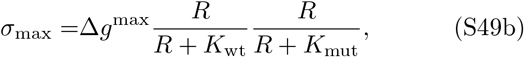

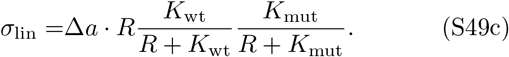

Here *a* = *g*^max^/*K* is the specific affinity, and Δ*a* is the difference in specific affinities between the mutant and the wild-type. The first component *σ*_max_ quantifies growth rate difference at excess conditions, where both strains grow close to their maximum growth rates. The second component *σ*_lin_ measures growth rate differences in the opposite regime, where both strains grow below their half-saturation concentration. The relative size of the two components varies shifts with resource concentration and also depends on the mutation effect on maximum growth rate and specific affinity.

The effective parameter *Z* acts as an intrinsic scale in the resource dependence. Normalizing for different relative mutation effects, both components contribute equally to growth rate difference exactly at external concentration *R* = *Z* such that

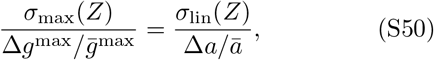

where the effective specific affinity *ā* is defined (in analogy with the effective maximum growth rate defined in Eq. (S45)) as

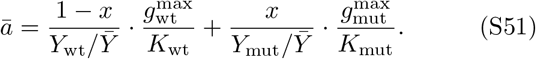

This means, at concentration *Z* both components of the growth rate difference in Eq. (S49) receive equal selection pressure.

The decomposition in Eq. (S49) more generally sugests an alternative parametrization of the Monod model and its selection coefficient. We can replace the half-saturation concentration *K* by the specific affinity *a* = *g*^max^/*K*. This alternative trait corresponds to the growth rate in the limit of low resource concentrations where the Monod model behaves linearly (see Sec. S2). The selecion coefficient in Eq. (S47) can be rewritten as

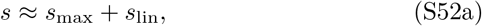

where

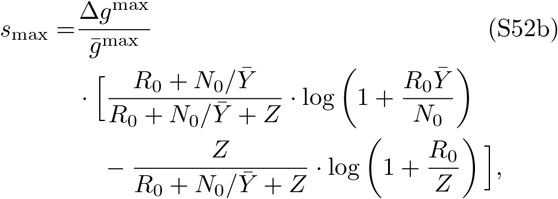

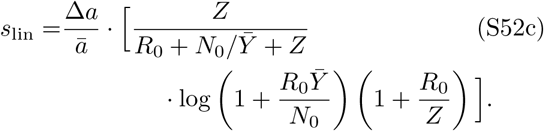

The selection coefficient maps the life-history traits to relative fitness, and the parametrization in *a* is well-suited to study the structure of this map under environmental variation. In the limit of high nutrient concentrations, the total resources are large compared to the critical concentration *Z*. The selection coefficient then reduces to the component of maximum growth:

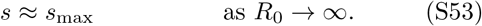

In the opposite limit, the selection coefficient only acts on the growth rate at low concentrations. In this sense, the selection coefficient recovers the limiting behaviour of the underlying growth response:

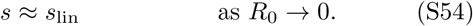

This means that *g*^max^ and *a* are the marginal traits that exclusively control growth in the limiting environments. The selection coefficient reduces to one component or the other. For the parametrization based on *K* given below, this is not true — both *g*^max^ and *K* contribute at low concentrations.

### S9. DERIVATION OF THE SELECTION COEFFICIENT FOR CHEMOSTAT DYNAMICS

For a population in chemostat conditions (Eq. (S26)), the instantaneous selection coefficient *σ*(*t*) (Eq. (S30)) only depends on the difference in growth rates. At a given resource concentration *R*(*t*), this growth rate difference can be decomposed in to two trait components

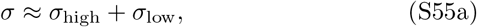

where

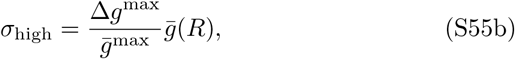

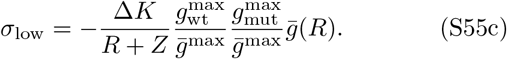

We can derive this by multiplying Eq. (S44) with the mean growth rate for the Monod model

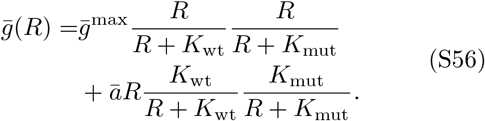

The two components *σ*_high_ and *σ*_low_ are consistent with our results for batch culture conditions (Eq. (S47). By integrating the instantaneous component *σ*_high_ over the growth cycle, we recover the component *s*_high_ for batch-culture growth.

We assume a specific scenario for selection in chemostat populations, where mutants arise at small frequency *x* on top of a wild-type population. This is plausible if mutations occur not too frequently, such that the chemostat population is replaced by a mutant and reaches the new steady state before the next mutation arises. The wild-type population under steady-state chemostat conditions has a resource concentration given by (Eq. S28)

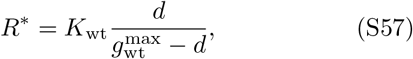

where growth rate matches the dilution factor *g*_wt_(*R*\*) = *d* (Eq. S27). After the mutant appears, the resource concentration *R*(*t*) ≈ *R*\* remains constant over a short timespan while the mutant still has low frequency *x* ≪ 1. In this time window, the mean growth rate Eq. (S56) is set by the wild-type only and thus equals the dilution rate:

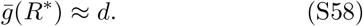

We insert Eq. (S57) and Eq. (S58) into Eq. (S55) to calculate the selection coefficient at invasion with small mutant frequency *x* ≪ 1:

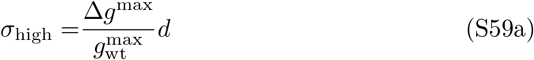

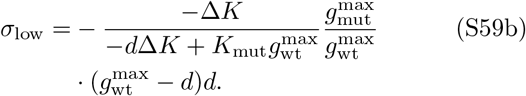

Note that if we express this selection coefficient in terms of the relative mutation effect Δ*K/K*_wt_, then the selection coefficient is independent of the wild-type trait *K*_wt_ (compare to Fig. S20 for batch culture, where the selection coefficient increases with *K*_wt_ for fixed relative mutation effect). This has been observed independently in calculations by Dykhuizen et al. [33], who similarly decompose the growth rate difference in chemostats. As in the case of batch dynamics, the chemostat selection coefficient in Eq. (S59) can also be rewritten in terms of the specific affinity *a* = *g*^max^/*K* instead of the half-saturation concentration *K*.

### S10. DEPENDENCE OF SELECTION ON RESOURCE CONCENTRATION

In this section, we use the explicit formula for s in batch culture (Eq. (S47)) to describe how selection varies with the initial resource concentration *R*_0_ of the growth cycle. For fixed initial biomass *N*_0_, there is an optimum concentration that maximizes selection on the half-saturation concentration *K*. Figure S17A shows nonmonotonic behavior of *s*_low_ with initial resource concentration *R*_0_ for an example mutation with beneficial effects on both the maximum growth rate *g*^max^ and the half-saturation *K*. In particular, this optimum does not rely on a tradeoff between the two traits. Instead, Fig. S17B demonstrates that *s*_low_ is the product of two opposing forces: the overall budget for selection in the growth cycle (equivalent to number of generations) increases with *R*_0_, but the relative selection pressure on the half-saturation concentration decreases. We can identify these two factors from Eq. (S47c) for *s*_low_ on the half-saturation concentration: the selection coefficient is the product of a trait term

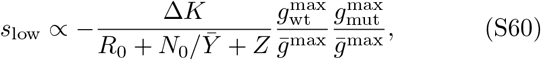

which decreases (in magnitude) with *R*_0_, and a logarithmic term

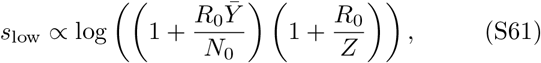

which increases with *R*_0_ via the number of generations in the growth cycle. The optimum concentration, in general, is determined by the wild-type half-saturation concentration (compare Fig. S17C). Figure S17D shows how this causes the distribution of fitness effects to vary in width non-monotonically with the resource concentration as well; the width of this distribution is generally proportional to the speed of adaptation [34], which thus also displays a local maximum and minimum over resource concentrations.

These effects are not observed in batch dynamics with fixed-dilution factor, where selection *s*_low_ decreases strictly monotonically with resource concentration. The same example mutation in Fig. S18 reaches peak selection at the lowest nutrient concentration *R*_0_. Intuitively, the fixed dilution factor *D* means the total budget for selection (number of generations) is independent of the initial concentration *R*_0_ and low concentrations mean a larger fraction of time spent in deceleration, but not fewer generations.

### S11. MODEL OF EVOLUTIONARY DYNAMICS UNDER STRONG-SELECTION WEAK-MUTATION

We can map the dynamics of the mutant frequency over batch growth cycles to the Wright-Fisher model of population genetics, where each batch growth cycle corresponds to a discrete time step [5, 35]. First, we assume the mutation arises only at the beginning of the growth cycle at frequency 1/*N*_0_, where *N*_0_ is the bottleneck population size measured in number of cells. Let *s*(*x*) be the selection coefficient for the mutant over a whole batch growth cycle, with explicit dependence on the frequency *x* of the mutant at the beginning of the cycle. In the limit of large population size (*N*_0_ ≫ 1) and weak selection (|*s*(*x*)| ≪ 1), the fixation probability for the mutant is [36]

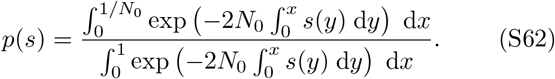

However, if the selection coefficient *s*(*x*) is approximately constant over mutant frequencies *x*, we can simplify this to

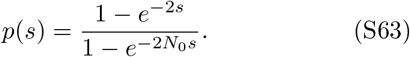

We briefly describe the scheme for simulating trait evolution. In general, a mutation can change both growth traits

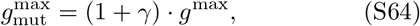

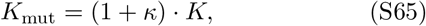

where *γ* is the mutation effect on the wild-type maximum growth rate *g*^max^ and *κ* is the relative effect on the half-saturation concentration *K*. Given the absence of correlation between *g*^max^ and *K* for autotrophs on phosphate, nitrate and ammonium (Figs. 3C–D, S10E) and for heterotrophs on glucose (Figs. 3F, S10G), we assume that mutations affect *K* independently of maximum growth rate (*γ* = 0). We simulate evolutionary trajectories of the half-saturation concentration *K* by first randomly sampling a mutation effect *κ* from a uniform distribution on the interval (−0.1, 0.1). We then calculate the selection coefficient of this mutation using Eq. (S47) and the fixation probability according to Eq. (S63). We randomly accept or reject the mutation according to this probability, and then the cycle repeats with a new mutation (Fig. S19). We also test the effect of frequency-dependence selection using the fixation probability of Eq. (S62), but Fig. S22D-F shows that it does not noticeably affect evolution of the half-saturation concentration.

### S12. DERIVATION OF SELECTION-DRIFT BALANCE CONDITION

In the limit of weak selection (*s* ≪ 1), we can expand Eq. (S63) to leading order in *s*:

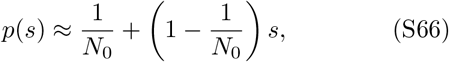

where the first term captures the probability of fixation due purely to demographic fluctuations (genetic drift), while the second term captures the correction due to selection. The balance between selection and drift therefore occurs when these two contributions are approximately equal, which gives us *s* ≈ 1/*N*_0_ (Eq. (3) from the main text) under the additional assumption that *N*_0_ is large.

Now we consider the effect of a mutation arising at some intermediate time *t* during a growth cycle. Since at this time there are *N*_wt_(*t*) wild-type cells, the initial frequency of the mutant is 1/*N*_wt_(*t*), and the amount of remaining resources is *R*(*t*) = *R*_0_ – (*N*_wt_(*t*) – *N*_wt_(0))/*Y*_wt_. Therefore the frequency of the mutant at the end of this cycle is

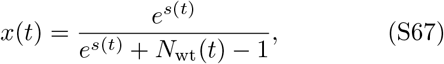

where *s*(*t*) is the selection coefficient for this mutant arising at time *t*, assuming a growth cycle that starts when the mutation arises (so we use *R*(*t*) as the initial amount of resources and *N*_wt_(*t*) as the initial population size).

Let *p*(*t*) be the probability that this mutant ultimately fixes. This is the probability that *n* mutant cells survive the transfer, multiplied by the probability those mutants fix, averaged over all possible *n*:

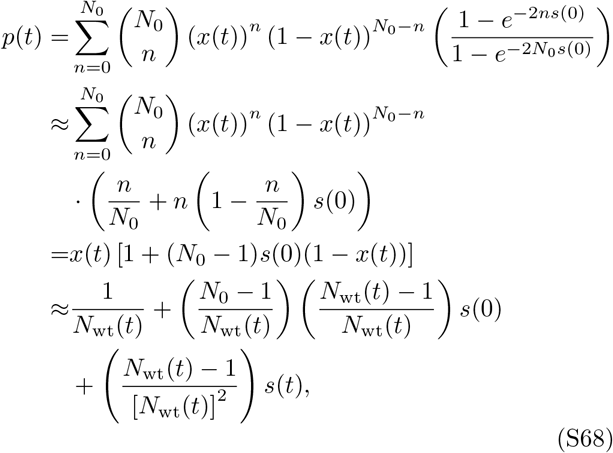

where we have invoked the weak-selection approximation to the fixation probability (Eq. (S66)) on the second line, evaluated moments of the binomial distribution on the third line, and then expanded the frequency *x*(*t*) (Eq. (S67)) to leading order in *s*(*t*) on the last line. By neglecting terms that are higher-order in 1/*N*_wt_(*t*) and *s*(*t*), we obtain

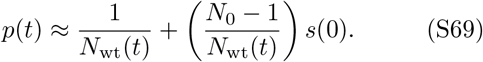

Note that this only depends on the selection coefficient of the mutant starting at the beginning of the cycle; to leading order there is no dependence on the selection coefficient during that first cycle *s*(*t*). If we calculate the condition for selection-drift balance as before, we obtain *s*(0) ≈ 1/*N*_0_ as before. That is, the dependence on the wild-type population size at which the mutant first arises *N*_wt_(*t*) is irrelevant to the selection-drift balance. Therefore mutations arising during growth cycles have no effect on the selection-drift balance condition to leading order.

### S13. THE EVOLVED HALF-SATURATION CONCENTRATION AT SELECTION-DRIFT BALANCE

In this section, we calculate the evolved half-saturation concentration *K*_evo_ as a function of environmental con-centration *R*_0_ and effective population size *N*_e_. We assume mutations have a maximum relative effect |*κ*_max_| = |Δ*K/K*_wt_| on the half-saturation concentration, but no effect on maximum growth rate or biomass yield. Therefore the maximum possible selection coefficient for any mutant on the background of a wild-type trait *K* is thus

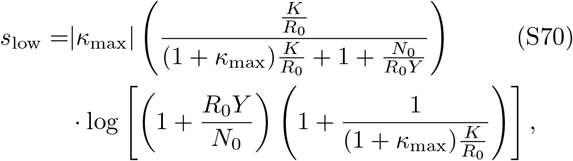

where we have rewritten the selection coefficient (Eq. (S47)) in terms of the ratio *K/R*_0_ between the wild-type half-saturation concentration and the initial resource concentration. Note that we write Y for the wildtype biomass yield, which remains unchanged throughout evolution.

To simplify Eq. (S70), we assume that the maximum mutation effect is small (|*κ*_max_| ≪ 1), the value of the half-saturation concentration *K* relative the initial resource concentration is small (*K/R*_0_ ≪ 1), and the fold-change over the growth cycle is large (*R*_0_*Y/N*_0_ ≫ 1). This is true for growth cycles in typical laboratory evolution experiments, with typical dilution factors between *D* = 100 [37] and *D* = 1500 [38]. We therefore approximate the selection coefficient in Eq. (S70) by keeping only leading-order terms in these parameters:

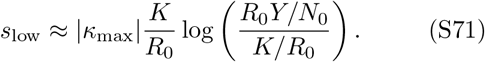

The evolved half-saturation concentration *K*_evo_ is defined as the value of the half-saturation *K* such that the selection coefficient for a mutation on this half-saturation equals the fixation probability of a neutral mutation. We must therefore also assume that the maximum strength of selection, which occurs for large *K*, is greater than the neutral fixation probability (Fig. S20A). In the limit of small |*κ*_max_| and large *R*_0_*Y/N*_0_, the maximum selection coefficient is |*κ*_max_| log(*R*_0_*Y/N*_0_), and so this must be greater than 1/*N*_e_. To solve for *K*_evo_, we then set the selection coefficient in Eq. (S71) equal to 1/*N*_e_ (using Eq. (3)) and solve to obtain

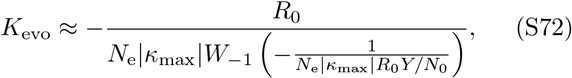

where *W*_−1_(*z*) is the −1 branch of the Lambert *W* function, defined as the solution of the equation *ye^y^* = *z* for −*e*^−1^ ≤ *z* < 0 [39]. The latter condition is met since the argument of the *W* function, −1/(*N*_e_|*κ*_max_|*R*_0_*Y/N*_0_) is certainly less than zero, but also

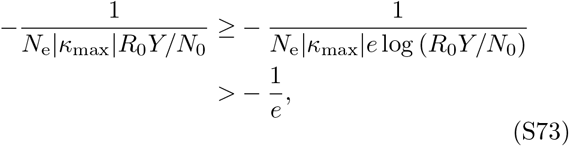

where on the first line we have used the fact that *e* log(*R*_0_*Y/N*_0_) ≤ *R*_0_*Y/N*_0_ and on the second line we have used *N*_e_|*κ*_max_| log(*R*_0_*Y/N*_0_) > 1 from our previous assumption that the maximum strength of selection is bigger than genetic drift. We can further simplify Eq. (S72) using the approximation *W*_−1_(*z*) ≈ log(−*z*) for |*z*| ≪ 1, which gives us Eq. (4) in the main text.

We note that this calculation does not work for the chemostat selection coefficient (Eq. (S59)) since it does not depend on the wild-type trait *K*_wt_ outside of the relative mutation effect Δ*K*/*K*_wt_. Therefore the selection coefficient does not decrease as *K* evolves lower, and there is no selection-drift balance.

### S14. EVOLUTION TO SELECTION-DRIFT BALANCE FOR THE SPECIFIC AFFINITY

In this section we repeat our evolutionary analysis using the specific affinity *a* = *g*^max^/*K*, instead of the half-saturation concentration *K*, as the focal trait for mutation and selection. First we simulate evolution in the SSWM regime, then we predict the evolved trait from a selection-drift balance condition and derive a scaling relationship with resource concentration *R*_0_ and effective population size Ne. In combination with the maximum growth rate *g*^max^, the specific affinity *a* gives an alternative parametrization of the Monod model of growth. Equation (S52) decomposes the total selection coefficient s in batch culture, where the component *s*_lin_ captures the trait differences in the specific affinity *a* = *g*^max^/*K*.

We assume mutations have a relative effect *α* on the specific affinity

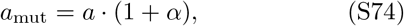

but leave the maximum growth rate *g*^max^ and biomass yield *Y* unchanged. The effect size *α* is sampled at random from a uniform distribution, with maximum value *α*_max_ > 0. This means a single mutation can increase the specific affinity at most by a fixed fraction *α*_max_. This set of assumptions mirrors the evolutionary simulations carried out for the half-saturation *K*. We simulate the trait evolution over long times, where each new mutation either fixes or goes extinct before the next mutation arises.

Figure S23 shows that evolution of the specific affinity *a* = *g*^max^/*K* leads to behavior that is analogous to when mutations target the half-saturation concentration *K*: the specific affinity *a* evolves upwards over successive mutations, improving the growth rate at low concentration, but eventually the trait *a* stalls in adaptation around an upper limit. The limiting value depends on the effective population size *N*_e_ between transfers (compare panels in Fig. S23). Following the same reasoning as in Sec. S13, we define the evolved trait *a*_evo_ as the trait value where selection-drift balance is achieved:

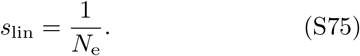

Figure S23 shows that the simulated trajectories are predicted well by Eq. (S75), which we solve numerically for specific affinity *a*_evo_ at selection-drift balance.

We follow the same steps as in Sec. S13 to derive a similar scaling relationship for *a*_evo_ as a function of the resource concentration *R*_0_ and the effective population size *N*_e_. The maximum possible selection coefficient for any mutation on the background of a wild-type trait *a* is

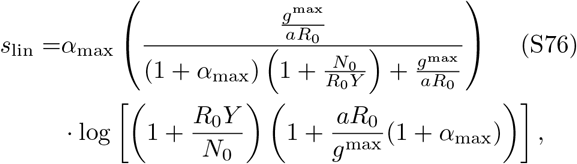

where we have rewritten the selection component (Eq. (S52c)) in terms of the ratio *g*^max^/(*aR*_0_) = *K/R*_0_ between the wild-type traits and the initial resource concentration. To simplify Eq. (S76), we assume that the maximum mutation effect is small (*α*_max_ ≪ 1), the foldchange over the growth cycle is large (*R*_0_*Y/N*_0_ ≫ 1), and the evolved value of the specific affinity *a* is large relative to the initial resource concentration (*g*^max^/(*aR*_0_) ≪ 1). This last assumption is equivalent to assuming a highly-adapted half-saturation concentration (*K/R*_0_ ≪ 1), just as we did in Sec. S13. We thus approximate the selection coefficient in Eq. (S76) by keeping only the leading-order terms in these parameters:

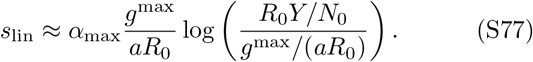

The evolved specific affinity *a*_evo_ is defined as the value of the specific affinity such that the selection coefficient for a mutation on this trait value equals the fixation probability of a neutral mutation. Again, we must assume that the maximum strength of selection, which occurs for small *a*, is greater than the neutral fixation probability (Fig. S20B). In the limit of small *α*_max_ and large *R*_0_*Y/N*_0_, the maximum selection coefficient is *α*_max_log(*R*_0_*Y/N*_0_) so this must be greater than 1/*N*_e_. To calculate *a*_evo_, we then set the selection coefficient in Eq. (S77) equal to 1/*N*_e_ and solve to obtain

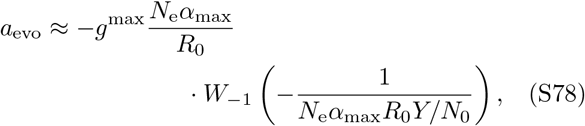

where *W*_−1_(*z*) is the −1 branch of the Lambert W function, introduced above in Eq. (S72). Just as before, we confirm that the evolved trait *a*_evo_ is confined to this solution branch and use the approximation *W*_−1_(*z*) ≈ log(−*z*) to arrive at the final scaling relationship

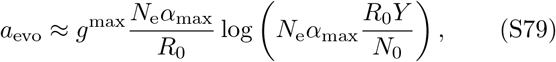

which is the analogous result to Eq. (4) in the main text.

How does the evolved specific affinity *a*_evo_ (Eq. (S78)) compare to the evolved half-saturation concentration *K*_evo_ (Eq. (S72))? They are mathematically equivalent if the mutation effects sizes *α*_max_ and |*κ*_max_| are equal, which holds in the limit where they are both small. That is, if we express the relation *a*_mut_ = *a*(1 + *α*_max_) for the mutation effect on *a* as 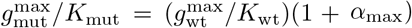, and then use the fact that *g*^max^ is unchanged by the mutation 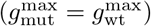, we then get

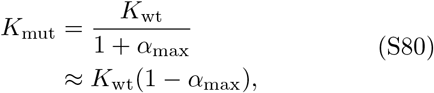

which, compared with the definition of *κ* = (*K*_mut_ – *K*_wt_)/*K*_wt_, shows that *α*_max_ = |*κ*_max_| when both are small.

Altogether this shows that focusing on specific affinity *a* leads to equivalent evolutionary outcomes as focusing on the half-saturation concentration *K*, including the dependence on the resource concentration *R*_0_ and the mode of population dynamics (fixed-bottleneck or fixed-dilution batch dynamics, or chemostat dynamics). This makes sense since mutations that affect *a* but leave *g*^max^ constant must therefore only affect *K*, and thus the only difference between these approaches is the choice of mathematical parameterization. We can also speculate what would happen if mutations affect both the maximum growth rate *g*^max^ and the specific affinity *a* simultaneously (but assuming no correlation in effects). We expect that the maximum growth rate will evolve to the highest physiologically-feasible value, which will serve as the effective maximum growth rate to convert between *a* and *K*. Intuitively, this would still lead to identical selection-drift balance for the half-saturation concentration *K* and the specific affinity *a*.

### S15. EFFECT OF EVOLVED HALF-SATURATION CONCENTRATION VALUES ON MEASUREMENT APPROACHES

In the main text we present a survey of empirical values for the half-saturation concentration *K*, as well as an evolutionary model suggesting that *K* should generally be much smaller than the concentration of the corresponding resource in the evolutionary environment. Here we explore what these values of *K* mean for three approaches to measuring *K* under laboratory conditions.

## A. Inferring half-saturation concentrations under chemostat growth

Arguably the most direct approach to measuring *K* is to use a chemostat (Sec. S5). This setup takes an inverse approach to the Monod model relation in Eq. (1): instead of varying the resource concentration *R* and measuring the growth rate *g*, as suggested by the functional form of the model, we vary the growth rate (by controlling the dilution rate *d*, which must equal the growth rate *g* in steady state) and measure the corresponding resource concentration *R*. We first identify the maximum growth rate *g*^max^ by gradually increasing the dilution rate *d* until the population collapses; the maximum dilution rate that the population can sustain equals the maximum growth rate *g*^max^. Then we set the dilution rate to half the maximum growth rate (*d* = *g*^max^/2) and measure the resource concentration at this state, which by definition of the Monod model (Eq. (1)) must equal the half-saturation concentration *K*.

In light of what we know about typical values of the half-saturation concentration *K*, what challenges does this pose for such measurements? We must either directly measure resource concentrations in the medium around the value *K* (which may be difficult depending on the sensitivity of such a measurement), or infer the resource concentration from the biomass concentration *N*\* = (*R*_source_ – *K*)*Y* (Eq. (S29)) In the latter case, we would also need to know the source concentration *R*_source_ we are supplying to the culture as well as the yield *Y*. However, we are not limited by low biomass concentrations in the chemostat, as we can arbitrarily increase the biomass concentration by increasing the source concentration *R*_source_. For example, for *E. coli* on glucose, the half-saturation concentration is *K* ~ 10 μM (Fig. 2B), the yield is *Y* = 3.3 × 10^8^ cells/μmol [30], and a typical laboratory concentration of glucose to provide could be *R*_source_ = 11000 μM (0.2% w/v). In this case the concentration of *E. coli* would be 3.6 × 10^9^ cells/mL, which is high enough to easily measure through different standard techniques. For example, this cell density corresponds to an optical density (OD) of approximately 3.6 (using 1 OD = 10^9^ cells/mL, for wavelengths of 600 nm and a path length of 1 cm), which is easily measured in a standard spectrophotometer.

## B. Inferring half-saturation concentrations under batch growth using the initial growth rate

A second approach uses cultures under batch growth. This takes a direct approach to the Monod model compared to the chemostat: we vary the initial concentration of the resource over some range around the concentration *K* and measure the initial growth rate of the biomass as a function of these concentrations. We then fit this data to the Monod model (Eq. (1)) and infer the concentration *K*. Note that this assumes that the population can rapidly adjust its growth rate to the external resource concentration, so that the measurement is not biased by the previous state of the culture (e.g., under starvation).

Therefore we need to perform this experiment with initial resource concentrations *R*_0_ that are around the value of *K*. The total biomass concentration at the end of such a batch growth cycle would be *KY* + *N*_0_, where *N*_0_ is the initial biomass concentration. Using the previous example of *E. coli* on glucose, the biomass concentration *KY* is approximately 3.3 × 10^6^ cells/mL, which corresponds to an OD of 3.3 × 10^−3^. However, to measure growth, we must start at a concentration at least 10–100 times lower than this to have a sufficiently large dynamic range of the biomass to accurately measure the growth rate. This range of concentrations is too low to be detected on typical spectrophotometers, which usually have a lower limit of 10^−3^ to 10^−2^ OD, so only methods with greater sensitivity to low concentrations (e.g., colony counting on plates or luminescence) would be suitable. In this case, note that the difficulty with measuring *K* this way is not due to its magnitude relative to a typical glucose concentration *R*, but that the biomass produced by this resource concentration (*KY*) is low compared to the lower limit of typical detection methods.

## C. Inferring half-saturation concentrations under batch dynamics using the deceleration into starvation

The third approach also uses batch cultures, but instead of considering how the initial growth rate varies with initial resource concentration, we use a fixed initial resource concentration *R*_0_ and infer *K* from how growth rate spontaneously decelerates into starvation at the end of the growth cycle. Equation S11 defines the ODEs for batch growth with a wild-type and mutant strain. If we simplify this to a single strain, insert the Monod model for growth rate (Eq. (1)), and integrate the resource consumption equation (to express resource *R*(*t*) in terms of biomass *N*(*t*), as in Eq. (S12)), we obtain a single non-linear ODE for the biomass concentration:

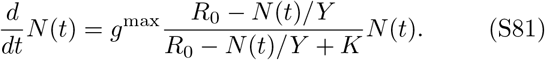

In principle we can fit this ODE to time-series data for the biomass concentration *N*(*t*) (the growth curve) and infer the half-saturation concentration *K*.

Intuitively, though, this only works if the growth curve has enough data during the deceleration phase of growth where the half-saturation *K* is relevant; see Fig. S24 for a schematic example. Previous work has studied this as a problem of statistical estimation, calculating parameter sensitivities to identify the optimum measurement concentration and discussing variable transformations to simplify the regression (see Robinson [40] for an overview). The basic conclusion is that the initial resource concentration *R*_0_ must be near the value of the half-saturation concentration *K* itself for the fit to work robustly.

We can justify the intuition for this conclusion as follows. If the initial resource concentration *R*_0_ is instead much greater than the half-saturation concentration *K*, then the fold-change during deceleration will be too small to provide sufficient dynamic range for a fit. That is, deceleration approximately begins at the time *t*_decel_ when *R*(*t*_decel_) = *K*, so that the biomass concentration is *N*(*t*_decel_) = *N*_0_ + (*R*_0_ – *K*)*Y*. Since the final biomass concentration at saturation is *N*(*t*_sat_) = *N*_0_ + *R*_0_*Y*, the fold-change during deceleration is therefore

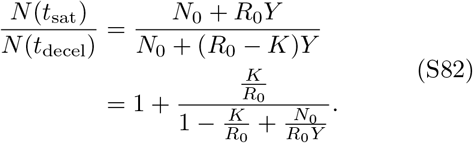

However, if *R*_0_ is much larger than *K*, then this fold-change is approximately 1 + *K*/*R*_0_, meaning that it is very close to 1 (corresponding to no growth during deceleration). Visually, this appears as a growth curve with an abrupt transition from the maximum growth rate *g*^max^ to zero growth (inset of Fig. 4). Since typical concentrations of many resources (such as glucose) used in the laboratory are indeed much larger than the *K* half-saturation concentrations, this is why these growth curves usually do not contain useful data on the half-saturations *K*.

On the other hand, if *R*_0_ is much less than *K*, then the growth dynamics are approximately logistic:

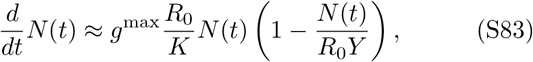

which we obtain similarly with Eq. (S81) but in the limit *R*_0_ ≪ *K*. In this case, one can only infer the combined parameter *g*^max^*R*_0_/*K* from the growth curve and not the half-saturation concentration *K* by itself. Therefore the half-saturation *K* can only be inferred from the growth curve if the initial concentration *R*_0_ is around the value of *K* itself. However, this is the same parameter regime as needed for the previous method of inferring *K* from the initial growth rates, and thus it poses the same practical challenges, such as sensitivity to very low biomass concentrations.

## Notes

### Competing Interest Statement

The authors have declared no competing interest.

### Summary of Updates

added new Figure S8 on temperature dependence; adjusted wording to use 'half-saturation concentration' for the Monod parameter K

